# Genomic Landscapes of Natural Selection in Great Apes

**DOI:** 10.1101/2025.08.29.673040

**Authors:** Xin Huang, Simon Chen, Sojung Han, Martin Kuhlwilm

**Affiliations:** Department of Evolutionary Anthropology, University of Vienna, Vienna, Austria.; Human Evolution and Archaeological Sciences (HEAS), University of Vienna, Vienna, Austria.

**Keywords:** Positive selection, Balancing selection, Distribution of fitness effects, Great apes, Population genetics

## Abstract

**Background:** Great apes are a primate clade with unique biological features. The different species and subspecies have adapted to their respective habitats and followed various evolutionary trajectories influencing their diversity. Hence, studying patterns of natural selection across this clade is an important aspect in understanding their biology.

**Results:** We analyzed a curated panel of genomic diversity for bonobos, chimpanzees, eastern and western gorillas, and Bornean and Sumatran orangutans to explore their selection landscapes. A genome-wide screen was conducted for recent positive and long-term balancing selection, revealing candidate loci potentially related to sensory and immune systems, environmental pressures such as diet or altitude, and reproductive strategies in these species. In addition, the distribution of fitness effects (DFE) was determined, along with the correlations of DFEs across lineages.

**Conclusions:** We interpret the potential biological meaning of the landscapes of recent positive and long-term balancing selection in our closest living relatives, revealing novel lineage-specific candidate genes and gene categories, as well as recurrent targets across lineages. Our investigation of the deleterious DFE indicates that most non-synonymous mutations fall into either the nearly neutral or strongly deleterious categories, with high correlations between closely related lineages.

## Background

The genomic basis of primate diversity and adaptation is an important field of evolutionary biology [1]. Great apes, our closest living relatives, exhibit remarkable diversity in their morphology, physiology, and behavior. Understanding how these differences are shaped by evolutionary forces, particularly natural selection, is directly relevant to our own biology.

Previously, a landmark study, the Great Ape Genome Project (GAGP), characterized genomic variation across great apes and provided a dataset that has served as the foundation for many subsequent studies [2]. Building on the GAGP, different methodologies have been applied to generate a more comprehensive picture of how natural selection shaped great ape diversity. For example, neutrality tests have been used to identify both balancing and positive selection at different timescales [3], and it has been proposed that the impact of selection on diversity increases with the effective population size [4]. Later, ancient selection on protein-coding genes has been investigated [5], while a recent study on telomere-to-telomere genome assemblies [6] leveraged a larger panel of great ape genomes [7–9]. Further studies have contributed to a more comprehensive picture of great ape diversity, which was recently compiled into a curated genome panel [10].

Within great apes, the *Pan* genus (chimpanzees and bonobos) is the most closely related to humans (Fig. 1), resulting in specific interest in these species. Four sub-species of chimpanzees (*Pan troglodytes*) have been described [2], while bonobos (*Pan paniscus*) show population substructure [11]. Bonobos are found exclusively in the Congo basin in central Africa, with unique sociosexual and physiological characteristics. These have been linked to genomic changes in genes related to female reproduction, and sensory genes such as taste receptors [6, 12]. The chimpanzee sub-species with divergence times up to 600 thousand years ago (kya) [7] between lineages have been investigated for population-specific signatures of selection, with only few overlapping genes between subspecies [3, 13]. Selection on spermatogenesis and the immune system as well as diet-related adaptation has been reported [3, 6, 13–15]. Particularly strong evidence from exposure to the Simian Immunodeficiency Virus in eastern and central chimpanzees has been provided [16, 17]. Finally, patterns of local adaptation to both forest and savannah habitats with association with pathogen response have been reported [18].

**Fig. 1:**
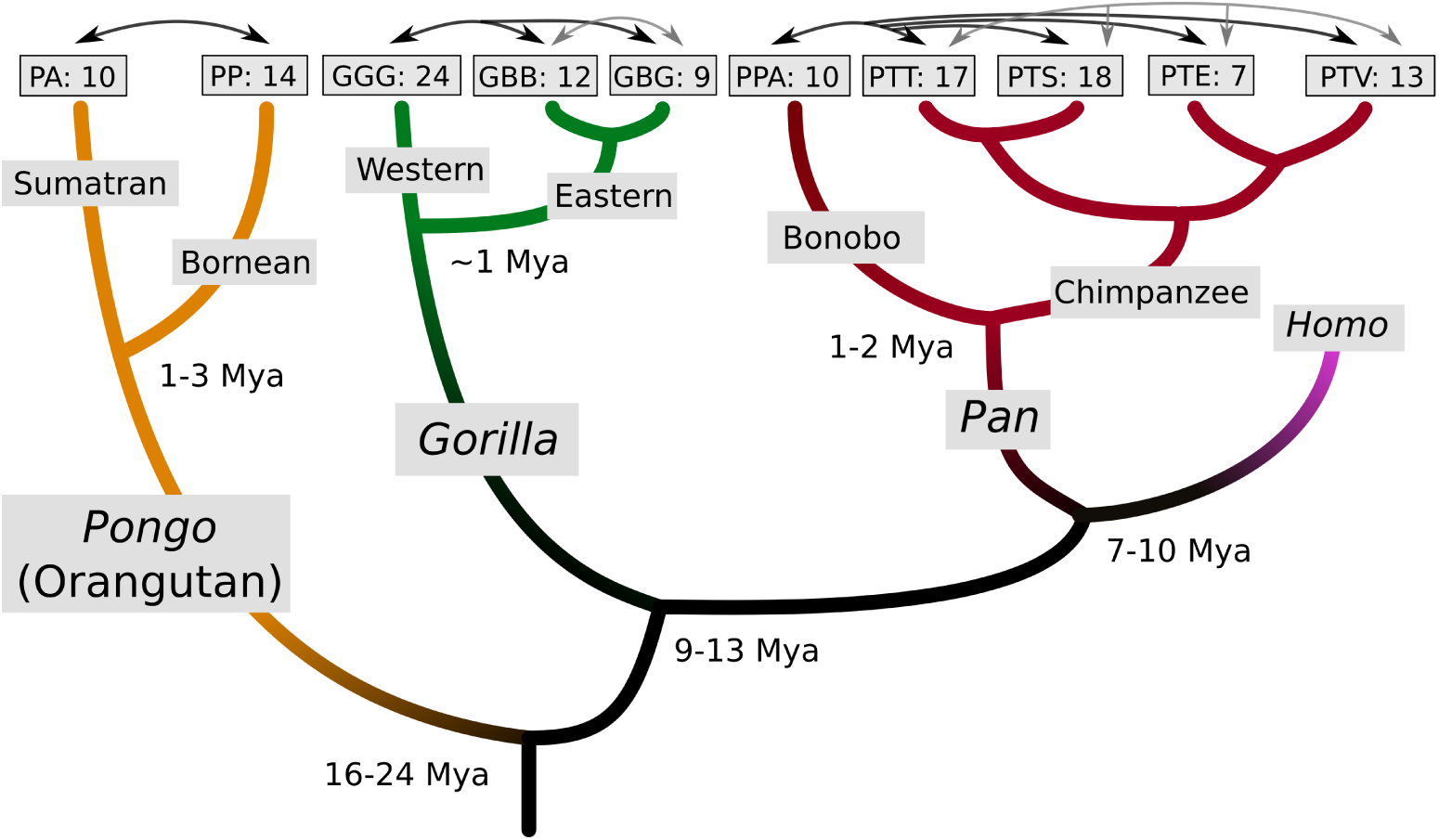
Phylogeny of the great ape species and subspecies considered in this study. Orangutans diverged from African great apes, with two species included here: *Pongo abelii* (PA) and *Pongo pygmaeus* (PP). For gorillas, two species have been described: *Gorilla gorilla*, with the subspecies western lowland gorilla (*Gorilla gorilla gorilla*, GGG) included here, and *Gorilla beringei* (eastern gorilla) with two subspecies: mountain gorilla (*Gorilla beringei beringei*, GBB) and eastern lowland gorilla (*Gorilla beringei graueri*, GBG). The genus *Pan* includes two species: bonobos (*Pan paniscus*, PPA) and chimpanzees (*Pan troglodytes*). For chimpanzees, the four known subspecies are included: central chimpanzee (*Pan troglodytes troglodytes*, PTT), eastern chimpanzee (*Pan troglodytes schweinfurthii,* PTS), Nigeria-Cameroon chimpanzee (*Pan troglodytes ellioti,* PTE), western chimpanzee (*Pan troglodytes verus*, PTV). Approximate divergence times displayed at the nodes rely on previous work [7, 9, 32, 33]. Sample numbers per population are shown at the edges [10]. Black arrows indicate species-level comparisons, gray arrows subspecies-level comparisons.

The African great ape genus *Gorilla* is the first outgroup to *Pan* and *Homo* (Fig. 1). For the western gorilla species (*Gorilla gorilla*) in midwest Africa, evidence for adaptation has been reported to pathogen response and cognitive or neurodevelopmental phenotypes, as well as bitter taste receptors, possibly important to avoiding harmful substances [3, 6, 19–21]. The two subspecies of eastern gorilla (*Gorilla beringei*), inhabiting lowland forests and mountain rainforests in eastern Africa, have been proposed to show signatures of selection in genes involved in various traits including diet, behavior and immunity in either subspecies or their ancestor [19]. Whether adaptation to high altitude has a genetic basis in mountain gorillas, as is the case in humans and other species [22], still remains unclear [8], although adaptation through their microbiome has been suggested as an alternative [23].

Less attention has been paid to selection in orangutans (genus *Pongo*), the only Asian great ape (Fig. 1), partially due to smaller datasets, but also larger diversity compared to other species [24]. For the Sumatran orangutan (*Pongo abelii*), immune function [3] as well as possible candidate genes related to neurological functions [25] have been proposed. In Bornean orangutans (*Pongo pygmaeus*), physiological adaptation to fluctuating environments might have occurred [25]. Across orangutans as well as all other great ape lineages, the major histocompatibility complex (MHC) locus represents a common target of both balancing and positive selection, likely as a consequence of a complex adaptation to changing pathogen exposures [3, 6, 26].

In addition to lineage-specific signatures of natural selection, the distribution of fitness effects (DFE)the proportion of new mutations that are effectively neutral, deleterious, or beneficialis an important component in evolutionary biology, as it allows characterizing selection regimes at the genome scale [27]. Unlike selection scans, which highlight particular loci or candidate targets of natural selection, DFE inference summarizes how selection generally acts on new mutations across the genome, typically by contrasting synonymous and non-synonymous allele frequency spectra (AFS) while controlling for demographic history. Compared with detecting selection signatures, DFE estimation has been less frequently applied in non-human primates, in part because until recently only a limited set of tools and limited datasets were available. Using the GAGP dataset, previous studies have suggested that most non-synonymous mutations appear deleterious, as inferred with DFE-alpha [3], and that DFE parameters are broadly similar across species when using polyDFE [28]. However, a recent simulation-based study demonstrated that the performance of DFE inference tools is context-dependent, with different methods performing optimally in different scenarios [29]. Therefore, performing DFE inference using multiple tools is informative, as the best method based on the demographic history of great apes remains unknown.

In parallel to increasing knowledge on genetic variation in great apes beyond the GAGP, methodological developments have expanded DFE inference from AFS in single populations to joint AFS in two populations, enabling direct quantification of correlations in DFEs between populations [30]. Moreover, advances in population genetic inference have enhanced extended haplotype homozygosity (EHH)-based statistics to unphased data [31]. EHH-based methods are powerful tools for detecting recent positive selection; however, because genomic data from great apes are often unphased, previous studies could not readily apply them. In light of these advances in both genomic resources and population genetic methodology, here we perform a unified investigation of the genomic landscapes of natural selection in the great apes, focusing on recent positive selection, long-term balancing selection, and the DFE.

## Results

Throughout the analyses, we use a curated panel of high-coverage genomes from multiple studies [10], restricted to segregating sites of wild-born individuals as described in previous publications. This includes 10 bonobos and 55 chimpanzees, 24 western lowland gorillas and 21 eastern gorillas, as well as 10 Sumatran orangutans and 14 Bornean orangutans (Fig. 1).

### The landscape of recent positive selection

We used the unphased version of XP-EHH and XP-nSL statistics (see Methods) to detect signatures of recent positive selection within each lineage (Fig. S1–S28 and Tables S1–S3). Using a conservative threshold (top 0.005% scores), we determined numerous candidate genes across the different comparisons (Fig. 2). Gene ontology (GO) enrichment tests yielded very few significant GO categories after multiple testing correction (*p<*0.05), in total seven categories across all lineages and selection tests (Table 1). However, given the conservative implementation of the GO test in Gowinda [34], we also present categories nominally significant (nominal *p<*0.01) (Tables S16– S27). Numerous genes on our list overlap with genes that were presented as potential targets of positive selection in previous studies [3, 6, 13, 14, 19, 35] (Tables S58–S69).

**Fig. 2:**
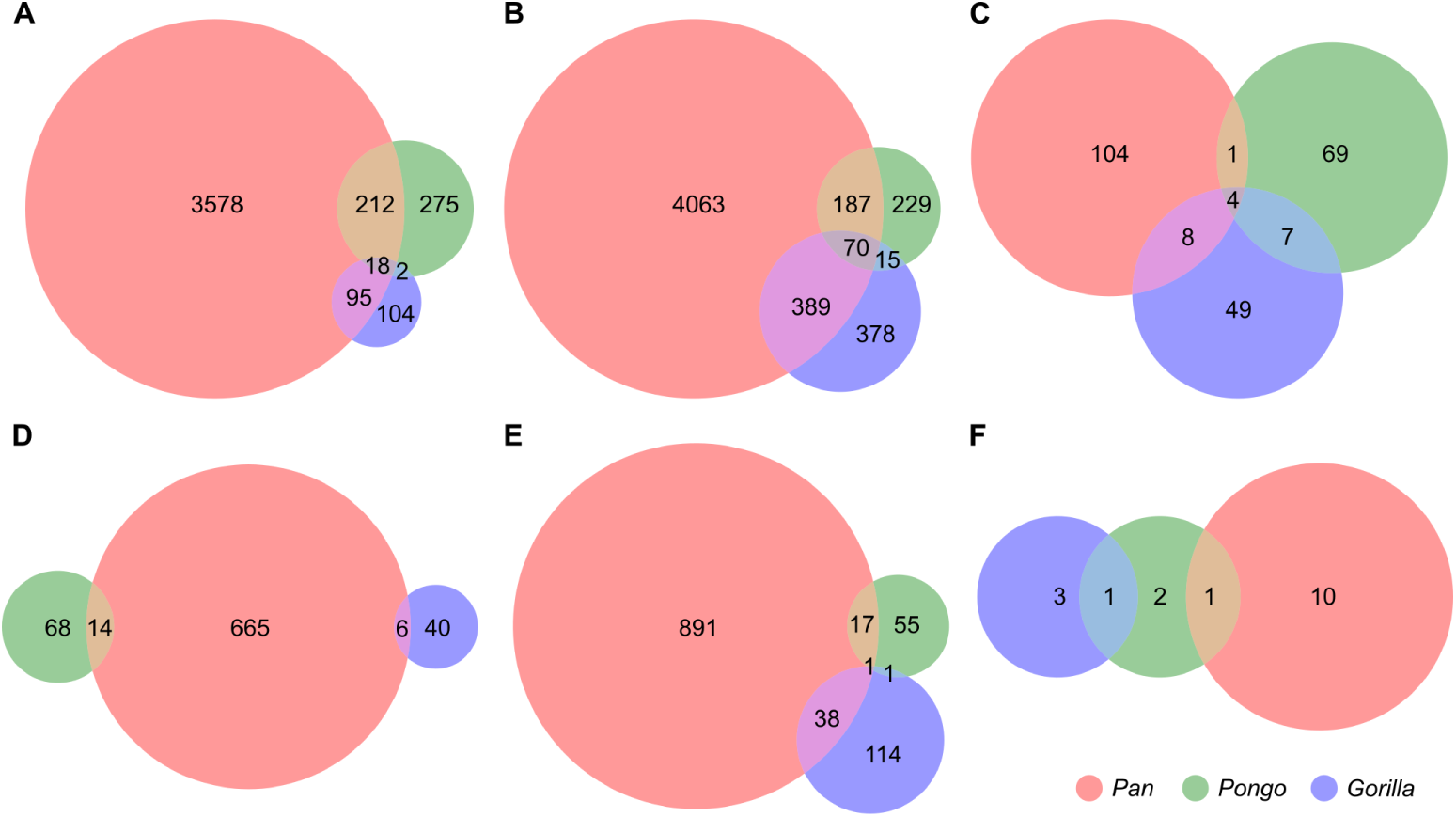
Overlap of candidate genes under recent positive selection and long-term balancing selection in *Pan*, *Pongo*, and *Gorilla*. Each genus-level set was defined as the union of genes detected in any lineage within that genus. **A** XP-EHH (unphased), top 0.05% threshold. **B** XP-nSL (unphased), top 0.05% threshold. **C** *β*^(1)^, top 0.05% threshold. **D** XP-EHH (unphased), top 0.005% threshold. **E** XP-nSL (unphased), top 0.005% threshold. **F** *β*^(1)^, top 0.005% threshold.

**Table 1:**
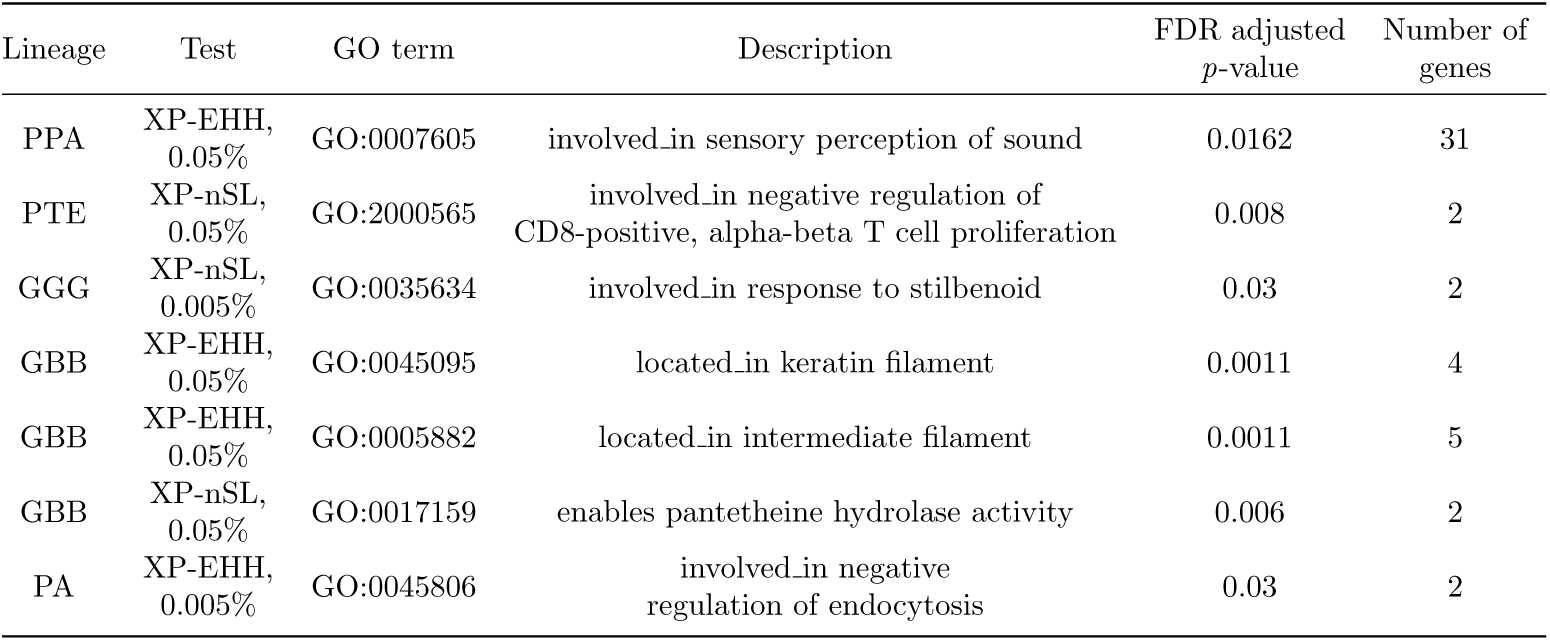
Significantly enriched GO categories after FDR correction across comparisons for positive selection tests across lineages and thresholds.

We also identify multiple instances of a selection signature on the same target genes across different species (Table S86). For instance, selection on *TTC29* in bonobos, chimpanzees and Bornean orangutans might be related to male fertility [36], given that more male competition in Bornean than Sumatran orangutans has been reported [37], or sperm competition might be high in bonobos due to their mating patterns [38]. Loci in *HERC2*, a candidate gene for selection in modern human populations associated to pigmentation [39, 40], show signatures of selection in bonobos as well as both central and eastern chimpanzees. *CNTNAP2*, a candidate gene for human lineage-specific changes [41, 42], carries loci putatively under positive selection on the bonobo, western gorilla and several chimpanzee lineages, possibly related to differences in social behaviors between species. Selection on sensory traits might have occurred as well, with *PCDH15* (Bornean orangutan, central chimpanzee) as an example of a gene involved in retina development [43]. We also found signatures of selection which were also described in selection screens in modern humans, *e.g.* for *PTPRN2* and *NRXN3* [44–46] , which might be related to phenotypes in insulin secretion and immunity for *PTPRN2* [47] or neuron development and synapse formation [48] for *NRXN3*.

### Lineage-specific signatures of positive selection

#### Selection in bonobos and chimpanzees

In bonobos, the GO category ”involved in sensory perception of sound” shows a significant enrichment in the XP-EHH-based test (Table 1). Bonobos are known for their high-pitched calls, compared to chimpanzees [49], with their inner-ear bone morphology being different from both humans and chimpanzees [50], which might be associated with this finding. Although not passing multiple testing correction, ”acts upstream of or within vocal learning” was also among the significant categories for bonobos (Table S22), which has to do with auditory ability. Two genes, *MCC*, which is overexpressed in ovary, and *SHROOM3* have been previously described as candidates for this species in multiple studies [3, 6]. A protein-coding change (Table S3) in the *KHNYN* gene might be associated with immune function [51], as well as another change in *ERAP2* for recent selection [52].

One of the candidate loci with the highest XP-EHH scores in bonobos (Table S3) might be related to reproduction traits, falling near *DPP10* [53], which is associated to age at menarche in humans and found across all four comparisons involving bonobos (Table S10). The gene *MRPS11*, associated with the onset of male puberty [54], carries another protein-coding change. *ZNF804A* carries five protein-coding changes in bonobos with a signature of positive selection. This gene is overexpressed in brain regions including hypothalamus, anterior cingulate cortex and nucleus accumbens, and has been associated with multiple neurological diseases such as Schizophrenia [55]. A locus near *LRP1B* is found as a candidate for selection (XP-EHH) across all four chimpanzee subspecies tests, a gene which has been previously described as a candidate for reproduction-related bonobo-specific traits [12]. Genes related to the specific biology of bonobos might also be suggested by the categories ”involved in negative regulation of determination of dorsal identity” and ”acts upstream of or within dorsal/ventral pattern formation” (nominally enriched, Table S22), which are possibly associated to the characteristic ventro-ventral copulation of bonobos based on their anatomical features [56, 57].

In contrast, no GO categories reached significance in the chimpanzee lineage as a whole. Loci with the highest XP-EHH scores (Table S3) in chimpanzees are involved in hearing loss (*SLC26A4* [58]), or innate immunity (*BIRC2* [59]). Protein-coding changes are found in genes related to various functions such as immune response (*CLECL1* [60] , *MS4A2* [61]), brain development (*KDM7A* [62]), or cell division (*CPAP* [63], *CENPN* [64]). A previously described candidate gene [14] is also involved in spermatogenesis (*CDYL* [65]). Recurrently supported targets of positive selection [3, 6, 13, 14] are *APBB2*, which might be involved in neuron development [66], and *FAF1* with many functions, among which virus response might have been relevant in this context [67]. However, given the divergence times of chimpanzee subspecies, the methods applied here might be more informative on the subspecies level, in particular for cross-population analyzes of positive selection.

#### Selection in chimpanzee subspecies

The only significant GO category in chimpanzees (Table 1) is for the immunity-related category ”involved in negative regulation of CD8-positive, alpha-beta T cell proliferation” in Nigeria-Cameroon chimpanzees, suggesting recent selective pressures in immune response. A variant in *FUT9*, a gene associated with susceptibility to placental malaria infection [68], shows signatures of selection in comparison to bonobos as well as Nigeria-Cameroon chimpanzees, which is interesting given previously described patterns of selection on malaria in this subspecies [18].

In both western and central chimpanzee subspecies, suggestive evidence for selection on immunity is reflected by various MHC-related categories (Table S25, nominally enriched). A top candidate locus in western chimpanzees in *SULT1C3* is in a gene involved in xenobiotic metabolic processes [69], which might reflect selection pressures related to novel environments during recent population expansions of this subspecies [70]. A protein-coding change under selection in *NPHS1* might be due to kidney functions of filtering plasma macromolecules [71].

A top candidate locus in central chimpanzees falls in the *EFCAB6* gene, which regulates the androgen receptor and may play a role in spermatogenesis [72], a trait with known differences between great ape species [73]. This subspecies also carries a protein-coding change at a candidate locus (XP-EHH) in *SHOC1*, a gene for which mutations cause azoospermia in humans and mice [74]. Further non-synonymous mutations in *ADAD2*, *CFAP70* or *SUN1* (XP-nSL) might also indicate lineage-specific selection on spermatogenesis ([75–77]). Among genes recurrently found across studies [6, 13] are *CENPP*, a spindle protein required for cell division [78], and *CORIN*, which might play a role in female fertility by promoting trophoblast invasion [79].

Protein-coding changes under selection in eastern chimpanzees (XP-EHH) encompass olfactory receptors (*OR2B6*, *OR2B2*), or a bile acid transporter (*SLC51B* [80]) which might be related to dietary adaptation. Other such changes (XP-nSL) occur in genes related to male fertility (*CFAP70* [81], *MOV10L1* [82]). A recurrently found target gene in this subspecies [3, 6, 13] is *GLG1*, a gene which is predicted to enable fibroblast growth factor binding [83], potentially influencing processes such as chondrocyte differentiation.

#### Selection in gorillas

In western gorillas, a GO category related to response to stilbenoid reached significance using the conservative threshold (0.005%) for the XP-nSL test (Table S21). Stilbenoids belong to phenolic compounds found in many plant species [84], some of which western gorillas might be able to particularly exploit [85]. Multiple further GO categories related to cholesterol, steroid or lipid metabolism are nominally enriched (Table S21), which might be related to their dietary components. In contrast, nominally enriched categories associated with the immune system in eastern gorillas (Table S20) might reflect their history of response to pathogen infections.

Notably, three out of the seven significant GO categories across lineages and tests were observed in mountain gorillas (Table 1), which might be associated with their unique adaptation to high altitude with a heavily leave-based diet. Two keratin- and intermediate filament-related categories (XP-EHH) are likely linked to their thick hair, and a pantetheine-related category (XP-nSL) is potentially associated with energy metabolism. One gene from this category is among the top 5 scores for mountain gorillas (*KRTAP25-1*) and carries a non-synonymous mutation in mountain gorillas (Ser70Cys, XP-EHH). Moreover, four non-synonymous mutations with a selection signature are observed in *KRTAP27-1*, two in *KRTAP24-1* and one in *KRTAP23-1*. Apart from those, only two more non-synonymous mutations show a signature of selection in mountain gorillas, both in *VNN1*, a gene upregulated in high-altitude pulmonary hypertension patients, likely due to a role in oxidative stress regulation under hypoxic conditions [86]. Another non-synonymous change in *FN1* (XP-nSL) might have played a role in altitude adaptation as well [87]. Other top candidate loci are in *EDNRB-AS1*, associated to the ABCD syndrome which entails pigmentation and intestinal phenotypes [88], or *MYRIP*, which might regulate insulin secretion [89].

A previously described candidate gene [19] confirmed here is the lipoprotein lipase-coding gene *LPL*, involved in lipid metabolism and potentially involved in pathways related to altitude adaptation [90].

#### Selection in orangutans

Genes under positive selection in Bornean orangutans are nominally enriched for the GO category ”involved in positive regulation of behavioral fear response” (adjusted *p*-value 0.053, Table S17, XP-nSL). This is interesting, as Bornean orangutans are known to be less social compared to Sumatran orangutans [91, 92], given that orangutans in general are considered solitary. Non-synonymous mutations putatively under selection in Bornean orangutans occur in the collagen-encoding gene *COL11A1*, which might possibly be related to differences in hair and skin morphology [93], and in *MGAM*, encoding for a starch digestion-related enzyme that might relate to differences in diet between Orangutan species [94].

In Sumatran orangutans, the GO category ”involved in negative regulation of endocytosis” is enriched for genes under selection (XP-EHH, 0.005% threshold). Being a fundamental cellular process, this could be related to various biological traits, including immune response [95]. Among top candidate loci in this species, we find *COX19*, which is overexpressed in heart and possibly related to cardiomyopathy, which is interesting considering the high prevalence of cardiovascular lesions in captive orangutans [96]. As described above, *NRXN3* has been found as target of selection in humans, while it has been previously described for Sumatran orangutans as well [25].

### The landscape of long-term balancing selection

We find signatures for balancing selection (see Methods) in immune-related loci across the different lineages (Fig. S29–S38 and Tables S28–S37), in terms of candidate genes as well as by GO enrichment within all lineages (*p*-value*<*0.05, corrected for multiple testing, Tables S38–S57). This is in line with previous studies in great apes, humans and other species [3, r6, 26, 97–99], and most likely reflects continuous exposure to the quickly evolving pathogens in the biodiversity-rich habitats of the great ape populations [100]. Hence, these are expected results for this analysis, which help refining the knowledge on balancing selection in great apes.

The HLA gene *HLA-DQB1* is found to be a target of balancing selection across most great ape lineages in this study (Table S87), emphasizing its importance in exogenous antigen presentation. This gene has been described as a target of balancing selection in previous work for both orangutan species, western gorillas, Nigeria-Cameroon, western and central chimpanzees [3, 14], which is supported and expanded to further lineages in this study (Tables S70–S79). Similarly, *HLA-DPB1* is a candidate for balancing selection in both orangutan species as well several chimpanzee subspecies, while also putatively under positive selection in chimpanzees (Table S15). Again, this confirms and expands previous findings [3, 14]. Another HLA gene (*HLA-DQA1*) is also observed across all gorilla lineages and Bornean orangutans, as previously reported [3].

Notably, the gene *CSMD1* shows signatures of balancing selection in gorilla, orangutan and chimpanzee lineages, and positive selection in bonobos, several chimpanzee subspecies, Bornean orangutans and western gorillas (Table S87). This gene has also been suggested previously to have been under positive selection [3] in chimpanzees and bonobos and carries a protein-coding change specific to bonobos [12], possibly with a function in the reproductive system [101]. However, the underlying reason for multiple signatures of selection across multiple lineages should be investigated in future studies. Among other candidate genes for balancing selection occurring across multiple lineages, *CNTNAP2* is a gene also showing candidate loci for positive selection, with loci under balancing selection in bonobos and gorillas (Table S87).

However, given the functions of this gene likely in cognition- and behavior-related traits such as autism [102], it is unclear why variation in this gene might have been under balancing selection in these lineages.

### The distribution of fitness effects

We utilized dadi, a commonly used software for demographic and DFE inference that implements the fit*∂*a*∂*i approach [103], to estimate the deleterious DFEs and gain insights into the strength of purifying selection across different lineages (see Methods). Our analysis was based on coding region variants from each lineage. Initially, we fitted a two-epoch demographic model to synonymous SNPs, under the assumption that these variants are neutral and therefore not subject to natural selection. Following this, we applied a lognormal distribution to model the DFE using non-synonymous SNPs, as these are more likely to be under purifying selection due to their potential impact on protein function. Conditional on the demographic model inferred from synonymous SNPs, we reduced the influence of demographic history on the DFE inference, ensuring that our estimation was not confounded by population history.

The maximum likelihood estimates of the AFS obtained from both the demographic model and the DFE model (Table S88 and Fig. 3) were compared with the observed data, and generally, these models fit the empirical data well (Fig. S39–S58). Building upon these DFE models, we estimated the proportions of deleterious mutations categorized by their respective selection strengths (Fig. S59–S68), using the 2*N_e_|s|* metric, where *|s|* represents the absolute selection coefficient and *N_e_* is the ancestral effective population size [29]. Specifically, we divided the mutations into four categories based on 2*N_e_|s|* values: *<* 1, 1–10, 10–100, and *>* 100, which correspond to different magnitudes of selection strengths.

**Fig. 3:**
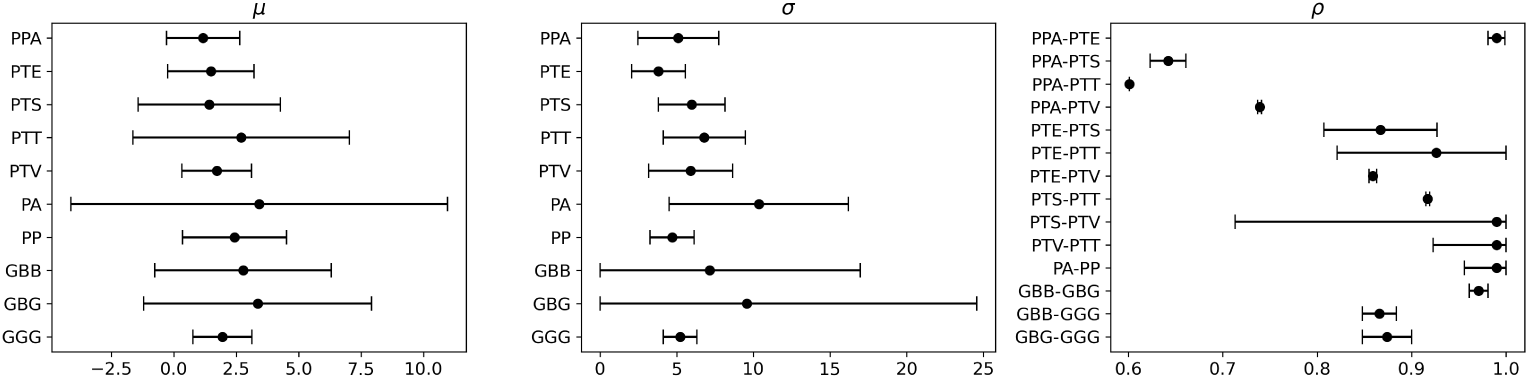
Estimated DFE parameters with 95% confidence intervals. *µ*, mean of the lognormal DFE distribution; *σ*, standard deviation of the lognormal DFE distribution; *ρ*, correlation coefficient of the joint DFE distribution, assuming a bivariate lognormal distribution

Our results reveal a consistent pattern across all the lineages, with the exception of Nigeria-Cameroon chimpanzees: the proportions of mutations in the nearly neutral (2*N_e_|s| <* 1) or extremely deleterious (2*N_e_|s| >* 100) categories were the most frequent, typically ranking first or second, with proportions usually between 25% to 45%. This pattern indicates that most lineages are subject to varying degrees of purifying selection, with a significant fraction of mutations being nearly neutral or experiencing extremely strong selection.

We further quantified the correlations of DFEs using the joint AFS from a pair of lineages (see Methods). Similar to the DFE inference, we employed a two-step approach: first, fitting a split-migration model to the synonymous AFS, and then applying a bivariate lognormal distribution to model the joint DFE based on the non-synonymous AFS, conditional on the inferred demographic model (Table S89). Comparison between the observed and inferred AFS suggests that, in general, our demographic and joint DFE models fit the data well (Fig. S69–S96). Most pairs exhibit high correlations, particularly within closely related lineages such as chimpanzees, eastern gorillas, and orangutans, typically ranging from 0.85 to 0.99 (Fig. 3). This pattern aligns with their positions in the phylogenetic tree (Fig. 1). However, with the exception of Nigeria-Cameroon chimpanzees, lower correlations are observed between bonobos and chimpanzees.

## Discussion

In this study, we investigate the landscapes of selection across great apes; unlike previous studies, including most extant lineages. Genomic diversity in the *Pan* clade is relatively well characterized [2, 7], and recently also population-level data has become available for different gorilla lineages [2, 8, 9, 104]. However, for the Cross-river sub-species (*Gorilla gorilla diehlii*), only three high-coverage genomes have been published so far [104]), which precluded an investigation of selection in this lineage. The population history of orangutans is highly complex, including a speciation process with a species currently represented by a single high-coverage genome (*Pongo tapanuliensis* [33]), which was also omitted here. If future studies provide population-scale genomic data for the lineages which could not be included here, it will be interesting to study the landscape of selection on those. Furthermore, the complex history and potentially deep population structure especially of orangutans [33] potentially impacts the performance of the approaches applied here. Hence, more data across their geographic range might help refining our observations in the future.

Still, for most lineages, for the first time patterns of natural selection were investigated using all publicly available high-coverage genomes [10] and using a unified workflow. We note that in this case all genomes were mapped to the human reference genome, in order to leverage the high quality annotations. In previous work, [6], data mapped to their respective reference genomes was used to detect recent selective sweeps. While potentially increasing accuracy, this might not allow a more direct comparison to patterns of selection in humans. Another important aspect is that the genomic diversity panel used here included 14 more orangutan individuals [33] than the previous study [105], substantially increasing the ability to detect signatures of selection in this genus. Single wild-born high-coverage western gorilla and western chimpanzee individuals were also included [104, 106]. Moreover, genomes from the GAGP of low average coverage (less than 12-fold), with cross-contamination and captive individuals were excluded here [10], since these could be confounding factors for an analysis of wild-born populations.

We present candidate genes under positive selection for the great ape lineages and interpret those given the biological context of each species. An important aspect of the sexuality of primates are sexual selection and sperm competition [107], and we find candidate loci within or near genes related to spermatogenesis or similar functions in bonobos, eastern and central chimpanzees, as well as Bornean orangutans. Biological adaptations to sensory challenges might have resulted in positive selection on genes involved in retina development or the perception of sound in several great ape lineages. Adaptation to different dietary components might have resulted in signatures of selection in western chimpanzees, western gorillas, mountain gorillas and Bornean Orangutans. Notably, we find an enrichment of keratin genes in mountain gorillas, with multiple non-synonymous changes on this lineage, possibly related to the longer and thicker hair in mountain gorillas [108]. Together with candidate loci in genes playing roles in oxidative stress regulation and lipid metabolism, selection on these genes could have facilitated the adaptation to high altitudes in mountain gorillas. This adaptation seems likely to be the result of a combination of selected traits, possibly aided by the microbiome [23]. Thus, promising routes for the understanding of this trait may open through the observations of genes with protein-coding changes and significant enrichment are presented here. Finally, candidate genes for bonobos described in this study might have contributed to their uniquely peaceful nature [109], including previously described genes with lineage-specific protein-coding changes [12]. The immune system, challenged by quickly evolving pathogens, has been the target of positive as well as balancing selection across multiple great ape lineages, in agreement with previous work [3, 6, 13, 16]. In particular, the MHC locus with the *HLA-DQB1* and *HLA-DPB1* genes is a prime example for this pattern in great apes. Other recurrent targets of both modes of selection are *CSMD1* and *CNTNAP2*, with possible roles in reproduction or behavioral traits, respectively. However, it will require further investigations to reveal the mechanisms and biological functions of candidate variants in these genes.

We also conducted DFE inference across lineages. Unlike previous studies that used DFE-alpha [110] or polyDFE [111], which assume a gamma distribution for DFEs, we assume lognormal DFE distributions in this study. Our results are consistent with previous studies [3, 21], suggesting that the majority of mutations are either nearly neutral or strongly deleterious in great apes. However, compared with previous studies in humans involving hundreds of samples [103], the dataset used in this study is still relatively small; therefore, future studies with larger sample sizes can improve precision.

Finally, we also quantified the correlations of DFEs. Similar to a previous study [30] on humans, fruit flies, and wild tomatoes, the correlations were high within closely related lineages, while comparably more diverged lineages within a genus showed much lower correlations. One exception is the pair of bonobos and Nigeria-Cameroon chimpanzees, with an unexpectedly high correlation, which may be due to complex demographic factors. Given the overall complexity of the demographic history of the species in this clade [2, 7, 9, 33], future studies with fine-tuned demographic models are more data may improve such estimates. This will likely be promoted by integrating other evolutionary processes such as introgression, or applying emerging techniques such as deep learning in the field of population genetics [112, 113]. Expanding beyond great apes, future work may also leverage large-scale datasets of other primate species [114].

## Conclusion

In conclusion, our study provides a comprehensive picture of the landscape of natural selection in great apes, using high-coverage whole genomes covering most of the known lineages. Our results confirmed recurrent positive selection targets such as immune- or reproduction-related genes, while also detecting novel genes and gene categories which are likely associated with the unique biological traits of each lineage. Immune response-related genes were also main long-term balancing selection targets, as expected, in a complex interplay with positive selection signatures on some of the same genes. Finally, we provide a thorough analysis of the DFE, supporting that most mutations are either nearly neutral or strongly deleterious, while highly correlated among closely related lineages.

## Methods

### Data preprocessing

A curated dataset of great ape genomes was analyzed, with all variants mapped to the human reference genome hg38 and processed accordingly [10]. Only samples from wild individuals were included in the analysis. Only bi-allelic single nucleotide polymorphisms (SNPs) were considered. For *Gorilla* and *Pongo*, SNPs that were heterozygous in all individuals across the species, within a population pair, or within a single population, were excluded to remove loci likely located in repetitive regions, as these species lack comprehensive repeat annotations. To polarize the variants, the UCSC Genome Browser liftover chain from hg38 to rheMac10 was used to convert hg38 coordinates to the rheMac10 reference genome and to assign the corresponding allele in rheMac10 as the ancestral allele. The high-quality macaque genome served as an out-group, under the assumption that variable sites within each genus (*Gorilla*, *Pan*, and *Pongo*) represent mutations specific to that lineage.

### Detection of recent positive selection

To identify candidate genes under recent positive selection, selscan (version 2.0.3) was used to compute unphased extended haplotype homozygosity (EHH)-based statistics [31]. Specifically, cross-population statistics XP-EHH and XP-nSL were calculated for all pairwise population comparisons within each species, as these methods are robust to small sample sizes and unphased data [31, 115, 116]. Only SNPs with a minor allele frequency (MAF) greater than 0.05 were used as core SNPs. All other parameters were set to the default values in selscan. The resulting scores were normalized using default settings based on genome-wide distributions via the bundled norm program (version 1.3.0). Following normalization, SNPs were ranked by the absolute values of their scores, and the top 0.05% and 0.005% were selected as candidate sites.

### Detection of long-term balancing selection

To identify candidate genes under long-term balancing selection in each population, BetaScan was applied to compute the *β*^(1)^ statistic [117]. The analysis included only SNPs with a MAF greater than 0.05. SNPs located within regions annotated by the RepeatMasker, simple repeats, or segmental duplications tables from the UCSC Table Browser were excluded. SNPs were also removed with exact Hardy-Weinberg equilibrium *p*-values less than 10*^−^*^3^, as determined by PLINK 1.9 [118]. For *β*^(1)^ score calculation, only SNPs with allele frequencies exceeding 0.15 were used as core sites. SNPs were ordered according to their *β*^(1)^ scores, and the highest-scoring 0.05% and 0.005% were retained as candidate sites.

### Lineage-specific candidates

Candidate variants from the XP-EHH and XP-nSL tests were stratified by lineage according to the sign of the statistic, which indicates the direction of the signal (positive for one population and negative for the other). Positive values were assigned to the population listed first in the comparison, denoted as (+), while negative values were assigned to the second population, denoted as (*−*). Based on this criterion, candidate variants were grouped into lineages as follows: bonobo (PPA), defined by PPA-PTE (+), PPA-PTS (+), PPA-PTT (+), PPA-PTV (+); chimpanzees (PT), defined by PPA-PTE (*−*), PPA-PTS (*−*), PPA-PTT (*−*), PPA-PTV (*−*); NigerianCameroonian chimpanzees (PTE), defined by PTE-PTS(+), PTE-PTV(+), and PTE-PTT(+); eastern chimpanzees (PTS), defined by PTS-PTV(+), PTS-PTT(+), and PTE-PTS(*−*); western chimpanzees (PTV), defined by PTV-PTT(+), PTE-PTV(*−*), and PTS-PTV(*−*); central chimpanzees (PTT), defined by PTE-PTT(*−*), PTS-PTT(*−*), and PTV-PTT(*−*). For gorillas, eastern gorillas (GB) were defined by GBB-GGG(+) and GBG-GGG(+), while western lowland gorillas (GGG) were defined by GBB-GGG(*−*) and GBG-GGG(*−*); mountain gorillas (GBB) were defined by GBB-GBG(+), and eastern lowland gorillas (GBG) by GBB-GBG(*−*). For orangutans, Sumatran orangutans (PA) were defined by PA-PP(+), and Bornean orangutans (PP) by PA-PP(*−*).

### Gene annotation and GO enrichment analysis

ANNOVAR (version 2022Oct05) [119] was used to annotate all SNPs with RefSeq, dbNSFP v4.2c, and dbSNP build 150 [120–122], based on the hg38 human reference genome. Gowinda (version 1.12) [34] was run in gene mode, as recommended by a previous study [18], to perform GO enrichment analysis for the candidate SNPs identified by selscan or betascan. GO term association files were downloaded from the NCBI FTP site. Gowinda was chosen because it accounts for differences in gene length. Literature search of functional aspects and gene expression was performed using the GeneCards database [123].

### Inference of the DFEs

To infer the DFEs of new mutations in the coding regions of each population from the site frequency spectrum (SFS), the Python package dadi (version 2.3.0) was employed via its command-line interface dadi-cli (version 0.9.1) [124, 125]. Singletons were masked in the SFS. To control for the confounding effects of demographic history on the SFS, a two-epoch demographic model was inferred using synonymous SNPs, followed by estimation of the deleterious DFE from non-synonymous SNPs, assuming a lognormal distribution [103]. The two-epoch model includes two parameters: the ratio of contemporary to *N_e_* and the time in the past when the size change occurred, expressed in units of 2*N_e_* generations. The deleterious DFE is modeled as a lognormal distribution of the population-scaled selection coefficients (2*N_e_s*), parameterized by the mean and standard deviation. An additional parameter accounting for ancestral allele misidentification was included in both demographic and DFE inference.

The grid size for numeric optimization was set to [300, 400, 500], and 50 gamma points were used to generate the cache of SFS with different selection coefficients. For the demographic model, initial parameter values were [0.5, 0.5, 0.5], with upper bounds [10, 10, 1] and lower bounds [10*^−^*^2^, 10*^−^*^2^, 10*^−^*^2^]. For the DFE, initial parameters were [5, 5, 0.5], with upper bounds [100, 100, 1] and lower bounds [10*^−^*^2^, 10*^−^*^2^, 10*^−^*^2^]. The ratio of non-synonymous to synonymous mutations was set to 2.31 based on an estimate from humans [126]. For both demographic and DFE inference, 100 independent optimizations were first performed. In each run, each initial parameter was perturbed by multiplying it with 2^(2*U*^*^−^*^1)^, where *U* is a uniform random value between 0 and 1. The top 10 parameter sets with the highest likelihood were subsequently used as starting points for 100 additional optimizations using the same perturbation scheme. The optimization process was then continued until convergence was achieved, as assessed with the maximum percentage difference in log-likelihoods set to 0.0001 relative to the best log-likelihood observed. To estimate confidence intervals of the inferred DFE parameters, an approach based on the Godambe information matrix was utilized with a step size of 0.001 for numerical derivatives and 100 bootstrap datasets generated by dividing the genome into 10 Mb chunks [127].

### Quantification of the DFE correlations

To quantify correlations in the DFE between populations, a splitmigration demographic model was first inferred using synonymous SNPs, and the joint deleterious DFE was subsequently estimated from nonsynonymous SNPs under the assumption of a bivariate log-normal distribution. The splitmigration model included four parameters: the ratio of the size of population 1 to *N_e_*, the ratio of the size of population 2 to *N_e_*, the split time in units of 2*N_e_*generations, and the migration rate scaled by 2*N_e_*. The parameter for ancestral allele misidentification was also included.

For the demographic model, the initial parameter values were set to [0.5, 0.5, 0.5, 0.5, 0.5], with upper bounds [10, 10, 10, 10, 1] and lower bounds [10*^−^*^2^, 10*^−^*^2^, 10*^−^*^2^, 10*^−^*^2^, 10*^−^*^2^]. For the joint DFE inference, the DFE parameters of each population were fixed to the values obtained from the single-population DFE analyses, and only the correlation coefficient and the ancestral allele misidentification parameter were estimated. Initial values were [0.5, 0.5], with upper bounds [0.99, 1] and lower bounds [0.99, 10*^−^*^2^]. All other parameters for numerical optimization and confidence interval estimation in dadi followed those used in the single-population DFE inference.

## Acknowledgements.

The authors thank the Life Science Compute Cluster at the University of Vienna and the Multi-Site Computer Austria for providing computing resources.

## Funding

This project has been funded by the Vienna Science and Technology Fund (WWTF) [10.47379/VRG20001] to M.K. S.H. was supported by the Austrian Science Fund (FWF) [10.55776/ESP546].

## Authors contributions

X.H. designed the study. X.H., S.C., S.H., M.K. analyzed the data and wrote the manuscript.

## Data availability

The curated great ape genomes and corresponding metadata can be downloaded from https://phaidra.univie.ac.at/pfsa/o 2066302/, last access August 26, 2025. The FASTA file for the reference genome hg38 can be found in https://hgdownload.soe.ucsc.edu/goldenPath/hg38/bigZips/hg38.fa.gz, last accessed August 26, 2025. The FASTA file for the reference genome rheMac10 can be found in https://hgdownload.soe.ucsc.edu/goldenPath/rheMac10/bigZips/rheMac10.fa.gz, last accessed August 26, 2025. The file for liftover chain from hg38 to rheMac10 can be found in https://hgdownload.soe.ucsc.edu/goldenPath/ hg38/liftOver/hg38ToRheMac10.over.chain.gz, last accessed August 26, 2025. The RepeatMasker Track from the UCSC Genome Browser can be found in https://hgdownload.soe.ucsc.edu/goldenPath/hg38/database/rmsk.txt.gz, last accessed August 26, 2025. The Segmental Dups Track from the UCSC Genome Browser can be found in https://hgdownload.soe.ucsc.edu/goldenPath/ hg38/database/genomicSuperDups.txt.gz, last accessed August 26, 2025. The Simple Repeats Track from the UCSC Genome Browser can be found in https://hgdownload.soe.ucsc.edu/goldenPath/hg38/database/simpleRepeat.txt.gz, last accessed August 26, 2025. Genome annotation can be found in https://ftp.ncbi. nih.gov/genomes/refseq/vertebrate mammalian/Homo sapiens/annotation releases/ GCF 000001405.40-RS 2024 08/GCF 000001405.40 GRCh38.p14 genomic.gtf.gz, last accessed August 26, 2025. GO term association can be found in https://ftp.ncbi.nlm.nih.gov/gene/DATA/gene2go.gz, last accessed August 26, 2025. ANNOVAR can be downloaded from https://www.openbioinformatics.org/annovar/ annovar download form.php, last accessed August 26, 2025. selscan can be downloaded from https://github.com/szpiech/selscan/archive/refs/tags/v2.0.3.tar.gz, last accessed August 26. BetaScan can be downloaded from https://github.com/ ksiewert/BetaScan, last accessed August 26, 2025. dadi-cli can be downloaded from https://anaconda.org/conda-forge/dadi-cli, last accessed August 26, 2025. Gowinda can be downloaded from https://sourceforge.net/projects/gowinda/, last accessed August 26, 2025. The Snakemake workflow for reproducing the analysis can be found in https://github.com/admixVIE/gas, last accessed August 26, 2025.

## Competing interests

The authors declare no conflict of interest.

## Notes

### Competing Interest Statement

The authors have declared no competing interest.

## References

[1] Roos, C., Seshadri, L., Zhang, L., Harris, R.A., Raveendran, M., Cuadros Espinoza, S.H., Kuderna, L.F.K., Manu, S., Umapathy, G., Boubli, J.P., Wu, H., Kuang, W., Yu, L., Zhao, X., Liu, Z., Zhu, P., Qi, J., Zhou, X., Li, M., Shao, Y., Wu, D., Farh, K.K.-H., Marques-Bonet, T., Zinner, D., Rogers, J.: Genomic basis of non-human-primate diversity and adaptation. Nature Reviews Biodiversity 1(6), 353–370 (2025) 10.1038/s44358-025-00039-8

[2] Prado-Martinez, J., Sudmant, P.H., Kidd, J.M., Li, H., Kelley, J.L., Lorente-Galdos, B., Veeramah, K.R., Woerner, A.E., O’Connor, T.D., Santpere, G., Cagan, A., Theunert, C., Casals, F., Laayouni, H., Munch, K., Hobolth, A., Halager, A.E., Malig, M., Hernandez-Rodriguez, J., Hernando-Herraez, I., Prüfer, K., Pybus, M., Johnstone, L., Lachmann, M., Alkan, C., Twigg, D., Petit, N., Baker, C., Hormozdiari, F., Fernandez-Callejo, M., Dabad, M., Wilson, M.L., Stevison, L., Camprubí, C., Carvalho, T., Ruiz-Herrera, A., Vives, L., Mele, M., Abello, T., Kondova, I., Bontrop, R.E., Pusey, A., Lankester, F., Kiyang, J.A., Bergl, R.A., Lonsdorf, E., Myers, S., Ventura, M., Gagneux, P., Comas, D., Siegismund, H., Blanc, J., Agueda-Calpena, L., Gut, M., Fulton, L., Tishkoff, S.A., Mullikin, J.C., Wilson, R.K., Gut, I.G., Gonder, M.K., Ryder, O.A., Hahn, B.H., Navarro, A., Akey, J.M., Bertranpetit, J., Reich, D., Mailund, T., Schierup, M.H., Hvilsom, C., Andŕes, A.M., Wall, J.D., Bustamante, C.D., Hammer, M.F., Eichler, E.E., Marques-Bonet, T.: Great ape genetic diversity and population history. Nature 499(7459), 471–5 (2013) 10.1038/nature12228

[3] Cagan, A., Theunert, C., Laayouni, H., Santpere, G., Pybus, M., Casals, F., Pr?fer, K., Navarro, A., Marques-Bonet, T., Bertranpetit, J., Andr?s, A.M.: Natural Selection in the Great Apes. Molecular Biology and Evolution 33(12), 3268–3283 (2016) 10.1093/molbev/msw215

[4] Nam, K., Munch, K., Mailund, T., Nater, A., Greminger, M.P., Krützen, M., Marqùes-Bonet, T., Schierup, M.H.: Evidence that the rate of strong selective sweeps increases with population size in the great apes. Proceedings of the National Academy of Sciences 114(7), 1613–1618 (2017) 10.1073/pnas.1605660114

[5] Zhao, S., Zhang, T., Liu, Q., Wu, H., Su, B., Shi, P., Chen, H.: Identifying Lineage-Specific Targets of Natural Selection by a Bayesian Analysis of Genomic Polymorphisms and Divergence from Multiple Species. Molecular Biology and Evolution 36(6), 1302–1315 (2019) 10.1093/molbev/msz046

[6] Yoo, D., Rhie, A., Hebbar, P., Antonacci, F., Logsdon, G.A., Solar, S.J., Antipov, D., Pickett, B.D., Safonova, Y., Montinaro, F., Luo, Y., Malukiewicz, J., Storer, J.M., Lin, J., Sequeira, A.N., Mangan, R.J., Hickey, G., Monfort Anez, G., Balachandran, P., Bankevich, A., Beck, C.R., Biddanda, A., Borchers, M., Bouffard, G.G., Brannan, E., Brooks, S.Y., Carbone, L., Carrel, L., Chan, A.P., Crawford, J., Diekhans, M., Engelbrecht, E., Feschotte, C., Formenti, G., Garcia, G.H., Gennaro, L., Gilbert, D., Green, R.E., Guarracino, A., Gupta, I., Haddad, D., Han, J., Harris, R.S., Hartley, G.A., Harvey, W.T., Hiller, M., Hoekzema, K., Houck, M.L., Jeong, H., Kamali, K., Kellis, M., Kille, B., Lee, C., Lee, Y., Lees, W., Lewis, A.P., Li, Q., Loftus, M., Loh, Y.H.E., Loucks, H., Ma, J., Mao, Y., Martinez, J.F.I., Masterson, P., McCoy, R.C., McGrath, B., McKinney, S., Meyer, B.S., Miga, K.H., Mohanty, S.K., Munson, K.M., Pal, K., Pennell, M., Pevzner, P.A., Porubsky, D., Potapova, T., Ringeling, F.R., Rocha, J.L., Ryder, O.A., Sacco, S., Saha, S., Sasaki, T., Schatz, M.C., Schork, N.J., Shanks, C., Smeds, L., Son, D.R., Steiner, C., Sweeten, A.P., Tassia, M.G., Thibaud-Nissen, F., Torres-Gonźalez, E., Trivedi, M., Wei, W., Wertz, J., Yang, M., Zhang, P., Zhang, S., Zhang, Y., Zhang, Z., Zhao, S.A., Zhu, Y., Jarvis, E.D., Gerton, J.L., Rivas-Gonźalez, I., Paten, B., Szpiech, Z.A., Huber, C.D., Lenz, T.L., Konkel, M.K., Yi, S.V., Canzar, S., Watson, C.T., Sudmant, P.H., Molloy, E., Garrison, E., Lowe, C.B., Ventura, M., ONeill, R.J., Koren, S., Makova, K.D., Phillippy, A.M., Eichler, E.E.: Complete sequencing of ape genomes. Nature 641(8062), 401–418 (2025) 10.1038/s41586-025-08816-3

[7] De Manuel, M., Kuhlwilm, M., Frandsen, P., Sousa, V.C.V.C.V.C., Desai, T., Prado-Martinez, J., Hernandez-Rodriguez, J., Dupanloup, I., Lao, O., Hallast, P., Schmidt, J.M.J.M., Heredia-Genestar, J.M.J.M., Benazzo, A., Barbujani, G., Peter, B.M.B.M., Kuderna, L.F.K.L.F.K., Casals, F., Angedakin, S., Arandjelovic, M., Boesch, C., Kühl, H., Vigilant, L., Langergraber, K., Novembre, J., Gut, M., Gut, I., Navarro, A., Carlsen, F., Andŕes, A.M.A.M.A.M., Siegismund, H.R.H.R., Scally, A., Excoffier, L., Tyler-Smith, C., Castellano, S., Xue, Y., Hvilsom, C., Marques-Bonet, T.: Chimpanzee genomic diversity reveals ancient admixture with bonobos. Science 354(6311) (2016) 10.1126/science.aag2602

[8] Xue, Y., Prado-martinez, J., Sudmant, P.H., Narasimhan, V., Ayub, Q., Szpak, M., Frandsen, P., Chen, Y., Yngvadottir, B., Cooper, D.N., Manuel, M.D., Hernandez-rodriguez, J., Lobon, I., Siegismund, H.R., Pagani, L., Quail, M.A., Hvilsom, C., Mudakikwa, A., Eichler, E.E., Cranfield, M.R., Marques-bonet, T.: Mountain gorilla genomes reveal the impact of long-term population decline and inbreeding. Science 348(6231), 242–245 (2015) 10.1126/science. aaa3952

[9] Pawar, H., Rymbekova, A., Cuadros-Espinoza, S., Huang, X., Manuel, M., Valk, T., Lobon, I., Alvarez-Estape, M., Haber, M., Dolgova, O., Han, S., Esteller-Cucala, P., Juan, D., Ayub, Q., Bautista, R., Kelley, J.L., Cornejo, O.E., Lao, O., Andŕes, A.M., Guschanski, K., Ssebide, B., Cranfield, M., Tyler-Smith, C., Xue, Y., Prado-Martinez, J., Marques-Bonet, T., Kuhlwilm, M.: Ghost admixture in eastern gorillas. Nature Ecology & Evolution (2023) 10.1038/s41559-023-02145-2

[10] Han, S., Riyahi, S., Huang, X., Kuhlwilm, M.: A curated great ape genome diversity panel. bioRxiv, 2025–0218638799 (2025) 10.1101/2025.02.18.638799

[11] Han, S., Filippo, C., Parra, G., Meneu, J.R., Laurent, R., Frandsen, P., Hvilsom, C., Gronau, I., Marques-Bonet, T., Kuhlwilm, M., Andŕes, A.M.: Deep genetic substructure within bonobos. Current Biology (2024) 10.1016/j.cub.2024.09.043

[12] Han, S., Andŕes, A.M., Marques-Bonet, T., Kuhlwilm, M.: Genetic variation in Pan species is shaped by demographic history and harbors lineage-specific functions. Genome Biology and Evolution, 047 (2019) 10.1093/gbe/evz047

[13] Nye, J., Mondal, M., Bertranpetit, J., Laayouni, H.: A fully integrated machine learning scan of selection in the chimpanzee genome. NAR Genomics and Bioinformatics 2(3), 061 (2020) 10.1093/nargab/lqaa061

[14] Zhao, S., Chi, L., Chen, H.: CEGA: a method for inferring natural selection by comparative population genomic analysis across species. Genome Biology 24(1), 219 (2023) 10.1186/s13059-023-03068-8

[15] Nielsen, R., Bustamante, C., Clark, A.G., Glanowski, S., Sackton, T.B., Hubisz, M.J., Fledel-Alon, A., Tanenbaum, D.M., Civello, D., White, T.J.J., Sninsky, J., Adams, M.D., Cargill, M.: A Scan for Positively Selected Genes in the Genomes of Humans and Chimpanzees. PLOS Biology 3(6), 170 (2005) 10.1371/journal.pbio.0030170

[16] Pawar, H., Ostridge, H.J., Schmidt, J.M., Andŕes, A.M.: Genetic adaptations to SIV across chimpanzee populations. PLOS Genetics 18(8), 1010337 (2022) 10.1371/journal.pgen.1010337

[17] Schmidt, J.M., Manuel, M., Marques-Bonet, T., Castellano, S., Andŕes, A.M.: The impact of genetic adaptation on chimpanzee subspecies differentiation. PLOS Genetics 15(11), 1008485 (2019) 10.1371/journal.pgen.1008485

[18] Ostridge, H.J., Fontsere, C., Lizano, E., Soto, D.C., Schmidt, J.M., Saxena, V., Alvarez-Estape, M., Barratt, C.D., Gratton, P., Bocksberger, G., Lester, J.D., Dieguez, P., Agbor, A., Angedakin, S., Assumang, A.K., Bailey, E., Barubiyo, D., Bessone, M., Brazzola, G., Chancellor, R., Cohen, H., Coupland, ., Danquah, E., Deschner, T., Dotras, L., Dupain, J., Egbe, V.E., Granjon, A.-C., Head, J., Hedwig, D., Hermans, V., Hernandez-Aguilar, R.A., Jeffery, K.J., Jones, S., Junker, J., Kadam, P., Kaiser, M., Kalan, A.K., Kambere, M., Kienast, I., Kujirakwinja, D., Langergraber, K.E., Lapuente, J., Larson, B., Laudisoit, A., Lee, K.C., Llana, M., Maretti, G., Martín, R., Meier, A.C., Morgan, D., Neil, E., Nicholl, S., Nixon, S., Normand, E., Orbell, C., Ormsby, L.J., Orume, R., Pacheco, L., Preece, J., Regnaut, S., Robbins, M.M., Rundus, A., Sanz, C., Sciaky, L., Sommer, V., Stewart, F.A., Tagg, N., Tédonzong, L.R., Schijndel, J., Vendras, E., Wessling, E.G., Willie, J., Wittig, R.M., Yuh, Y.G., Yurkiw, K., Vigilant, L., Piel, A.K., Boesch, C., Kühl, H.S., Dennis, M.Y., Marques-Bonet, T., Arandjelovic, M., Andŕes, A.M.: Local genetic adaptation to habitat in wild chimpanzees. Science 387(6730), 7954 (2025) 10.1126/science.adn7954

[19] Valk, T., Jensen, A., Caillaud, D., Guschanski, K.: Comparative genomic analyses provide new insights into evolutionary history and conservation genomics of gorillas. BMC Ecology and Evolution 24(1), 14 (2024) 10.1186/s12862-023-02195-x

[20] Scally, A., Dutheil, J.Y., Hillier, L.W., Jordan, G.E., Goodhead, I., Herrero, J., Hobolth, A., Lappalainen, T., Mailund, T., Marques-Bonet, T., McCarthy, S., Montgomery, S.H., Schwalie, P.C., Tang, Y.A., Ward, M.C., Xue, Y., Yngvadottir, B., Alkan, C., Andersen, L.N., Ayub, Q., Ball, E.V., Beal, K., Bradley, B.J., Chen, Y., Clee, C.M., Fitzgerald, S., Graves, T.A., Gu, Y., Heath, P., Heger, A., Karakoc, E., Kolb-Kokocinski, A., Laird, G.K., Lunter, G., Meader, S., Mort, M., Mullikin, J.C., Munch, K., O’Connor, T.D., Phillips, A.D., Prado-Martinez, J., Rogers, A.S., Sajjadian, S., Schmidt, D., Shaw, K., Simpson, J.T., Stenson, P.D., Turner, D.J., Vigilant, L., Vilella, A.J., Whitener, W., Zhu, B., Cooper, D.N., Jong, P., Dermitzakis, E.T., Eichler, E.E., Flicek, P., Goldman, N., Mundy, N.I., Ning, Z., Odom, D.T., Ponting, C.P., Quail, M.A., Ryder, O.A., Searle, S.M., Warren, W.C., Wilson, R.K., Schierup, M.H., Rogers, J., Tyler-Smith, C., Durbin, R.: Insights into hominid evolution from the gorilla genome sequence. Nature 483(7388), 169–175 (2012) 10.1038/nature10842

[21] McManus, K.F., Kelley, J.L., Song, S., Veeramah, K., Woerner, A.E., Stevison, L.S., Ryder, O.A., Project, G.A.G., Kidd, J.M., Wall, J.D., Bustamante, C.D., Hammer, M.F.: Inference of Gorilla demographic and selective history from whole genome sequence data. Molecular Biology and Evolution 32, 600–612 (2015) 10.1093/molbev/msu394

[22] Witt, K.E., Huerta-Śanchez, E.: Convergent evolution in human and domesticate adaptation to high-altitude environments. Philosophical Transactions of the Royal Society B: Biological Sciences 374(1777), 20180235 (2019) 10.1098/rstb.2018.0235

[23] Moraitou, M., Forsythe, A., Fellows Yates, J.A., Brealey, J.C., Warinner, C., Guschanski, K.: Ecology, Not Host Phylogeny, Shapes the Oral Microbiome in Closely Related Species. Molecular Biology and Evolution 39(12), 263 (2022) 10.1093/molbev/msac263

[24] Kuhlwilm, M., Manuel, M.d., Nater, A., Greminger, M.P., Krützen, M., Marques-Bonet, T.: Evolution and demography of the great apes. Current Opinion in Genetics & Development (2016) 10.1016/j.gde.2016.09. 005

[25] Mattle-Greminger, M.P., Bilgin Sonay, T., Nater, A., Pybus, M., Desai, T., Valles, G., Casals, F., Scally, A., Bertranpetit, J., Marques-Bonet, T., Schaik, C.P., Anisimova, M., Krützen, M.: Genomes reveal marked differences in the adaptive evolution between orangutan species. Genome Biology 19(1), 193 (2018) 10.1186/s13059-018-1562-6

[26] Teixeira, J.C., Filippo, C., Weihmann, A., Meneu, J.R., Racimo, F., Dannemann, M., Nickel, B., Fischer, A., Halbwax, M., Andre, C., Atencia, R., Meyer, M., Parra, G., Pääbo, S., Andŕes, A.M.: Long-Term Balancing Selection in LAD1 Maintains a Missense Trans-Species Polymorphism in Humans, Chimpanzees, and Bonobos. Molecular Biology and Evolution 32(5), 1186–1196 (2015) 10.1093/molbev/msv007

[27] Eyre-Walker, A., Keightley, P.D.: The distribution of fitness effects of new mutations. Nature Reviews Genetics 8, 610–618 (2007) 10.1038/nrg2146

[28] Castellano, D., Macìa, M.C., Tataru, P., Bataillon, T., Munch, K.: Comparison of the Full Distribution of Fitness Effects of New Amino Acid Mutations Across Great Apes. Genetics 213(3), 953–966 (2019) 10.1534/genetics.119.302494

[29] Gower, G., Pope, N.S., Rodrigues, M.F., Tittes, S., Tran, L.N., Alam, O., Cavassim, M.I.A., Fields, P.D., Haller, B.C., Huang, X., Jeffrey, B., Korfmann, K., Kyriazis, C.C., Min, J., Rebollo, I., Rehmann, C.T., Small, S.T., Smith, C.C.R., Tsambos, G., Wong, Y., Zhang, Y., Huber, C.D., Gorjanc, G., Ragsdale, A., Gronau, I., Gutenkunst, R.N., Kelleher, J., Lohmueller, K.E., Schrider, D.R., Ralph, P.L., Kern, A.D.: Accessible, realistic genome simulation with selection using stdpopsim. bioRixv (2025) 10.1101/2025.03.23.644823

[30] Huang, X., Fortier, A.L., Coffman, A.J., Struck, T.J., Irby, M.N., James, J.E., Len-Burguete, J.E., Ragsdale, A.R., Gutenkunst, R.N.: Inferring genome-wide correlations of mutation fitness effects between populations. Mol Biol Evol 38, 4588–4602 (2021) 10.1093/molbev/msab162

[31] Szpiech, Z.A.: selscan 2.0: scanning for sweeps in unphased data. Bioinformatics 40, 006 (2024) 10.1093/bioinformatics/btae006

[32] Besenbacher, S., Hvilsom, C., Marques-Bonet, T., Mailund, T., Schierup, M.H.: Direct estimation of mutations in great apes reconciles phylogenetic dating. Nature Ecology & Evolution 3(2), 286–292 (2019) 10.1038/s41559-018-0778-x

[33] Nater, A., Mattle-Greminger, M.P., Nurcahyo, A., Nowak, M.G., Manuel, M., Desai, T., Groves, C., Pybus, M., Sonay, T.B., Roos, C., Lameira, A.R., Wich, S.A., Askew, J., Davila-Ross, M., Fredriksson, G., Valles, G., Casals, F., Prado-Martinez, J., Goossens, B., Verschoor, E.J., Warren, K.S., Singleton, I., Marques, D.A., Pamungkas, J., Perwitasari-Farajallah, D., Rianti, P., Tuuga, A., Gut, I.G., Gut, M., Orozco-terWengel, P., Schaik, C.P., Bertranpetit, J., Anisimova, M., Scally, A., Marques-Bonet, T., Meijaard, E., Krützen, M.: Morphometric, Behavioral, and Genomic Evidence for a New Orangutan Species. Current Biology 27(22), 3487–3498 (2017) 10.1016/j.cub.2017.09.047

[34] Kofler, R., Schlötterer, C.: Gowinda: unbiased analysis of gene set enrichment for genome-wide association studies. Bioinformatics 28(15), 2084–2085 (2012) 10.1093/bioinformatics/bts315

[35] Yousaf, A., Liu, J., Ye, S., Chen, H.: Current Progress in Evolutionary Comparative Genomics of Great Apes. Frontiers in Genetics 12 (2021) 10.3389/fgene.2021.657468

[36] Lorès, P., Dacheux, D., Kherraf, Z.-E., Nsota Mbango, J.-F., Coutton, C., Stouvenel, L., Ialy-Radio, C., Amiri-Yekta, A., Whitfield, M., Schmitt, A., Cazin, C., Givelet, M., Ferreux, L., Fourati Ben Mustapha, S., Halouani, L., Marrakchi, O., Daneshipour, A., El Khouri, E., Do Cruzeiro, M., Favier, M., Guillonneau, F., Chaudhry, M., Sakheli, Z., Wolf, J.-P., Patrat, C., Gacon, G., Savinov, S.N., Hosseini, S.H., Robinson, D.R., Zouari, R., Ziyyat, A., Arnoult, C., Dulioust, E., Bonhivers, M., Ray, P.F., Touŕe, A.: Mutations in ¡em¿TTC29¡/em¿, Encoding an Evolutionarily Conserved Axonemal Protein, Result in Asthenozoospermia and Male Infertility. The American Journal of Human Genetics 105(6), 1148–1167 (2019) 10.1016/j.ajhg.2019.10.007

[37] Noordwijk, M.A., LaBarge, L.R., Kunz, J.A., Marzec, A.M., Spillmann, B., Ackermann, C., Rianti, P., Vogel, E.R., Atmoko, S.S.U., Kruetzen, M., Schaik, C.P.: Reproductive success of Bornean orangutan males: scattered in time but clustered in space. Behavioral Ecology and Sociobiology 77(12), 134 (2023) 10.1007/s00265-023-03407-6

[38] Marvan, R., Stevens, J.M.G., Roeder, A.D., Mazura, I., Bruford, M.W., Ruiter, J.R.: Male Dominance Rank, Mating and Reproductive Success in Captive Bonobos (Pan paniscus). Folia Primatologica 77(5), 364–376 (2006) 10.1159/000093702

[39] Lao, O., Gruijter, J.M., Duijn, K., Navarro, A., Kayser, M.: Signatures of Positive Selection in Genes Associated with Human Skin Pigmentation as Revealed from Analyses of Single Nucleotide Polymorphisms. Annals of Human Genetics 71(3), 354–369 (2007) 10.1111/j.1469-1809.2006.00341.x

[40] Huang, X., Wang, S., Jin, L., He, Y.: Dissecting dynamics and differences of selective pressures in the evolution of human pigmentation. Biology Open 10, 056523 (2021) 10.1242/bio.056523

[41] Beńıtez-Burraco, A., Boeckx, C.: Possible functional links among brain- and skull-related genes selected in modern humans. Frontiers in Psychology 6(June), 794 (2015) 10.3389/fpsyg.2015.00794

[42] Kuhlwilm, M., Boeckx, C.: A catalog of single nucleotide changes distinguishing modern humans from archaic hominins. Scientific Reports 9(1), 8463 (2019) 10.1038/s41598-019-44877-x

[43] Ellingford, J.M., Barton, S., Bhaskar, S., O9Sullivan, J., Williams, S.G., Lamb, J.A., Panda, B., Sergouniotis, P.I., Gillespie, R.L., Daiger, S.P., Hall, G., Gale, T., Lloyd, I.C., Bishop, P.N., Ramsden, S.C., Black, G.C.M.: Molecular findings from 537 individuals with inherited retinal disease. Journal of Medical Genetics 53(11), 761–767 (2016) 10.1136/jmedgenet-2016-103837

[44] Deng, L., Pan, Y., Wang, Y., Chen, H., Yuan, K., Chen, S., Lu, D., Lu, Y., Mokhtar, S.S., Rahman, T.A., Hoh, B.-P., Xu, S.: Genetic Connections and Convergent Evolution of Tropical Indigenous Peoples in Asia. Molecular Biology and Evolution 39(2), 361 (2022) 10.1093/molbev/msab361

[45] Gelabert, P., Bickle, P., Hofmann, D., Teschler-Nicola, M., Anders, A., Huang, X., Hämmerle, M., Olalde, I., Fournier, R., Ringbauer, H., Akbari, A., Cheronet, O., Lazaridis, I., Broomandkhoshbacht, N., Fernandes, D.M., Buttinger, K., Callan, K., Candilio, F., Bravo Morante, G., Curtis, E., Ferry, M., Keating, D., Freilich, S., Kearns, A., Harney, ., Lawson, A.M., Mandl, K., Michel, M., Oberreiter, V., Zagorc, B., Oppenheimer, J., Sawyer, S., Schattke, C., Özdŏgan, K.T., Qiu, L., Workman, J.N., Zalzala, F., Mallick, S., Mah, M., Micco, A., Pieler, F., Pavuk, J., Šef̌ćaková, A., Lazar, C., Starovíc, A., Djuric, M., Krznaríc Škrivanko, M., Šlaus, M., Bedić, ., Novotny, F. D. Szabó, L., Cserpák-Laczi, O., Hága, T., Szolnoki, L., Hajdú, Z., Mirea, P., Nagy, E.G., Viŕag, Z.M., Horváth M.A., Horváth, L.A.T. Biŕo, K., Domboŕoczki, L., Szeniczey, T., Jakucs, J., Szelekovszky, M., Zoltán, F., Sztáncsuj, S.J., Tóth, K., Csengeri, P., Pap, I., Patay, R., Putica, A., Vasov, B., Havasi, B., Sebők, K., Raczky, P., Lovász, G., Tvrdý, Z., Rohland, N., Novak, M., Ruttkay, M., Krǒsĺaková, M., Bátora, J., Paluch, T., Boríc, D., Dani, J., Kuhlwilm, M., Palamara, P.F., Hajdu, T., Pinhasi, R., Reich, D.: Social and genetic diversity in first farmers of central Europe. Nature Human Behaviour (2024) 10.1038/s41562-024-02034-z

[46] Herzog, T., Larena, M., Kutanan, W., Lukas, H., Fieder, M., Schaschl, H.: Natural selection and adaptive traits in the Maniq, a nomadic hunter-gatherer society from Mainland Southeast Asia. Scientific Reports 15(1), 4809 (2025) 10.1038/s41598-024-83657-0

[47] Genovese, S., Bazzigaluppi, E., Gonçalves, D., Ciucci, A., Cavallo, M.G., Purrello, F., Anello, M., Rotella, C.M., Bardini, G., Vaccaro, O., Riccardi, G., Travaglini, P., Morenghi, E., Bosi, E., Pozzilli, P.: Clinical phenotype and *β*-cell autoimmunity in Italian patients with adult-onset diabetes. European Journal of Endocrinology 154(3), 441–447 (2006) 10.1530/eje.1.02115

[48] Südhof, T.C.: Synaptic Neurexin Complexes: A Molecular Code for the Logic of Neural Circuits. Cell 171(4), 745–769 (2017) 10.1016/j.cell.2017.10.024

[49] Waal, F.B.M.D.: The Communicative Repertoire of Captive Bonobos (Pan Paniscus), Compared To That of Chimpanzees. Behaviour 106(3-4), 183–251 (1988) 10.1163/156853988X00269

[50] Stoessel, A., David, R., Bornitz, M., Ossmann, S., Neudert, M.: Auditory thresholds compatible with optimal speech reception likely evolved before the human-chimpanzee split. Scientific Reports 13(1), 20732 (2023) 10.1038/s41598-023-47778-2

[51] Ficarelli, M., Wilson, H., Pedro Galão, R., Mazzon, M., Antzin-Anduetza, I., Marsh, M., Neil, S.J.D., Swanson, C.M.: KHNYN is essential for the zinc finger antiviral protein (ZAP) to restrict HIV-1 containing clustered CpG dinucleotides. eLife 8, 46767 (2019) 10.7554/eLife.46767

[52] Saulle, I., Marventano, I., Saresella, M., Vanetti, C., Garziano, M., Fenizia, C., Trabattoni, D., Clerici, M., Biasin, M.: ERAPs Reduce In Vitro HIV Infection by Activating Innate Immune Response. The Journal of Immunology 206(7), 1609–1617 (2021) 10.4049/jimmunol.2000991

[53] Feng, G.-J., Xu, Q., Zhao, Q.-G., Han, B.-X., Yan, S.-S., Zhu, J., Pei, Y.-F.: The genetic architecture of age at menarche and its causal effects on other traits. Journal of Human Genetics 69(12), 645–653 (2024) 10.1038/s10038-024-01287-w

[54] Hollis, B., Day, F.R., Busch, A.S., Thompson, D.J., Soares, A.L.G., Timmers, P.R.H.J., Kwong, A., Easton, D.F., Joshi, P.K., Timpson, N.J., Eeles, R.A., Henderson, B.E., Haiman, C.A., Kote-Jarai, Z., Schumacher, F.R., Olama, A.A.A., Benlloch, S., Muir, K., Berndt, S.I., Conti, D.V., Wiklund, F., Chanock, S., Gapstur, S., Stevens, V.L., Tangen, C.M., Batra, J., Clements, J., Tilley, W., Risbridger, G.P., Clements, J., Horvath, L., Taylor, R., Hayes, V., Butler, L., Yeadon, T., Eckert, A., Saunders, P., Haynes, A.-M., Papargiris, M., Srinivasan, S., Kedda, M.-A., Moya, L., Batra, J., Gronberg, H., Pashayan, N., Schleutker, J., Albanes, D., Wolk, A., West, C., Mucci, L., Cancel-Tassin, G., Koutros, S., Sorensen, K.D., Grindedal, E.M., Neal, D.E., Hamdy, F.C., Donovan, J.L., Travis, R.C., Hamilton, R.J., Ingles, S.A., Rosenstein, B.S., Lu, Y.-J., Giles, G.G., Kibel, A.S., Vega, A., Kogevinas, M., Penney, K.L., Park, J.Y., Stanford, J.L., Cybulski, C., Nordestgaard, B.G., Brenner, H., Maier, C., Kim, J., John, E.M., Teixeira, M.R., Neuhausen, S.L., De Ruyck, K., Razack, A., Newcomb, L.F., Lessel, D., Kaneva, R., Usmani, N., Claessens, F., Townsend, P.A., Gago-Dominguez, M., Roobol, M.J., Menegaux, F., Khaw, K.-T., Cannon-Albright, L., Pandha, H., Thibodeau, S.N., Agee, M., Alipanahi, B., Auton, A., Bell, R.K., Bryc, K., Elson, S.L., Fontanillas, P., Furlotte, N.A., Hinds, D.A., Huber, K.E., Kleinman, A., Litterman, N.K., McIntyre, M.H., Mountain, J.L., Noblin, E.S., Northover, C.A.M., Pitts, S.J., Sathirapongsasuti, J.F., Sazonova, O.V., Shelton, J.F., Shringarpure, S., Tian, C., Tung, J.Y., Vacic, V., Wilson, C.H., Ong, K.K., Perry, J.R.B., Consortium, T.P., (APCB), A.P.C.B., Team, a.R.: Genomic analysis of male puberty timing highlights shared genetic basis with hair colour and lifespan. Nature Communications 11(1), 1536 (2020) 10.1038/s41467-020-14451-5

[55] Kushima, I., Aleksic, B., Nakatochi, M., Shimamura, T., Okada, T., Uno, Y., Morikawa, M., Ishizuka, K., Shiino, T., Kimura, H., Arioka, Y., Yoshimi, A., Takasaki, Y., Yu, Y., Nakamura, Y., Yamamoto, M., Iidaka, T., Iritani, S., Inada, T., Ogawa, N., Shishido, E., Torii, Y., Kawano, N., Omura, Y., Yoshikawa, T., Uchiyama, T., Yamamoto, T., Ikeda, M., Hashimoto, R., Yamamori, H., Yasuda, Y., Someya, T., Watanabe, Y., Egawa, J., Nunokawa, A., Itokawa, M., Arai, M., Miyashita, M., Kobori, A., Suzuki, M., Takahashi, T., Usami, M., Kodaira, M., Watanabe, K., Sasaki, T., Kuwabara, H., Tochigi, M., Nishimura, F., Yamasue, H., Eriguchi, Y., Benner, S., Kojima, M., Yassin, W., Munesue, T., Yokoyama, S., Kimura, R., Funabiki, Y., Kosaka, H., Ishitobi, M., Ohmori, T., Numata, S., Yoshikawa, T., Toyota, T., Yamakawa, K., Suzuki, T., Inoue, Y., Nakaoka, K., Goto, Y.-i., Inagaki, M., Hashimoto, N., Kusumi, I., Son, S., Murai, T., Ikegame, T., Okada, N., Kasai, K., Kunimoto, S., Mori, D., Iwata, N., Ozaki, N.: Comparative Analyses of Copy-Number Variation in Autism Spectrum Disorder and Schizophrenia Reveal Etiological Overlap and Biological Insights. Cell Reports 24(11), 2838–2856 (2018) 10.1016/j.celrep.2018.08.022

[56] Blount, B.G.: Issues in Bonobo (Pan paniscus) Sexual Behavior. American Anthropologist 92(3), 702–714 (1990) 10.1525/aa.1990.92.3.02a00100

[57] Dahl, J.F.: The external genitalia of female pygmy chimpanzees. The Anatomical Record 211(1), 24–28 (1985) 10.1002/ar.1092110105

[58] Pourová, R., Janoúsek, P., Jurov̌ćık, M., Dvǒŕaková, M., Maĺıková, M., Rásková, D., Bendová, O., Leonardi, E., Murgia, A., Kabelka, Z., Astl, J., Seeman, P.: Spectrum and Frequency of SLC26A4 Mutations Among Czech Patients with Early Hearing Loss with and without Enlarged Vestibular Aqueduct (EVA). Annals of Human Genetics 74(4), 299–307 (2010) 10.1111/j.1469-1809.2010.00581.x

[59] Bertrand, M.J.M., Doiron, K., Labbé, K., Korneluk, R.G., Barker, P.A., Saleh, M.: Cellular Inhibitors of Apoptosis cIAP1 and cIAP2 Are Required for Innate Immunity Signaling by the Pattern Recognition Receptors NOD1 and NOD2. Immunity 30(6), 789–801 (2009) 10.1016/j.immuni.2009.04.011

[60] Ryan, E.J., Marshall, A.J., Magaletti, D., Floyd, H., Draves, K.E., Olson, N.E., Clark, E.A.: Dendritic Cell-Associated Lectin-1: A Novel Dendritic Cell-Associated, C-Type Lectin-Like Molecule Enhances T Cell Secretion of IL-41. The Journal of Immunology 169(10), 5638–5648 (2002) 10.4049/jimmunol.169.10.5638

[61] Hill, M.R., Cookson, W.O.C.M.: A New Variant of the *β* Subunit of the High-Affinity Receptor for Immunoglobulin E (Fc*ɛ*RI-*β* E237G): Associations with Measures of Atopy and Bronchial Hyper-Responsiveness. Human Molecular Genetics 5(7), 959–962 (1996) 10.1093/hmg/5.7.959

[62] Tsukada, Y.-i., Ishitani, T., Nakayama, K.I.: KDM7 is a dual demethylase for histone H3 Lys 9 and Lys 27 and functions in brain development. Genes & Development 24(5), 432–437 (2010) 10.1101/gad.1864410

[63] Kleylein-Sohn, J., Westendorf, J., Le Clech, M., Habedanck, R., Stierhof, Y.-D., Nigg, E.A.: Plk4-Induced Centriole Biogenesis in Human Cells. Developmental Cell 13(2), 190–202 (2007) 10.1016/j.devcel.2007.07.002

[64] Carroll, C.W., Silva, M.C.C., Godek, K.M., Jansen, L.E.T., Straight, A.F.: Centromere assembly requires the direct recognition of CENP-A nucleosomes by CENP-N. Nature Cell Biology 11(7), 896–902 (2009) 10.1038/ncb1899

[65] Lan, K.-C., Cheng, Y.-H., Chang, Y.-C., Wei, K.-T., Weng, P.-L., Kang, H.-Y.: Interaction between Chromodomain Y-like Protein and Androgen Receptor Signaling in Sertoli Cells Accounts for Spermatogenesis (2024). 10.3390/cells13100851

[66] Haenig, C., Atias, N., Taylor, A.K., Mazza, A., Schaefer, M.H., Russ, J., Riechers, S.-P., Jain, S., Coughlin, M., Fontaine, J.-F., Freibaum, B.D., Brusendorf, L., Zenkner, M., Porras, P., Stroedicke, M., Schnoegl, S., Arnsburg, K., Boeddrich, A., Pigazzini, L., Heutink, P., Taylor, J.P., Kirstein, J., Andrade-Navarro, M.A., Sharan, R., Wanker, E.E.: Interactome Mapping Provides a Network of Neurodegenerative Disease Proteins and Uncovers Widespread Protein Aggregation in Affected Brains. Cell Reports 32(7) (2020) 10.1016/j.celrep.2020.108050

[67] Dai, T., Wu, L., Wang, S., Wang, J., Xie, F., Zhang, Z., Fang, X., Li, J., Fang, P., Li, F., Jin, K., Dai, J., Yang, B., Zhou, F., Dam, H., Cai, D., Huang, H., Zhang, L.: FAF1 Regulates Antiviral Immunity by Inhibiting MAVS but Is Antagonized by Phosphorylation upon Viral Infection. Cell Host & Microbe 24(6), 776–790 (2018) 10.1016/j.chom.2018.10.006

[68] Sikora, M., Ferrer-Admetlla, A., Laayouni, H., Menendez, C., Mayor, A., Bardaji, A., Sigauque, B., Mandomando, I., Alonso, P.L., Bertranpetit, J., Casals, F.: A variant in the gene FUT9 is associated with susceptibility to placental malaria infection. Human Molecular Genetics 18(16), 3136–3144 (2009) 10.1093/hmg/ddp240

[69] Kurogi, K., Shimohira, T., Kouriki-Nagatomo, H., Zhang, G., Miller, E.R., Sakakibara, Y., Suiko, M., Liu, M.-C.: Human Cytosolic Sulphotransferase SULT1C3: genomic analysis and functional characterization of splice variant SULT1C3a and SULT1C3d. The Journal of Biochemistry 162(6), 403–414 (2017) 10.1093/jb/mvx044

[70] Fontsere, C., Kuhlwilm, M., Morcillo-Suarez, C., Alvarez-Estape, M., Lester, J.D., Gratton, P., Schmidt, J.M., Dieguez, P., Aebischer, T., Álvarez-Varona, P., Agbor, A., Angedakin, S., Assumang, A.K., Ayimisin, E.A., Bailey, E., Barubiyo, D., Bessone, M., Carretero-Alonso, A., Chancellor, R., Cohen, H., Danquah, E., Deschner, T., Dunn, A., Dupain, J., Egbe, V.E., Feliu, O., Goedmakers, A., Granjon, A.-C., Head, J., Hedwig, D., Hermans, V., Hernandez-Aguilar, R.A., Imong, I., Jones, S., Junker, J., Kadam, P., Kaiser, M., Kambere, M., Kambale, M.V., Kalan, A.K., Kienast, I., Kujirakwinja, D., Langergraber, K., Lapuente, J., Larson, B., Laudisoit, A., Lee, K., Llana, M., Llorente, M., Marrocoli, S., Morgan, D., Mulindahabi, F., Murai, M., Neil, E., Nicholl, S., Nixon, S., Normand, E., Orbell, C., Ormsby, L.J., Pacheco, L., Piel, A., Riera, L., Robbins, M.M., Rundus, A., Sanz, C., Sciaky, L., Sommer, V., Stewart, F.A., Tagg, N., Tédonzong, L.R., Ton, E., Schijndel, J., Vergnes, V., Wessling, E.G., Willie, J., Wittig, R.M., Yuh, Y.G., Yurkiw, K., Zuberbuehler, K., Hecht, J., Vigilant, L., Boesch, C., Andŕes, A.M., Hughes, D.A., Kühl, H.S., Lizano, E., Arandjelovic, M., Marques-Bonet, T.: Population dynamics and genetic connectivity in recent chimpanzee history. Cell Genomics 2(6) (2022) 10.1016/j.xgen.2022.100133

[71] Ruotsalainen, V., Ljungberg, P., Wartiovaara, J., Lenkkeri, U., Kestilä, M., Jalanko, H., Holmberg, C., Tryggvason, K.: Nephrin is specifically located at the slit diaphragm of glomerular podocytes. Proceedings of the National Academy of Sciences 96(14), 7962–7967 (1999) 10.1073/pnas.96.14.7962

[72] Niki, T., Takahashi-Niki, K., Taira, T., Iguchi-Ariga, S.M.M., Ariga, H.: DJBP: A Novel DJ-1-Binding Protein, Negatively Regulates the Androgen Receptor by Recruiting Histone Deacetylase Complex, and DJ-1 Antagonizes This Inhibition by Abrogation of This Complex. Molecular Cancer Research 1(4), 247–261 (2003)

[73] Anderson, M.J., Dixson, A.F.: Motility and the midpiece in primates. Nature 416(6880), 496 (2002) 10.1038/416496a

[74] Wang, W., Meng, L., He, J., Su, L., Li, Y., Tan, C., Xu, X., Nie, H., Zhang, H., Du, J., Lu, G., Luo, M., Lin, G., Tu, C., Tan, Y.-Q.: Bi-allelic variants in SHOC1 cause non-obstructive azoospermia with meiosis arrest in humans and mice. Molecular Human Reproduction 28(6), 015 (2022) https://doi.org/10.1093/molehr/gaac015

[75] Shi, B., Shah, W., Liu, L., Gong, C., Zhou, J., Abbas, T., Ma, H., Zhang, H., Yang, M., Zhang, Y., Ullah, N., Mahammad, Z., Khan, M., Murtaza, G., Ali, A., Khan, R., Sha, J., Yuan, Y., Shi, Q.: Biallelic mutations in RNA-binding protein ADAD2 cause spermiogenic failure and non-obstructive azoospermia in humans. Human Reproduction Open 2023(3), 022 (2023) 10.1093/hropen/hoad022

[76] Dacheux, D., Martinez, G., Broster Reix, C.E., Beurois, J., Lores, P., Tounkara, M., Dupuy, J.-W., Robinson, D.R., Loeuillet, C., Lambert, E., Wehbe, Z., Escoffier, J., Amiri-Yekta, A., Daneshipour, A., Hosseini, S.-H., Zouari, R., Mustapha, S.F.B., Halouani, L., Jiang, X., Shen, Y., Liu, C., Thierry-Mieg, N., Septier, A., Bidart, M., Satre, V., Cazin, C., Kherraf, Z.E., Arnoult, C., Ray, P.F., Toure, A., Bonhivers, M., Coutton, C.: Novel axonemal protein ZMYND12 interacts with TTC29 and DNAH1, and is required for male fertility and flagellum function. eLife 12, 87698 (2023) 10.7554/eLife.87698

[77] Meng, Q., Shao, B., Zhao, D., Fu, X., Wang, J., Li, H., Zhou, Q., Gao, T.: Loss of SUN1 function in spermatocytes disrupts the attachment of telomeres to the nuclear envelope and contributes to non-obstructive azoospermia in humans. Human Genetics 142(4), 531–541 (2023) 10.1007/s00439-022-02515-z

[78] Okada, M., Cheeseman, I.M., Hori, T., Okawa, K., McLeod, I.X., Yates, J.R., Desai, A., Fukagawa, T.: The CENP-HI complex is required for the efficient incorporation of newly synthesized CENP-A into centromeres. Nature Cell Biology 8(5), 446–457 (2006) 10.1038/ncb1396

[79] Cui, Y., Wang, W., Dong, N., Lou, J., Srinivasan, D.K., Cheng, W., Huang, X., Liu, M., Fang, C., Peng, J., Chen, S., Wu, S., Liu, Z., Dong, L., Zhou, Y., Wu, Q.: Role of corin in trophoblast invasion and uterine spiral artery remodelling in pregnancy. Nature 484(7393), 246–250 (2012) 10.1038/nature10897

[80] Ballatori, N., Christian, W.V., Lee, J.Y., Dawson, P.A., Soroka, C.J., Boyer, J.L., Madejczyk, M.S., Li, N.: OST*α*OST*β*: A major basolateral bile acid and steroid transporter in human intestinal, renal, and biliary epithelia. Hepatology 42(6) (2005) 10.1002/hep.20961

[81] Beurois, J., Martinez, G., Cazin, C., Kherraf, Z.-E., Amiri-Yekta, A., Thierry-Mieg, N., Bidart, M., Petre, G., Satre, V., Brouillet, S., Touŕe, A., Arnoult, C., Ray, P.F., Coutton, C.: CFAP70 mutations lead to male infertility due to severe astheno-teratozoospermia. A case report. Human Reproduction 34(10), 2071–2079 (2019) 10.1093/humrep/dez166

[82] Sarkardeh, H., Totonchi, M., Asadpour, O., Sadighi Gilani, M.A., Zamani Esteki, M., Almadani, N., Borjian Boroujeni, P., Gourabi, H.: Association of MOV10L1 gene polymorphisms and male infertility in azoospermic men with complete maturation arrest. Journal of Assisted Reproduction and Genetics 31(7), 865– 871 (2014) 10.1007/s10815-014-0240-1

[83] Mourelatos, Z., Gonatas, J.O., Cinato, E., Gonatas, N.K.: Cloning and Sequence Analysis of the Human MG160, a Fibroblast Growth Factor and E-Selectin Binding Membrane Sialoglycoprotein of the Golgi Apparatus. DNA and Cell Biology 15(12), 1121–1128 (1996) 10.1089/dna.1996.15.1121

[84] Akinwumi, B.C., Bordun, K.-A.M., Anderson, H.D.: Biological Activities of Stilbenoids. International Journal of Molecular Sciences 19(3) (2018) 10.3390/ijms19030792

[85] Elizabeth Rogers, M., Maisels, F., Williamson, E.A., Fernandez, M., Tutin, C.E.G.: Gorilla diet in the Lopé Reserve, Gabon:. Oecologia 84(3), 326–339 (1990) 10.1007/BF00329756

[86] Dan-zeng, Z.-g., Bai-ma, Y.-j., Huang, J., Suo-lang, W.-j., Ge-sang, Q.-z., Suona, Y.-z., Wang, Y.-s., Baima, Z.-g., Ge-sang, l.-b.: Integrative proteomics and metabolomics reveal important pathways and potential biomarkers in high-altitude pulmonary hypertension. Scientific Reports 15(1), 24999 (2025) 10.1038/s41598-025-09477-y

[87] Guo, B.-X., Zhang, Y., Sun, X.-Y., Sun, Y.-X., Lv, W.-J., Xu, S.-X., Yang, G., Ren, W.-H.: Convergent evolution in high-altitude and marine mammals: Molecular adaptations to pulmonary fibrosis and hypoxia . Zoological Research 45(6), 1209–1220 (2024) 10.24272/j.issn.2095-8137.2024.029

[88] Verheij, J.B.G.M., Kunze, J., Osinga, J., Essen, A.J., Hofstra, R.M.W.: ABCD syndrome is caused by a homozygous mutation in the EDNRB gene. American Journal of Medical Genetics 108(3), 223–225 (2002) 10.1002/ajmg.10172

[89] Waselle, L., Coppola, T., Fukuda, M., Iezzi, M., El-Amraoui, A., Petit, C., Regazzi, R.: Involvement of the Rab27 Binding Protein Slac2c/MyRIP in Insulin Exocytosis. Molecular Biology of the Cell 14(10), 4103–4113 (2003) 10.1091/mbc.e03-01-0022

[90] Ge, R.-L., Simonson, T.S., Gordeuk, V., Prchal, J.T., McClain, D.A.: Metabolic aspects of high-altitude adaptation in Tibetans. Experimental Physiology 100(11), 1247–1255 (2015) 10.1113/EP085292

[91] Noordwijk, M.A., Arora, N., Willems, E.P., Dunkel, L.P., Amda, R.N., Mardianah, N., Ackermann, C., Krützen, M., Schaik, C.P.: Female philopatry and its social benefits among Bornean orangutans. Behavioral Ecology and Sociobiology 66(6), 823–834 (2012) 10.1007/s00265-012-1330-7

[92] Mörchen, J., Luhn, F., Wassmer, O., Kunz, J.A., Kulik, L., Noordwijk, M.A., Rianti, P., Rahmaeti, T., Utami Atmoko, S.S., Widdig, A., Schuppli, C.: Orangutan males make increased use of social learning opportunities, when resource availability is high. iScience 27(2) (2024) 10.1016/j.isci.2024.108940

[93] Courtenay, J.: Inter- or Intra-Island Variation? An Assessment of the Differences Between Bornean and Sumatran Orang-Utans. Orangutan Biology, 19–29 (1988)

[94] Knott, C.D.: Changes in Orangutan Caloric Intake, Energy Balance, and Ketones in Response to Fluctuating Fruit Availability. International Journal of Primatology 19(6), 1061–1079 (1998) 10.1023/A:1020330404983

[95] Gleeson, P.A.: The role of endosomes in innate and adaptive immunity. Seminars in Cell & Developmental Biology 31, 64–72 (2014) 10.1016/j.semcdb.2014.03.002

[96] Besnard, M.-L., Raymond-Letron, I., Jourdan, G., Becker, C., Steinmetz, H.W.: MORBIDITY AND MORTALITY REVIEW OF THE CAPTIVE EUROPEAN ASSOCIATION OF ZOOS AND AQUARIA EX-SITU PROGRAMME ORANGUTAN (¡i¿PONGO¡/i¿ SPECIES) POPULATION BETWEEN 2000 AND 2018. Journal of Zoo and Wildlife Medicine 56(2), 237–247 (2025) 10.1638/2022-0137

[97] Andŕes, A.M., Hubisz, M.J., Indap, A., Torgerson, D.G., Degenhardt, J.D., Boyko, A.R., Gutenkunst, R.N., White, T.J., Green, E.D., Bustamante, C.D., Clark, A.G., Nielsen, R.: Targets of Balancing Selection in the Human Genome. Molecular Biology and Evolution 26(12), 2755–2764 (2009) 10.1093/molbev/msp190

[98] Dong, S., Zhang, B., Huang, K., Ying, M., Yan, J., Niu, F., Hu, H., Dunn, D.W., Ren, Y., Li, B., Zhang, P.: Balancing selection shapes population differentiation of major histocompatibility complex genes in wild golden snub-nosed monkeys. Current Zoology 70(5), 596–606 (2024) 10.1093/cz/zoad043

[99] Hedrick, P.W.: Balancing selection and MHC. Genetica 104(3), 207–214 (1998) 10.1023/A:1026494212540

[100] Jones, K.E., Patel, N.G., Levy, M.A., Storeygard, A., Balk, D., Gittleman, J.L., Daszak, P.: Global trends in emerging infectious diseases. Nature 451(7181), 990–993 (2008) 10.1038/nature06536

[101] Lee, A.S., Rusch, J., Lima, A.C., Usmani, A., Huang, N., Lepamets, M., Vigh-Conrad, K.A., Worthington, R.E., Mägi, R., Wu, X., Aston, K.I., Atkinson, J.P., Carrell, D.T., Hess, R.A., OBryan, M.K., Conrad, D.F.: Rare mutations in the complement regulatory gene CSMD1 are associated with male and female infertility. Nature Communications 10(1), 4626 (2019) 10.1038/s41467-019-12522-w

[102] Zhang, Q., Xing, M., Bao, Z., Xu, L., Bai, Y., Chen, W., Pan, W., Cai, F., Wang, Q., Guo, S., Zhang, J., Wang, Z., Wu, Y., Zhang, Y., Li, J.-D., Song, W.: Contactin-associated protein-like 2 (CNTNAP2) mutations impair the essential *α*-secretase cleavages, leading to autism-like phenotypes. Signal Transduction and Targeted Therapy 9(1), 51 (2024) 10.1038/s41392-024-01768-6

[103] Kim, B.Y., Huber, C.D., Lohmueller, K.E.: Inference of the distribution of selection coefficients for new nonsynonymous mutations using large samples. Genetics 206, 345–361 (2017) genetics.116.197145

[104] Alvarez-Estape, M., Pawar, H., Fontsere, C., Trujillo, A.E., Gunson, J.L., Bergl, R.A., Bermejo, M., Linder, J.M., McFarland, K., Oates, J.F., Sunderland-Groves, J.L., Orkin, J., Higham, J.P., Viaud-Martinez, K.A., Lizano, E., Marques-Bonet, T.: Past Connectivity but Recent Inbreeding in Cross River Gorillas Determined Using Whole Genomes from Single Hairs (2023). 10.3390/genes14030743

[105] Makova, K.D., Pickett, B.D., Harris, R.S., Hartley, G.A., Cechova, M., Pal, K., Nurk, S., Yoo, D., Li, Q., Hebbar, P., McGrath, B.C., Antonacci, F., Aubel, M., Biddanda, A., Borchers, M., Bornberg-Bauer, E., Bouffard, G.G., Brooks, S.Y., Carbone, L., Carrel, L., Carroll, A., Chang, P.-C., Chin, C.-S., Cook, D.E., Craig, S.J.C., Gennaro, L., Diekhans, M., Dutra, A., Garcia, G.H., Grady, P.G.S., Green, R.E., Haddad, D., Hallast, P., Harvey, W.T., Hickey, G., Hillis, D.A., Hoyt, S.J., Jeong, H., Kamali, K., Pond, S.L.K., LaPolice, T.M., Lee, C., Lewis, A.P., Loh, Y.-H.E., Masterson, P., McGarvey, K.M., McCoy, R.C., Medvedev, P., Miga, K.H., Munson, K.M., Pak, E., Paten, B., Pinto, B.J., Potapova, T., Rhie, A., Rocha, J.L., Ryabov, F., Ryder, O.A., Sacco, S., Shafin, K., Shepelev, V.A., Slon, V., Solar, S.J., Storer, J.M., Sudmant, P.H., Sweetalana, Sweeten A., Tassia, M.G., Thibaud-Nissen, F., Ventura, M., Wilson, M.A., Young, A.C., Zeng, H., Zhang, X., Szpiech, Z.A., Huber, C.D., dGerton, J.L., Yi, S.V., Schatz, M.C., Alexandrov, I.A., Koren, S., ONeill, R.J., Eichler, E.E., Phillippy, A.M.: The complete sequence and comparative analysis of ape sex chromosomes. Nature (2024) 10.1038/s41586-024-07473-2

[106] Venn, O., Turner, I., Mathieson, I., Groot, N., Bontrop, R., McVean, G.: Strong male bias drives germline mutation in chimpanzees. Science 344(6189), 1272–1275 (2014) 10.1126/science.344.6189.1272

[107] Martinez, G., Garcia, C.: Sexual selection and sperm diversity in primates. Molecular and Cellular Endocrinology 518, 110974 (2020) 10.1016/j.mce.2020.110974

[108] Cawthon Lang, K.: Primate Factsheets: Gorilla (Gorilla) Taxonomy, Morphology, & Ecology (0). http://pin.primate.wisc.edu/factsheets/entry/gorilla/taxon

[109] Furuichi, T.: Female contributions to the peaceful nature of bonobo society. Evolutionary Anthropology: Issues, News, and Reviews 20(4), 131–142 (2011) 10.1002/evan.20308

[110] Keightley, P.D., Eyre-Walker, A.: Joint Inference of the Distribution of Fitness Effects of Deleterious Mutations and Population Demography Based on Nucleotide Polymorphism Frequencies . Genetics 177, 2251–2261 (2007) 10.1534/genetics.107.080663

[111] Tataru, P., Mollion, M., Glmin, S., Bataillon, T.: Inference of Distribution of Fitness Effects and Proportion of Adaptive Substitutions from Polymorphism Data . Genetics 207, 1103–1119 (2017) 10.1534/genetics.117.300323

[112] Huang, X., Rymbekova, A., Dolgova, O., Lao, O., Kuhlwilm, M.: Harnessing deep learning for population genetic inference. Nature Reviews Genetics 25, 61–78 (2024) 10.1038/s41576-023-00636-3

[113] Huang, X., Hackl, J., Kuhlwilm, M.: Decoding genomic landscapes of introgression. Trends in Genetics (2025) 10.1016/j.tig.2025.07.001

[114] Kuderna, L.F.K., Gao, H., Janiak, M.C., Kuhlwilm, M., Orkin, J.D., Bataillon, T., Manu, S., Valenzuela, A., Bergman, J., Rousselle, M., Silva, F.E., Agueda, L., Blanc, J., Gut, M., Vries, D., Goodhead, I., Harris, R.A., Raveendran, M., Jensen, A., Chuma, I.S., Horvath, J.E., Hvilsom, C., Juan, D., Frandsen, P., Schraiber, J.G., Melo, F.R., Bertuol, F., Byrne, H., Sampaio, I., Farias, I., Valsecchi, J., Messias, M., Silva, M.N.F., Trivedi, M., Rossi, R., Hrbek, T., Andriaholinirina, N., Rabarivola, C.J., Zaramody, A., Jolly, C.J., Phillips-Conroy, J., Wilkerson, G., Abee, C., Simmons, J.H., Fernandez-Duque, E., Kanthaswamy, S., Shiferaw, F., Wu, D., Zhou, L., Shao, Y., Zhang, G., Keyyu, J.D., Knauf, S., Le, M.D., Lizano, E., Merker, S., Navarro, A., Nadler, T., Khor, C.C., Lee, J., Tan, P., Lim, W.K., Kitchener, A.C., Zinner, D., Gut, I., Melin, A.D., Guschanski, K., Schierup, M.H., Beck, R.M.D., Umapathy, G., Roos, C., Boubli, J.P., Rogers, J., Farh, K.K.-H., Marques Bonet, T.: A global catalog of whole-genome diversity from 233 primate species. Science 380(6648), 906–913 (2023) 10.1126/science.abn7829

[115] Sabeti, P.C., Varilly, P., Fry, B., Lohmueller, J., Hostetter, E., Cotsapas, C., al.: Genome-wide detection and characterization of positive selection in human populations. Nature 449, 913–8 (2007) 10.1038/nature06250

[116] Szpiech, Z.A., Novak, T.E., Bailey, N.P., Stevison, L.S.: Application of a novel haplotype-based scan for local adaptation to study high-altitude adaptation in rhesus macaques. Evol Lett 5, 408–21 (2021) 10.1002/evl3.232

[117] Siewert, K.M., Voight, B.F.: Detecting long-term balancing selection using allele frequency correlation. Molecular Biology and Evolution 34, 2996–3005 (2017) 10.1093/molbev/msx209

[118] Chang, C.C., Chow, C.C., Tellier, L.C.A.M., Vattikuti, S., Purcell, S.M., Lee, J.J.: Second-generation plink: rising to the challenge of larger and richer datasets. GigaScience 4, 13742–01500478 (2015) 10.1186/s13742-015-0047-8

[119] Wang, K., Li, M., Hakonarson, H.: Annovar: functional annotation of genetic variants from high-throughput sequencing data. Nucleic Acids Research 38, 164 (2010) 10.1093/nar/gkq603

[120] Sherry, S.T., Ward, M.H., Kholodov, M., Baker, J., Phan, L., Smigielski, E.M., Sirotkin, K.: dbSNP: The NCBI database of genetic variation. Nucleic Acids Research 29, 308–311 (2001) 10.1093/nar/29.1.308

[121] O’Leary, N.A., Wright, M.W., Brister, J.R., Ciufo, S., Haddad, D., McVeigh, R., Rajput, B., Robbertse, B., Smith-White, B., Ako-Adjei, D., Astashyn, A., Badretdin, A., Bao, Y., Blinkova, O., Brover, V., Chetvernin, V., Choi, J., Cox, E., Ermolaeva, O., Farrell, C.M., Goldfarb, T., Gupta, T., Haft, D., Hatcher, E., Hlavina, W., Joardar, V.S., Kodali, V.K., Li, W., Maglott, D., Masterson, P., McGarvey, K.M., Murphy, M.R., O’Neill, K., Pujar, S., Rangwala, S.H., Rausch, D., Riddick, L.D., Schoch, C., Shkeda, A., Storz, S.S., Sun, H., Thibaud-Nissen, F., Tolstoy, I., Tully, R.E., Vatsan, A.R., Wallin, C., Webb, D., Wu, W., Landrum, M.J., Kimchi, A., Tatusova, T., DiCuccio, M., Kitts, P., Murphy, T.D., Pruitt, K.D.: Reference sequence (RefSeq) database at NCBI: Current status, taxonomic expansion, and functional annotation. Nucleic Acids Research 44, 733–745 (2016) 10.1093/nar/gkv1189

[122] Liu, X., Li, C., Mou, C., Dong, Y., Tu, Y.: dbNSFP v4: A comprehensive database of transcript-specific functional predictions and annotations for human nonsynonymous and splice-site SNVs. Genome Medicine 12, 103 (2020) 10.1186/s13073-020-00803-9

[123] Stelzer, G., Rosen, N., Plaschkes, I., Zimmerman, S., Twik, M., Fishilevich, S., Stein, T.I., Nudel, R., Lieder, I., Mazor, Y., Kaplan, S., Dahary, D., Warshawsky, D., Guan-Golan, Y., Kohn, A., Rappaport, N., Safran, M., Lancet, D.: The GeneCards Suite: From Gene Data Mining to Disease Genome Sequence Analyses. Current Protocols in Bioinformatics 54(1), 1–1 (2016) 10.1002/cpbi.5

[124] Gutenkunst, R.N., Hernandez, R.D., Williamson, S.H., Bustamante, C.D.: Inferring the joint demographic history of multiple populations from multidimensional snp frequency data. PLoS Genetics 5, 1000695 (2009) 10.1371/journal.pgen.1000695

[125] Huang, X., Struck, T.J., Davey, S.W., Gutenkunst, R.N.: dadi-cli: automated and distributed population genetic model inference from allele frequency spectra. bioRixv, 2023–0615545182 (2023) 10.1101/2023.06.15.545182

[126] Huber, C.D., Kim, B.Y., Marsden, C.D., Lohmueller, K.E.: Determining the factors driving selective effects of new nonsynonymous mutations. Proceedings of the National Academy of Sciences of the United States of America 114, 4465– 4470 (2017) 10.1073/pnas.161950811

[127] Coffman, A.J., Hsieh, P., Gravel, S., Gutenkunst, R.N.: Computationally efficient composite likelihood statistics for demographic inference. Molecular Biology Evolution 33, 591–593 (2016) 10.1093/molbev/msv255

