## Supplementary Figures for "Genomic Landscapes of Natural Selection in Great Apes"

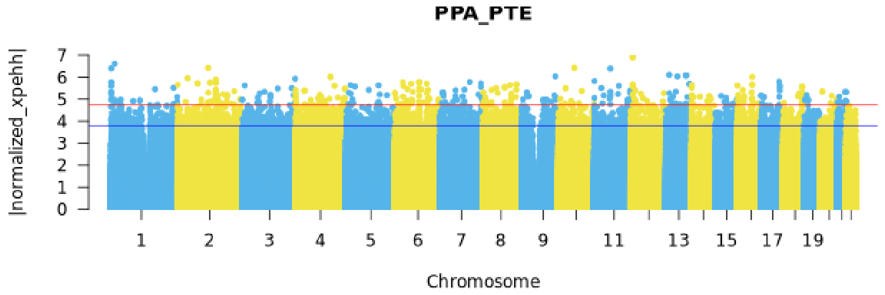


Fig. S1: Genome-wide Manhattan plot of unphased XP-EHH values for PPA vs. PTE. The blue horizontal line marks the top 0.05% cutoff, and the red horizontal line marks the top 0.005% cutoff.

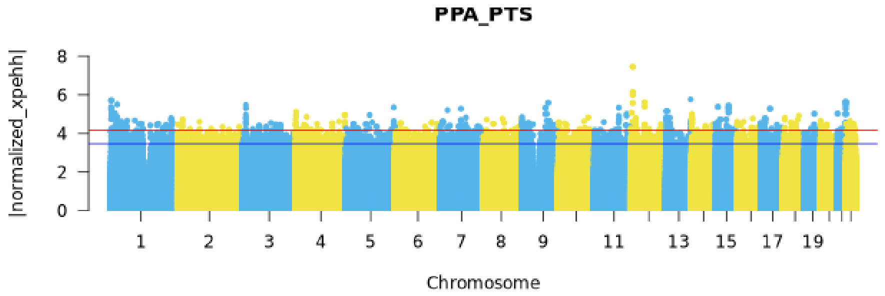


Fig. S2: Genome-wide Manhattan plot of unphased XP-EHH values for PPA vs. PTS. The blue horizontal line indicates the top 0.05% cutoff, and the red horizontal line indicates the top 0.005% cutoff.

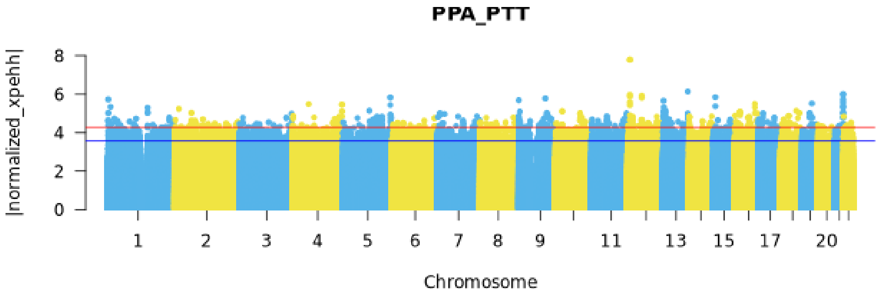


Fig. S3: Genome-wide Manhattan plot of unphased XP-EHH values for PPA vs. PTT. The blue horizontal line marks the top 0.05% cutoff, and the red horizontal line marks the top 0.005% cutoff.

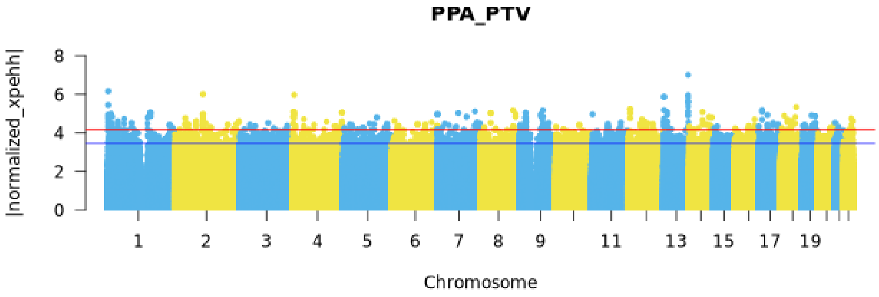


Fig. S4: Genome-wide Manhattan plot of unphased XP-EHH values for PPA vs. PTV. The blue horizontal line marks the top 0.05% cutoff, and the red horizontal line marks the top 0.005% cutoff.

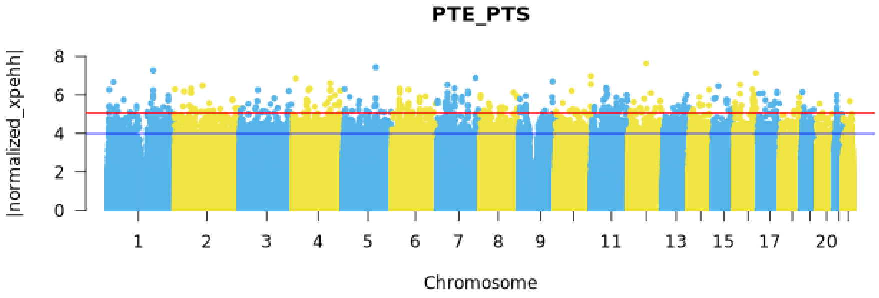


Fig. S5: Genome-wide Manhattan plot of unphased XP-EHH values for PTE vs. PTS. The blue horizontal line marks the top 0.05% cutoff, and the red horizontal line marks the top 0.005% cutoff.

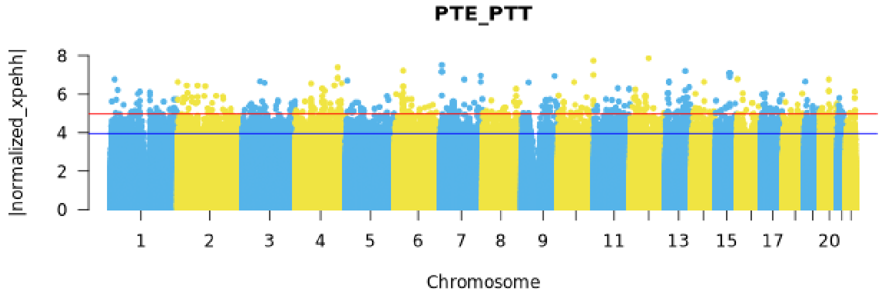


Fig. S6: Genome-wide Manhattan plot of unphased XP-EHH values for PTE vs. PTT. The blue horizontal line marks the top 0.05% cutoff, and the red horizontal line marks the top 0.005% cutoff.

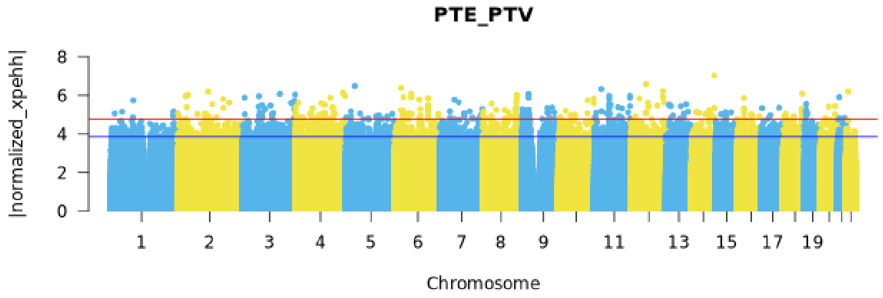


Fig. S7: Genome-wide Manhattan plot of unphased XP-EHH values for PTE vs. PTV. The blue horizontal line marks the top 0.05% cutoff, and the red horizontal line marks the top 0.005% cutoff.

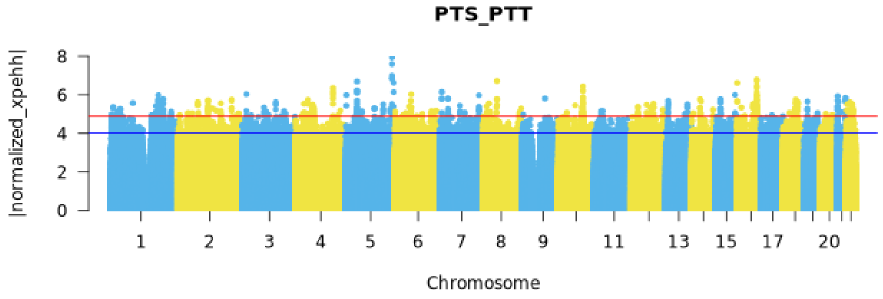


Fig. S8: Genome-wide Manhattan plot of unphased XP-EHH values for PTS vs. PTT. The blue horizontal line marks the top 0.05% cutoff, and the red horizontal line marks the top 0.005% cutoff.

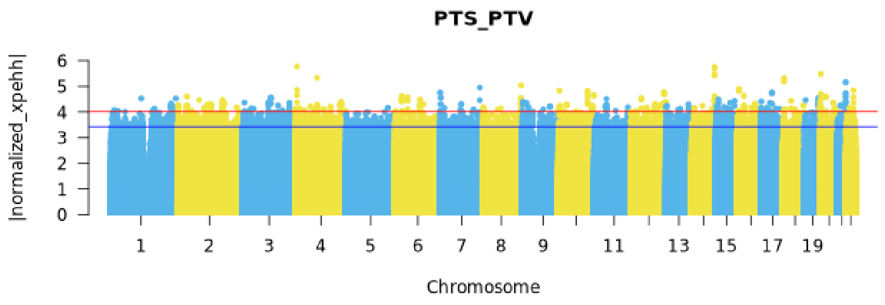


Fig. S9: Genome-wide Manhattan plot of unphased XP-EHH values for PTS vs. PTV. The blue horizontal line marks the top 0.05% cutoff, and the red horizontal line marks the top 0.005% cutoff.

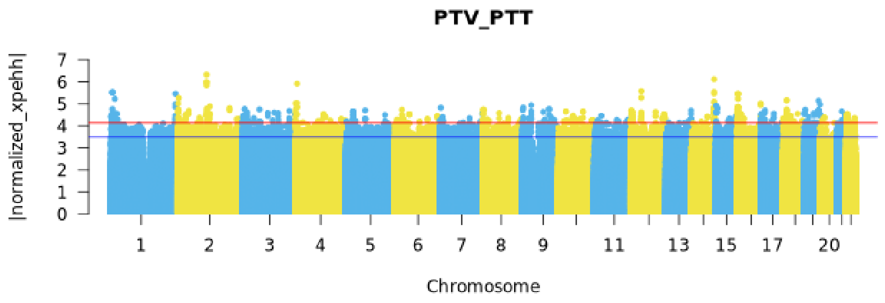


Fig. S10: Genome-wide Manhattan plot of unphased XP-EHH values for PTV vs. PTT. The blue horizontal line marks the top 0.05% cutoff, and the red horizontal line marks the top 0.005% cutoff.

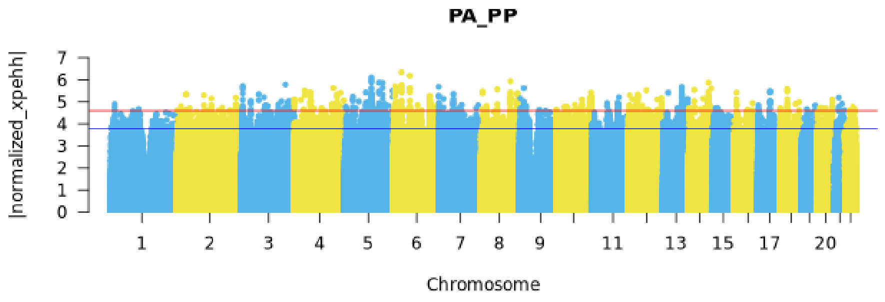


Fig. S11: Genome-wide Manhattan plot of unphased XP-EHH values for PA vs. PP. The blue horizontal line marks the top 0.05% cutoff, and the red horizontal line marks the top 0.005% cutoff.

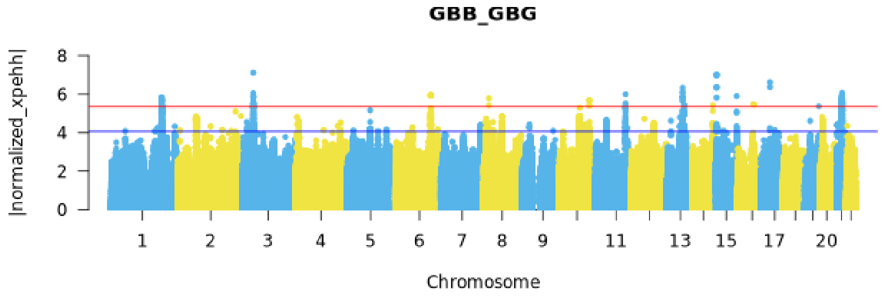


Fig. S12: Genome-wide Manhattan plot of unphased XP-EHH values for GBB vs. GBG. The blue horizontal line marks the top 0.05% cutoff, and the red horizontal line marks the top 0.005% cutoff.

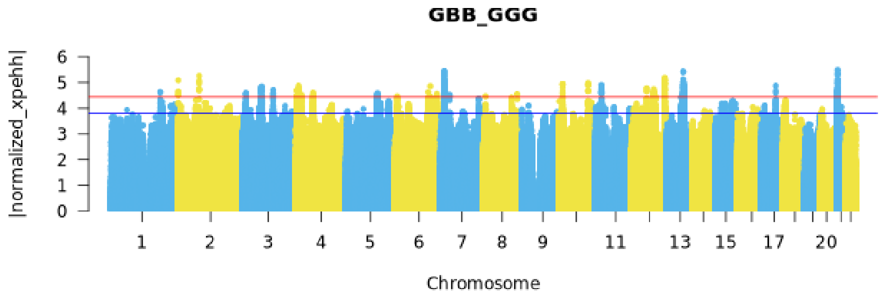


Fig. S13: Genome-wide Manhattan plot of unphased XP-EHH values for GBB vs. GGG. The blue horizontal line marks the top 0.05% cutoff, and the red horizontal line marks the top 0.005% cutoff.

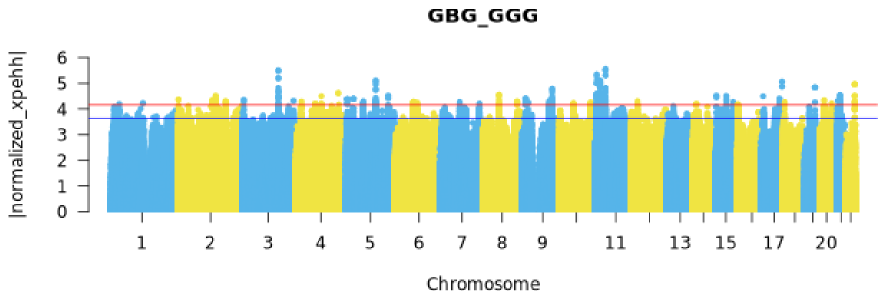


Fig. S14: Genome-wide Manhattan plot of unphased XP-EHH values for GBG vs. GGG. The blue horizontal line marks the top 0.05% cutoff, and the red horizontal line marks the top 0.005% cutoff.

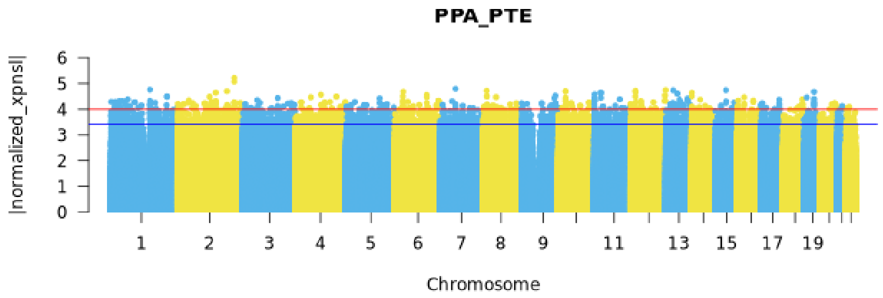


Fig. S15: Genome-wide Manhattan plot of unphased XP-nSL values for PPA vs. PTE. The blue horizontal line marks the top 0.05% cutoff, and the red horizontal line marks the top 0.005% cutoff.

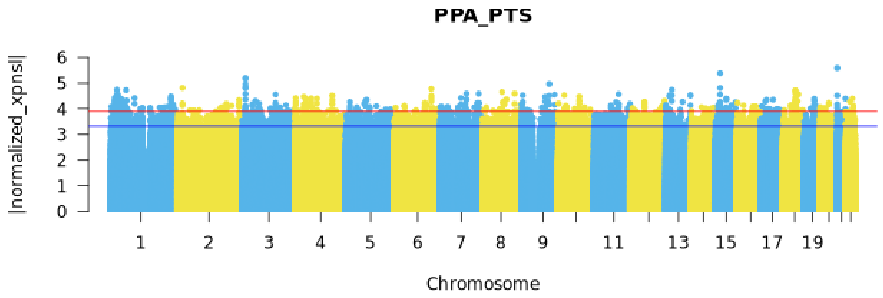


Fig. S16: Genome-wide Manhattan plot of unphased XP-nSL values for PPA vs. PTS. The blue horizontal line indicates the top 0.05% cutoff, and the red horizontal line indicates the top 0.005% cutoff.

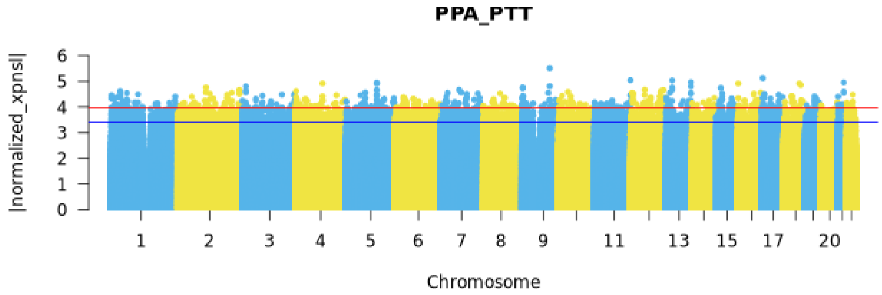


Fig. S17: Genome-wide Manhattan plot of unphased XP-nSL values for PPA vs. PTT. The blue horizontal line marks the top 0.05% cutoff, and the red horizontal line marks the top 0.005% cutoff.

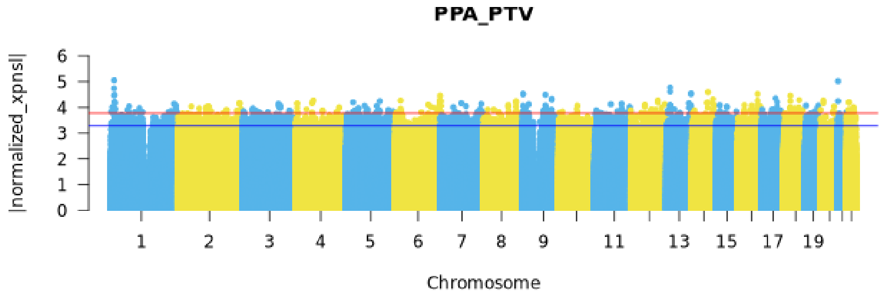


Fig. S18: Genome-wide Manhattan plot of unphased XP-nSL values for PPA vs. PTV. The blue horizontal line marks the top 0.05% cutoff, and the red horizontal line marks the top 0.005% cutoff.

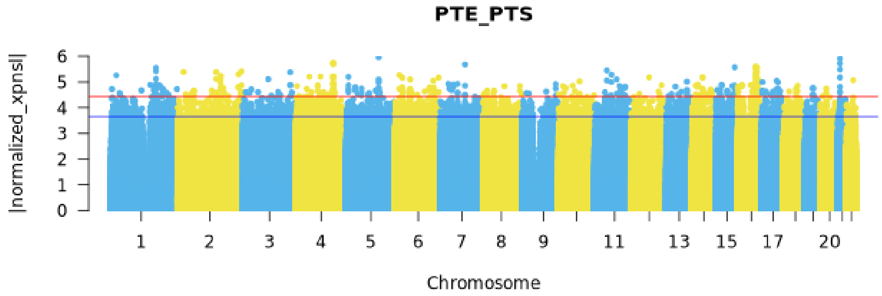


Fig. S19: Genome-wide Manhattan plot of unphased XP-nSL values for PTE vs. PTS. The blue horizontal line marks the top 0.05% cutoff, and the red horizontal line marks the top 0.005% cutoff.

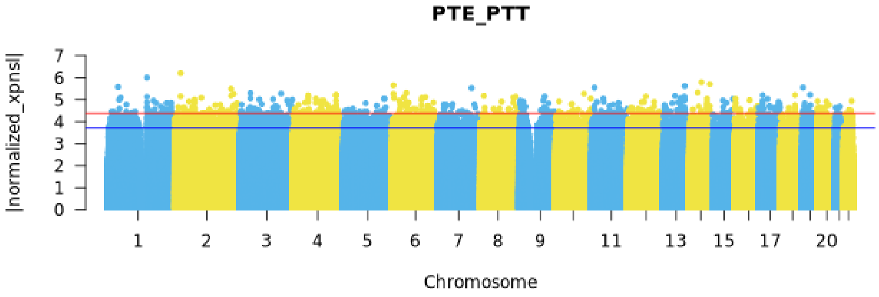


Fig. S20: Genome-wide Manhattan plot of unphased XP-nSL values for PTE vs. PTT. The blue horizontal line marks the top 0.05% cutoff, and the red horizontal line marks the top 0.005% cutoff.

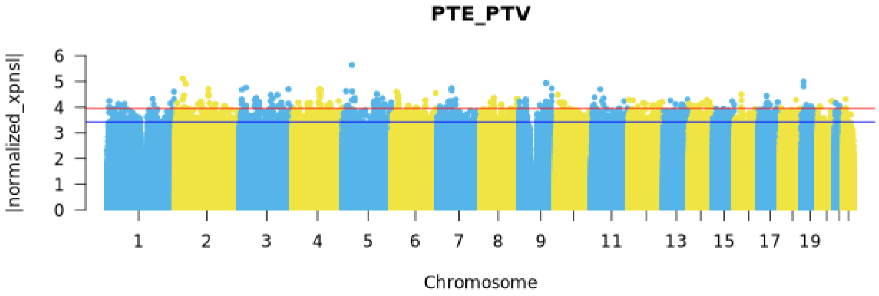


Fig. S21: Genome-wide Manhattan plot of unphased XP-nSL values for PTE vs. PTV. The blue horizontal line marks the top 0.05% cutoff, and the red horizontal line marks the top 0.005% cutoff.

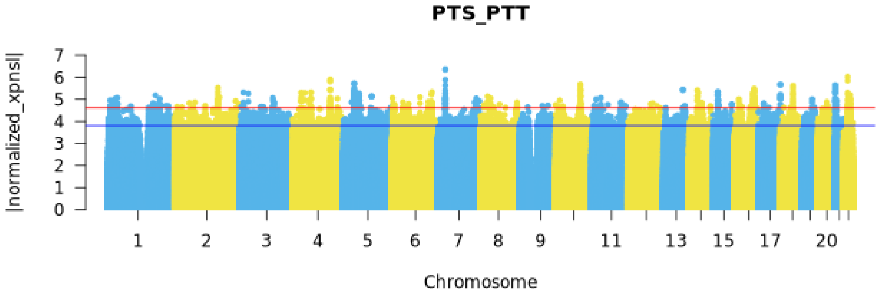


Fig. S22: Genome-wide Manhattan plot of unphased XP-nSL values for PTS vs. PTT. The blue horizontal line marks the top 0.05% cutoff, and the red horizontal line marks the top 0.005% cutoff.

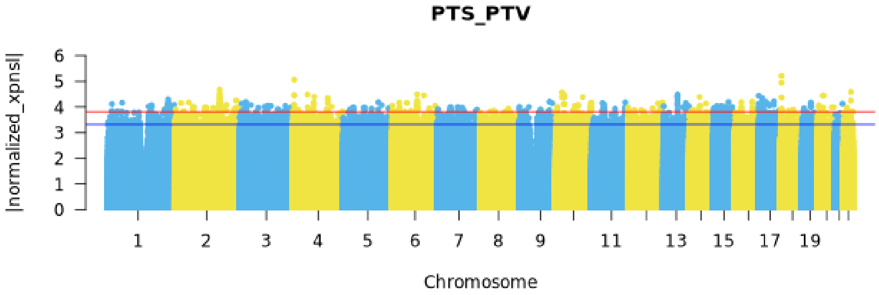


Fig. S23: Genome-wide Manhattan plot of unphased XP-nSL values for PTS vs. PTV. The blue horizontal line marks the top 0.05% cutoff, and the red horizontal line marks the top 0.005% cutoff.

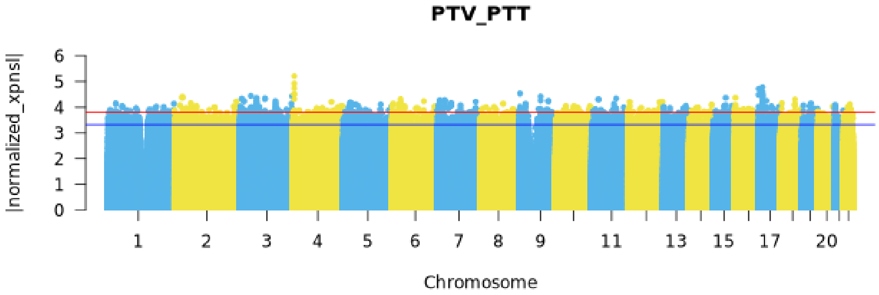


Fig. S24: Genome-wide Manhattan plot of unphased XP-nSL values for PTV vs. PTT. The blue horizontal line marks the top 0.05% cutoff, and the red horizontal line marks the top 0.005% cutoff.

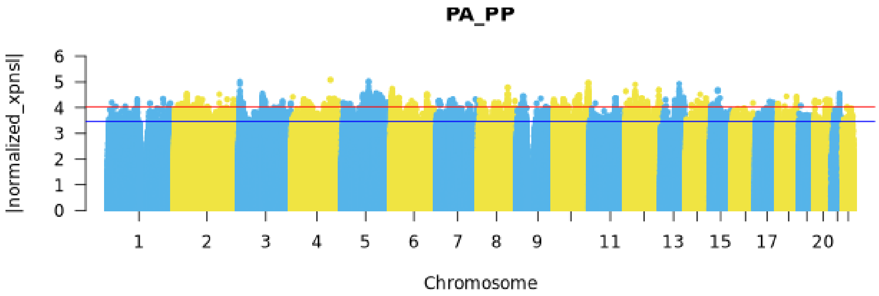


Fig. S25: Genome-wide Manhattan plot of unphased XP-nSL values for PA vs. PP. The blue horizontal line marks the top 0.05% cutoff, and the red horizontal line marks the top 0.005% cutoff.

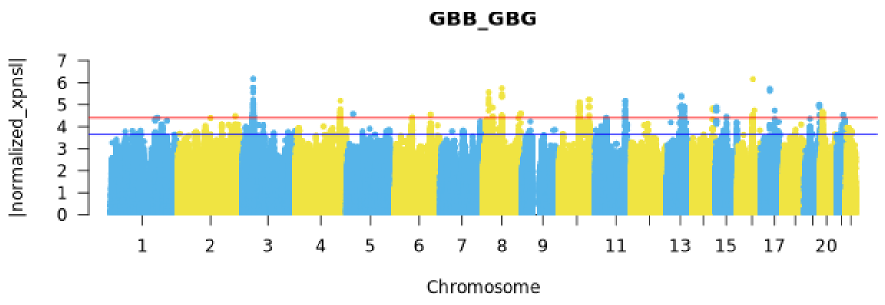


Fig. S26: Genome-wide Manhattan plot of unphased XP-nSL values for GBB vs. GBG. The blue horizontal line marks the top 0.05% cutoff, and the red horizontal line marks the top 0.005% cutoff.

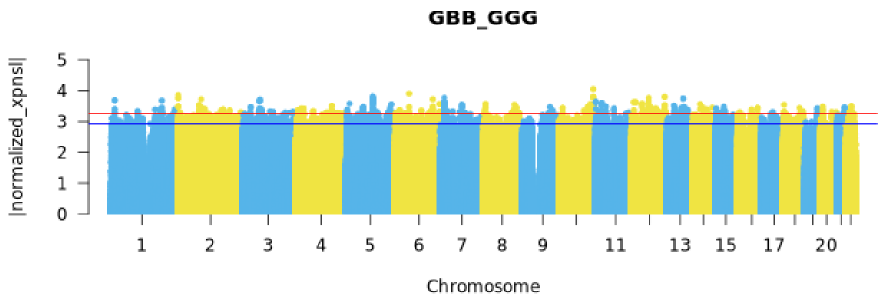


Fig. S27: Genome-wide Manhattan plot of unphased XP-nSL values for GBB vs. GGG. The blue horizontal line marks the top 0.05% cutoff, and the red horizontal line marks the top 0.005% cutoff.

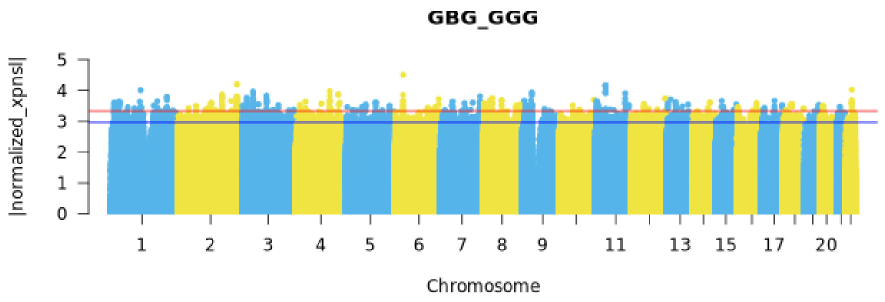


Fig. S28: Genome-wide Manhattan plot of unphased XP-EHH values for GBG vs. GGG. The blue horizontal line marks the top 0.05% cutoff, and the red horizontal line marks the top 0.005% cutoff.

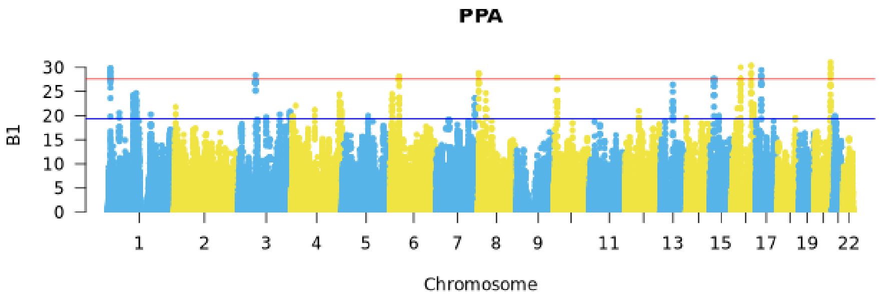


Fig. S29: Genome-wide Manhattan plot of $\beta^{(1)}$ values for PPA. The blue horizontal line marks the top 0.05% cutoff, and the red horizontal line marks the top 0.005% cutoff.

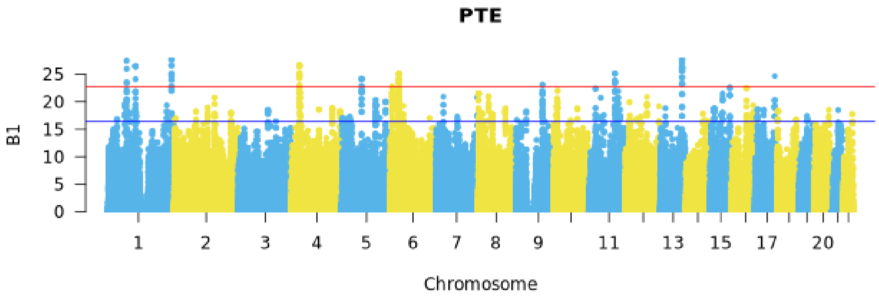


Fig. S30: Genome-wide Manhattan plot of $\beta^{(1)}$ values for PTE. The blue horizontal line indicates the top 0.05% cutoff, and the red horizontal line indicates the top 0.005% cutoff.

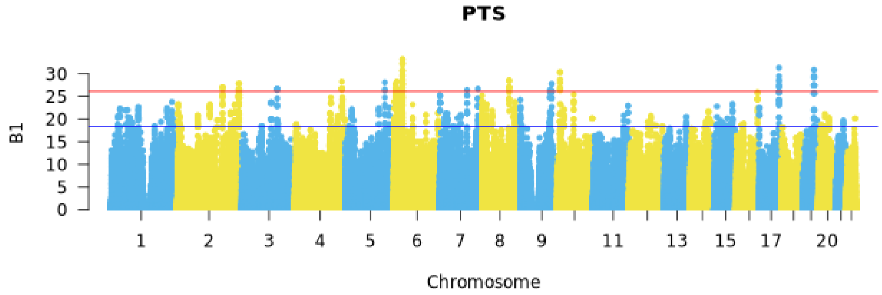


Fig. S31: Genome-wide Manhattan plot of $\beta^{(1)}$ values for PTS. The blue horizontal line marks the top 0.05% cutoff, and the red horizontal line marks the top 0.005% cutoff.

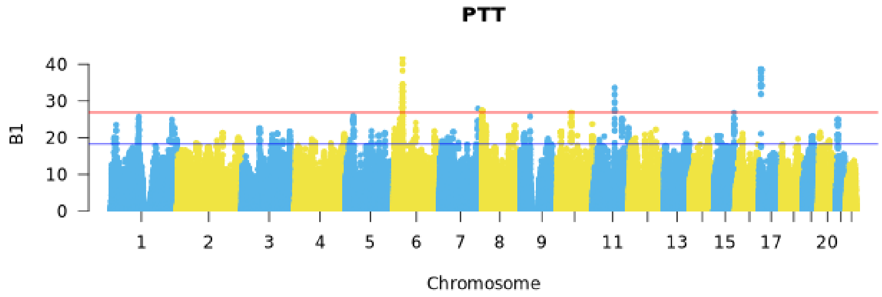


Fig. S32: Genome-wide Manhattan plot of $\beta^{(1)}$ values for PTT. The blue horizontal line marks the top 0.05% cutoff, and the red horizontal line marks the top 0.005% cutoff.

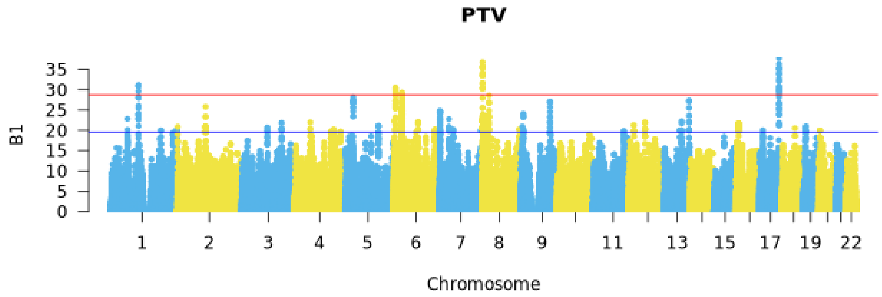


Fig. S33: Genome-wide Manhattan plot of $\beta^{(1)}$ values for PTV. The blue horizontal line marks the top 0.05% cutoff, and the red horizontal line marks the top 0.005% cutoff.

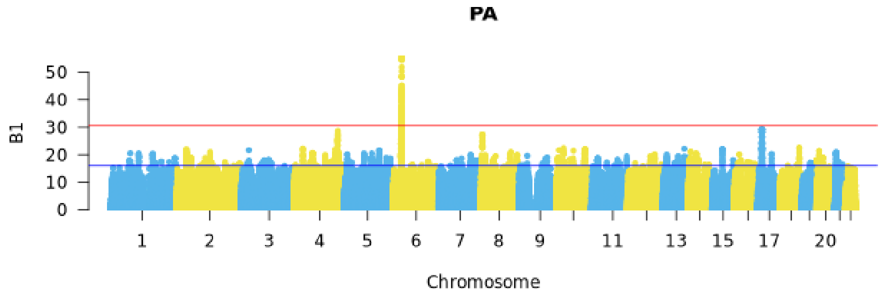


Fig. S34: Genome-wide Manhattan plot of $\beta^{(1)}$ values for PA. The blue horizontal line marks the top 0.05% cutoff, and the red horizontal line marks the top 0.005% cutoff.

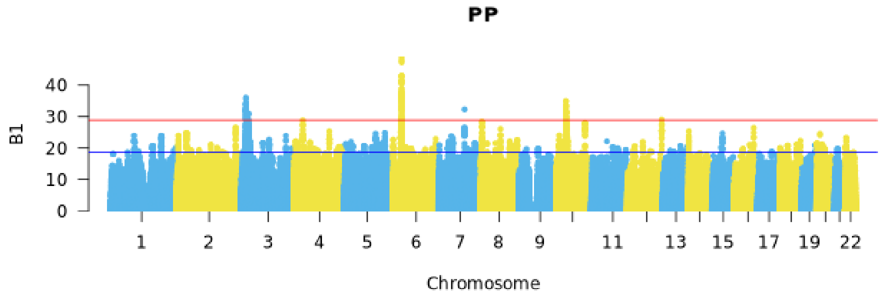


Fig. S35: Genome-wide Manhattan plot of $\beta^{(1)}$ values for PP. The blue horizontal line marks the top 0.05% cutoff, and the red horizontal line marks the top 0.005% cutoff.

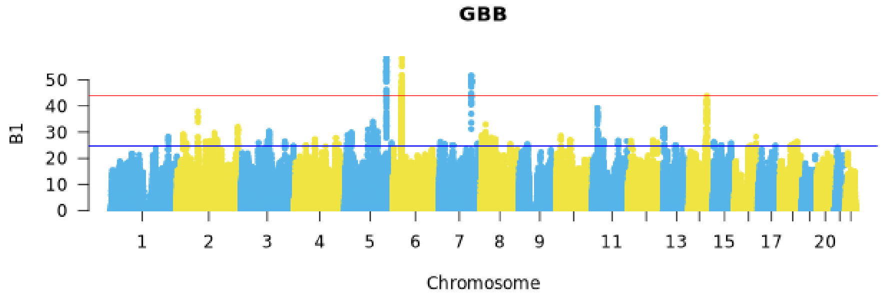


Fig. S36: Genome-wide Manhattan plot of $\beta^{(1)}$ values for GBB. The blue horizontal line marks the top 0.05% cutoff, and the red horizontal line marks the top 0.005% cutoff.

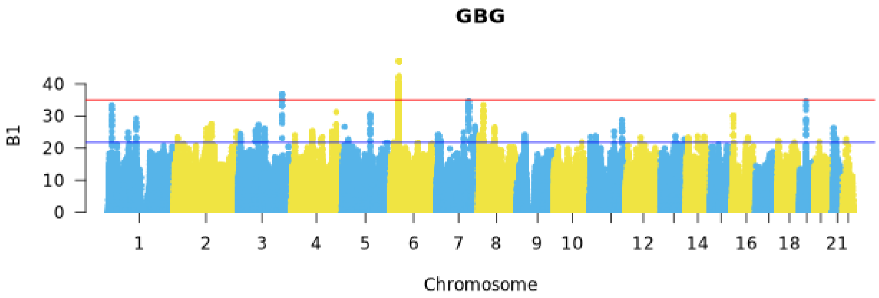


Fig. S37: Genome-wide Manhattan plot of $\beta^{(1)}$ values for GBG. The blue horizontal line marks the top 0.05% cutoff, and the red horizontal line marks the top 0.005% cutoff.

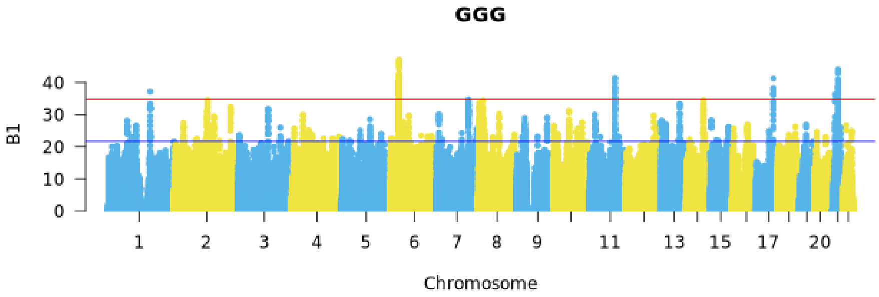


Fig. S38: Genome-wide Manhattan plot of $\beta^{(1)}$ values for GGG. The blue horizontal line marks the top 0.05% cutoff, and the red horizontal line marks the top 0.005% cutoff.

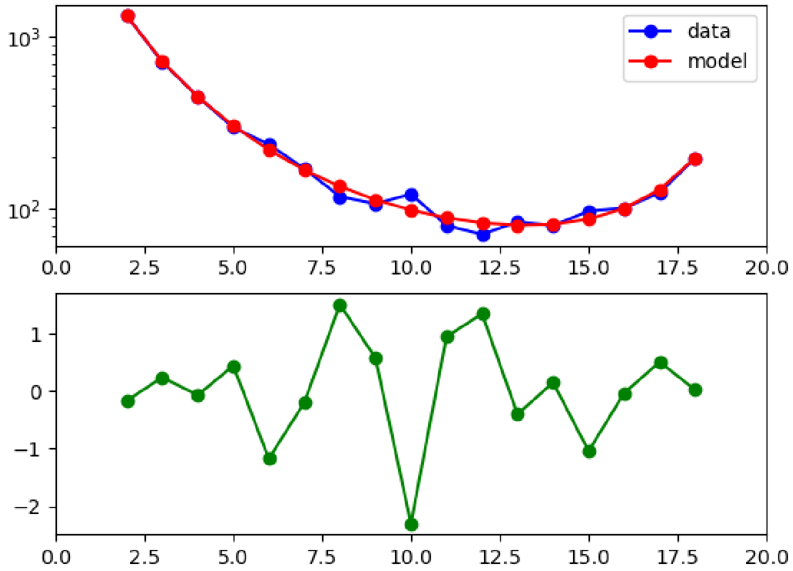


Fig. S39: Model fit to the AFS using synonymous SNPs from PPA. The blue line shows the observed AFS from the data, the red line shows the inferred AFS based on the maximum likelihood estimates in Table S88, and the green line shows the residual between the observed and model AFS.

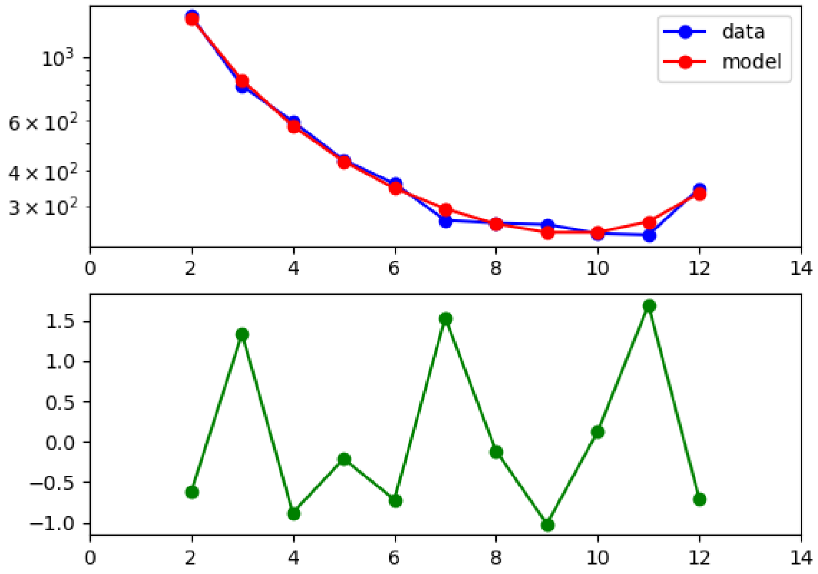


Fig. S40: Model fit to the AFS using synonymous SNPs from PTE. The blue line shows the observed AFS from the data, the red line shows the inferred AFS based on the maximum likelihood estimates in Table S88, and the green line shows the residual between the observed and model AFS.

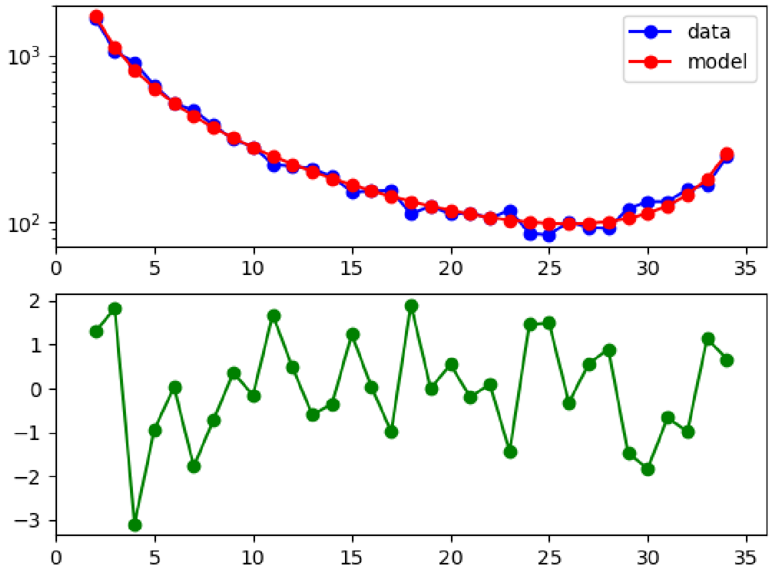


Fig. S41: Model fit to the AFS using synonymous SNPs from PTE. The blue line shows the observed AFS from the data, the red line shows the inferred AFS based on the maximum likelihood estimates in Table S88, and the green line shows the residual between the observed and model AFS.

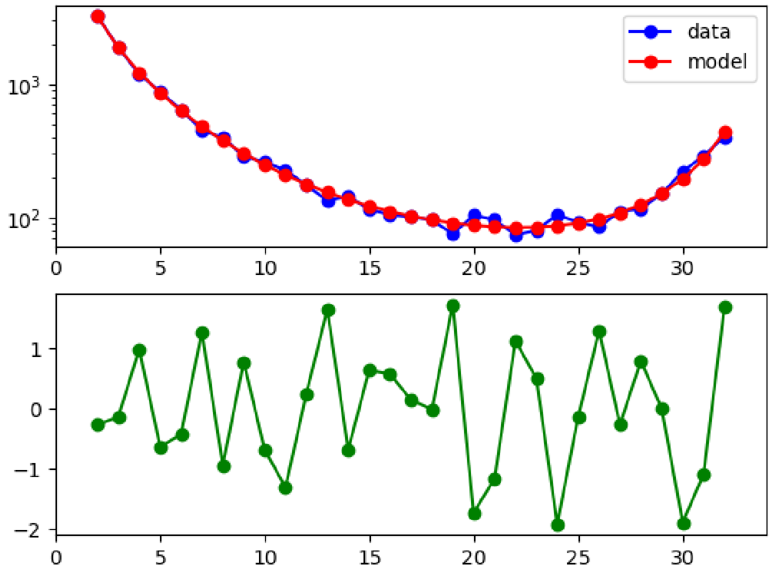


Fig. S42: Model fit to the AFS using synonymous SNPs from PTT. The blue line shows the observed AFS from the data, the red line shows the inferred AFS based on the maximum likelihood estimates in Table S88, and the green line shows the residual between the observed and model AFS.

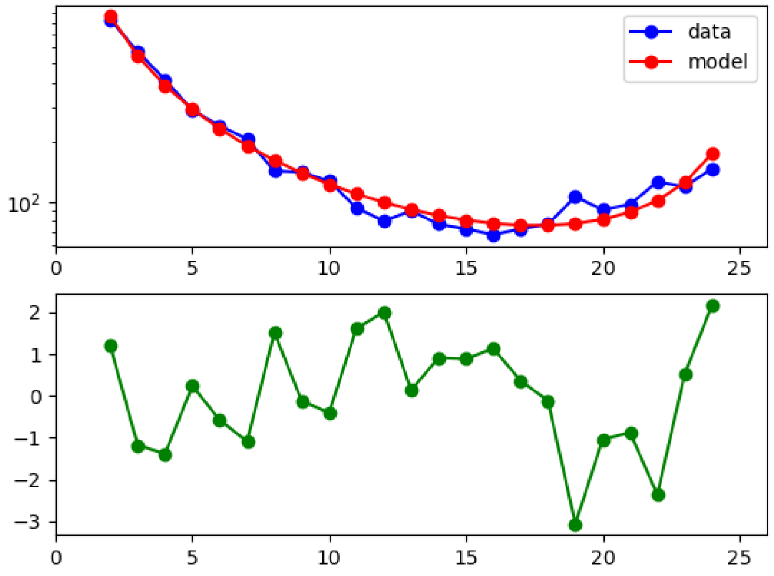


Fig. S43: Model fit to the AFS using synonymous SNPs from PTV. The blue line shows the observed AFS from the data, the red line shows the inferred AFS based on the maximum likelihood estimates in Table S88, and the green line shows the residual between the observed and model AFS.

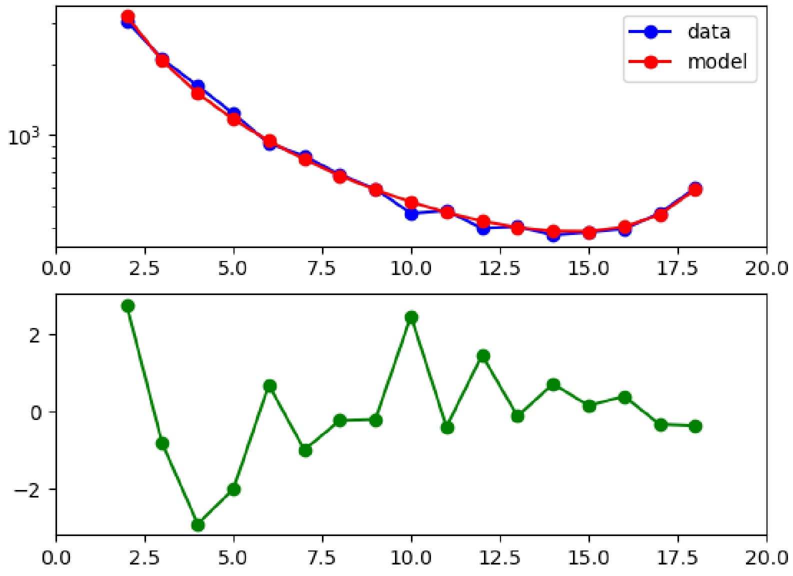


Fig. S44: Model fit to the AFS using synonymous SNPs from PA. The blue line shows the observed AFS from the data, the red line shows the inferred AFS based on the maximum likelihood estimates in Table S88, and the green line shows the residual between the observed and model AFS.

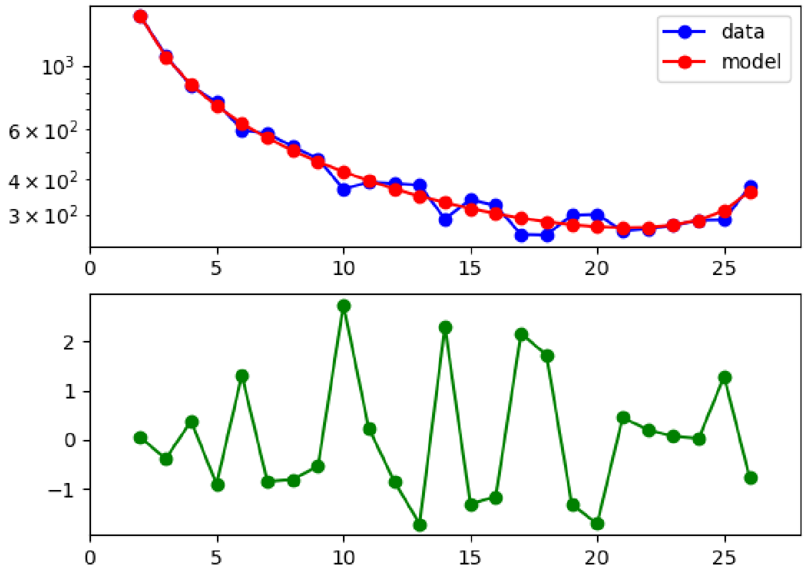


Fig. S45: Model fit to the AFS using synonymous SNPs from PP. The blue line shows the observed AFS from the data, the red line shows the inferred AFS based on the maximum likelihood estimates in Table S88, and the green line shows the residual between the observed and model AFS.

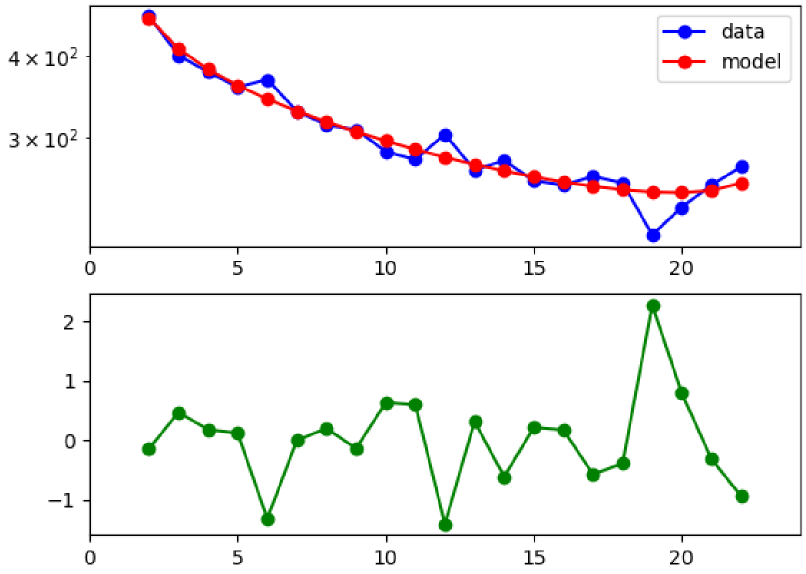


Fig. S46: Model fit to the AFS using synonymous SNPs from GBB. The blue line shows the observed AFS from the data, the red line shows the inferred AFS based on the maximum likelihood estimates in Table S88, and the green line shows the residual between the observed and model AFS.

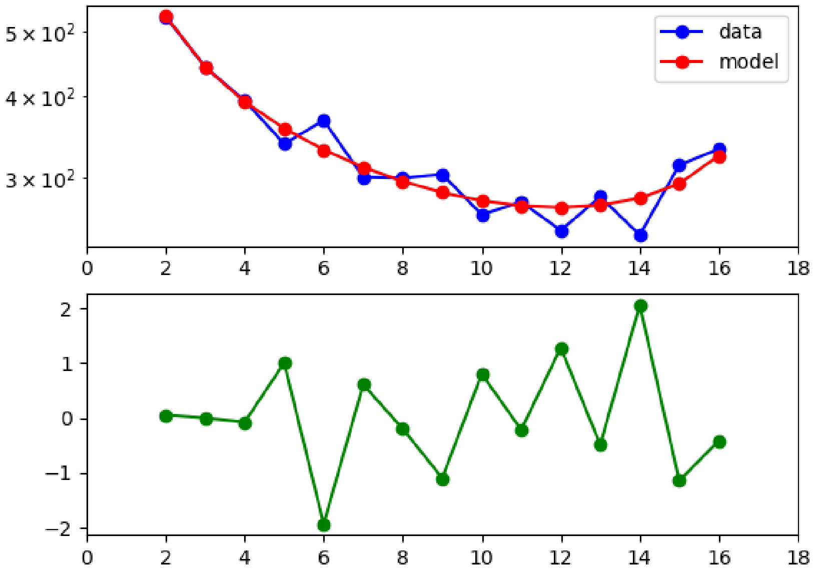


Fig. S47: Model fit to the AFS using synonymous SNPs from GBG. The blue line shows the observed AFS from the data, the red line shows the inferred AFS based on the maximum likelihood estimates in Table S88, and the green line shows the residual between the observed and model AFS.

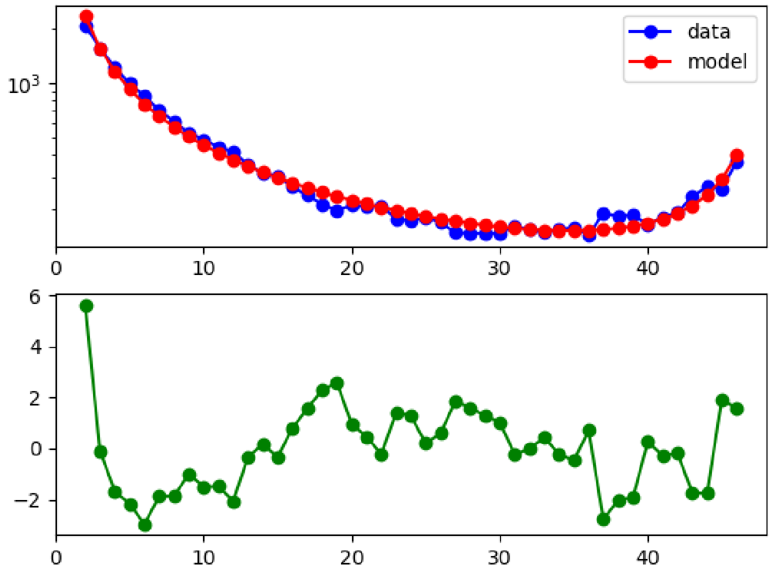


Fig. S48: Model fit to the AFS using synonymous SNPs from GGG. The blue line shows the observed AFS from the data, the red line shows the inferred AFS based on the maximum likelihood estimates in Table S88, and the green line shows the residual between the observed and model AFS.

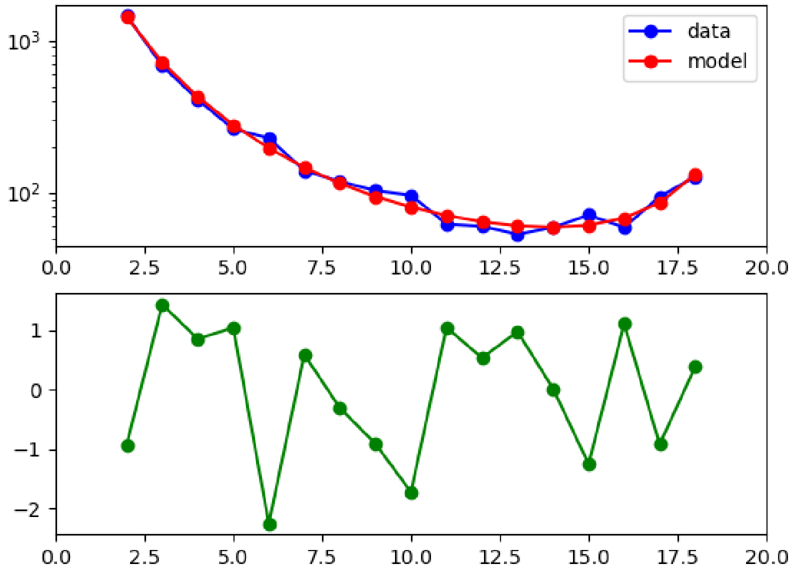


Fig. S49: Model fit to the AFS using non-synonymous SNPs from PPA. The blue line shows the observed AFS from the data, the red line shows the inferred AFS based on the maximum likelihood estimates in Fig. 3 and Table S88, and the green line shows the residual between the observed and model AFS.

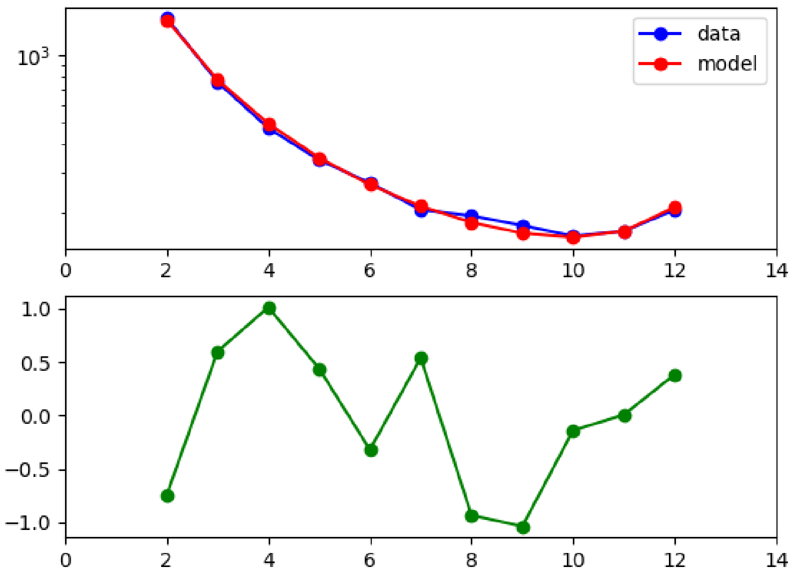


Fig. S50: Model fit to the AFS using non-synonymous SNPs from PTE. The blue line shows the observed AFS from the data, the red line shows the inferred AFS based on the maximum likelihood estimates in Fig. 3 and Table S88, and the green line shows the residual between the observed and model AFS.

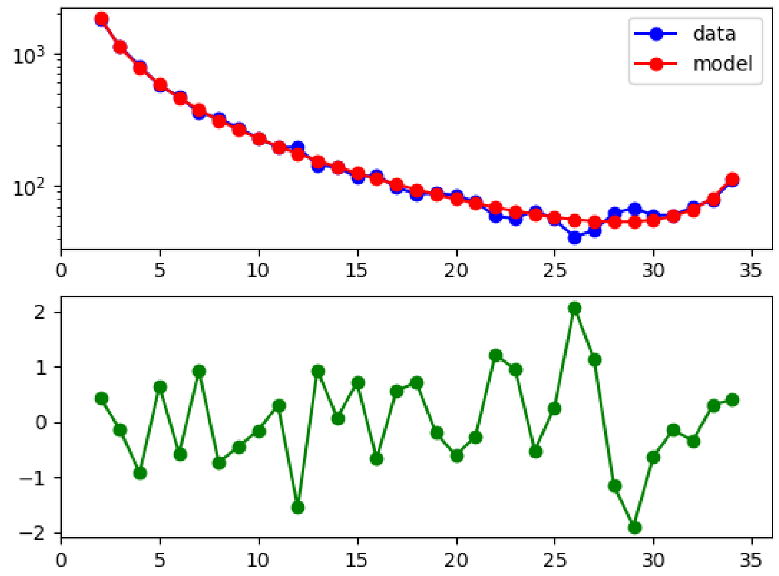


Fig. S51: Model fit to the AFS using non-synonymous SNPs from PTS. The blue line shows the observed AFS from the data, the red line shows the inferred AFS based on the maximum likelihood estimates in Fig. 3 and Table S88, and the green line shows the residual between the observed and model AFS.

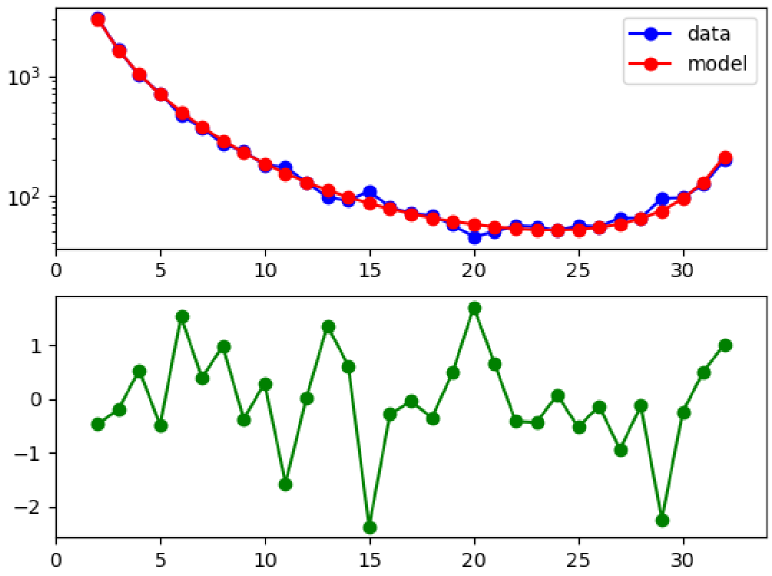


Fig. S52: Model fit to the AFS using non-synonymous SNPs from PTT. The blue line shows the observed AFS from the data, the red line shows the inferred AFS based on the maximum likelihood estimates in Fig. 3 and Table S88, and the green line shows the residual between the observed and model AFS.

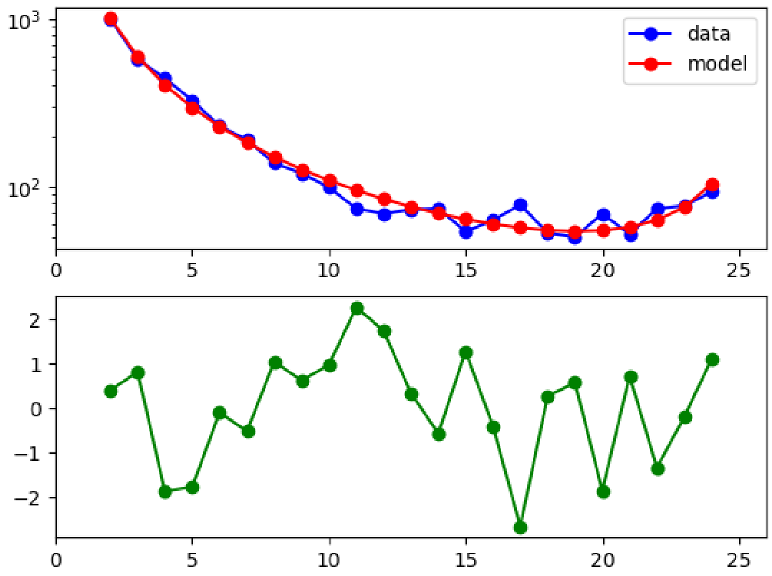


Fig. S53: Model fit to the AFS using non-synonymous SNPs from PTV. The blue line shows the observed AFS from the data, the red line shows the inferred AFS based on the maximum likelihood estimates in Fig. 3 and Table S88, and the green line shows the residual between the observed and model AFS.

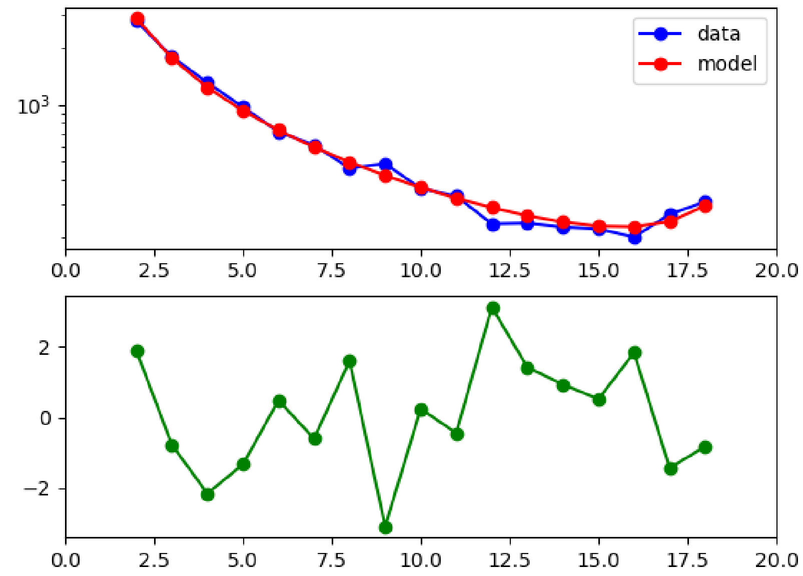


Fig. S54: Model fit to the AFS using non-synonymous SNPs from PA. The blue line shows the observed AFS from the data, the red line shows the inferred AFS based on the maximum likelihood estimates in Fig. 3 and Table S88, and the green line shows the residual between the observed and model AFS.

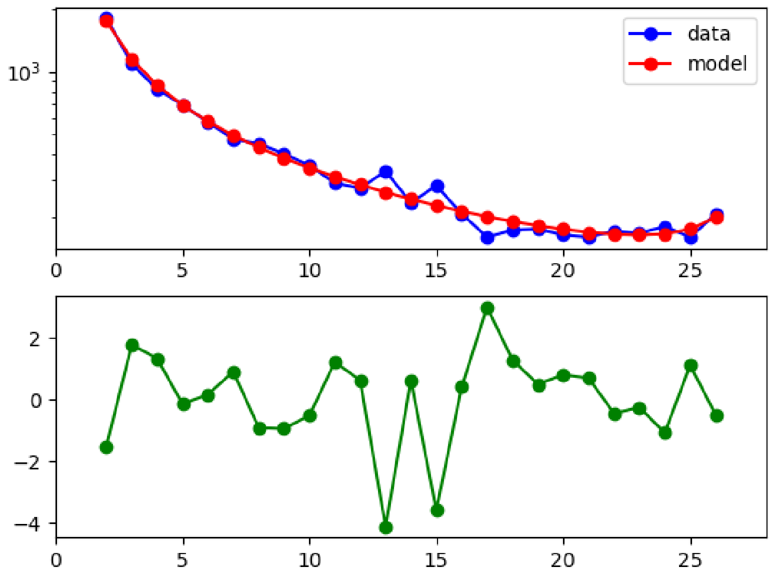


Fig. S55: Model fit to the AFS using non-synonymous SNPs from PP. The blue line shows the observed AFS from the data, the red line shows the inferred AFS based on the maximum likelihood estimates in Fig. 3 and Table S88, and the green line shows the residual between the observed and model AFS.

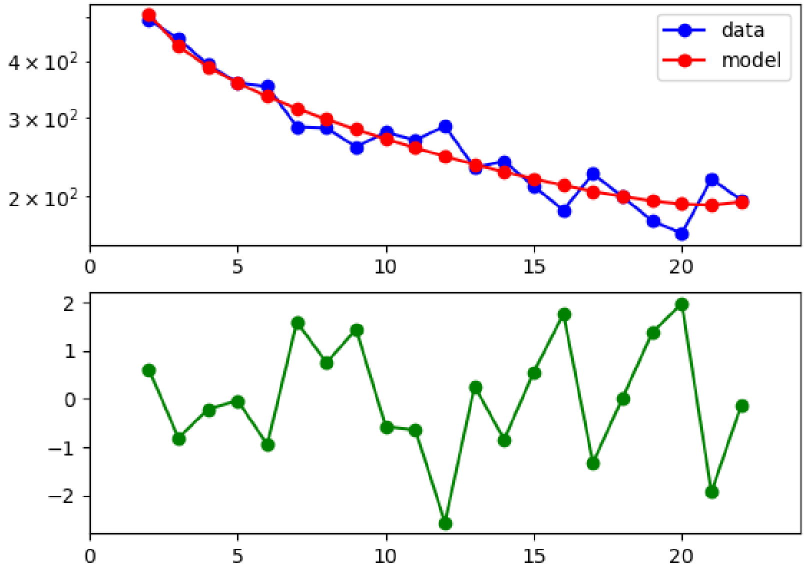


Fig. S56: Model fit to the AFS using non-synonymous SNPs from GBB. The blue line shows the observed AFS from the data, the red line shows the inferred AFS based on the maximum likelihood estimates in Fig. 3 and Table S88, and the green line shows the residual between the observed and model AFS.

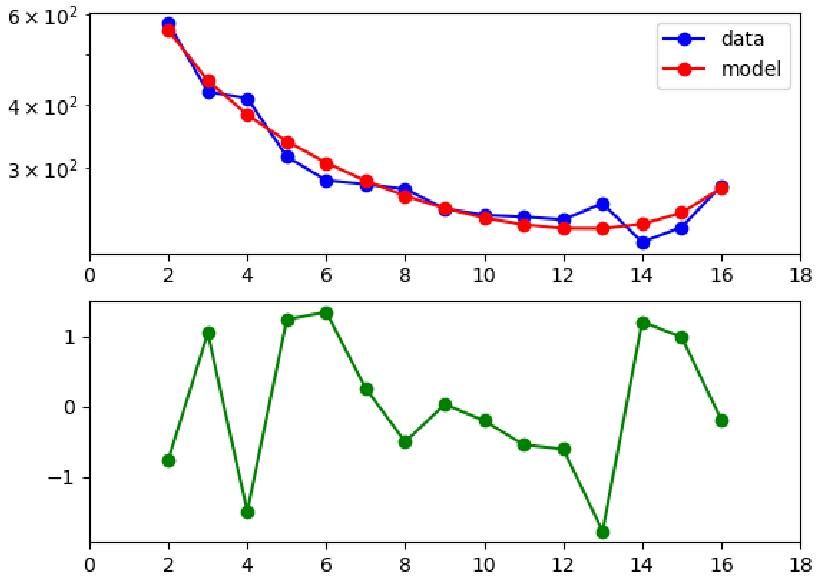


Fig. S57: Model fit to the AFS using non-synonymous SNPs from GBG. The blue line shows the observed AFS from the data, the red line shows the inferred AFS based on the maximum likelihood estimates in Fig. 3 and Table S88, and the green line shows the residual between the observed and model AFS.

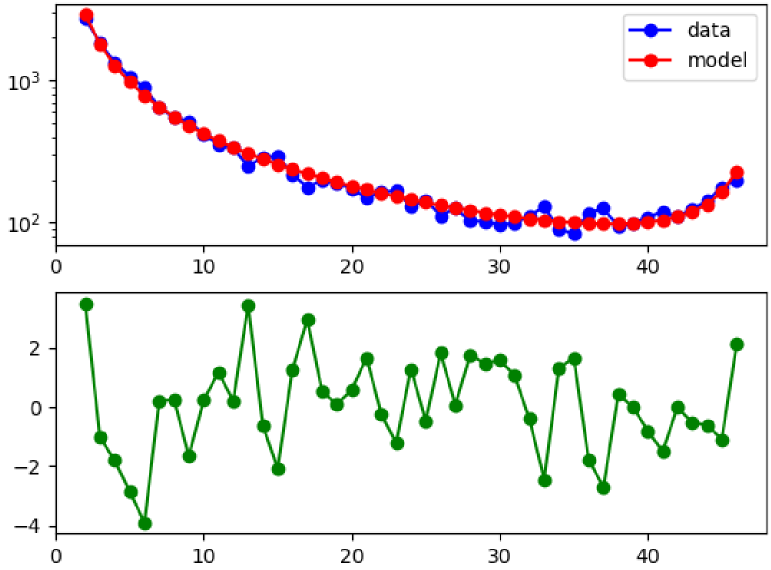


Fig. S58: Model fit to the AFS using non-synonymous SNPs from GGG. The blue line shows the observed AFS from the data, the red line shows the inferred AFS based on the maximum likelihood estimates in Fig. 3 and Table S88, and the green line shows the residual between the observed and model AFS.

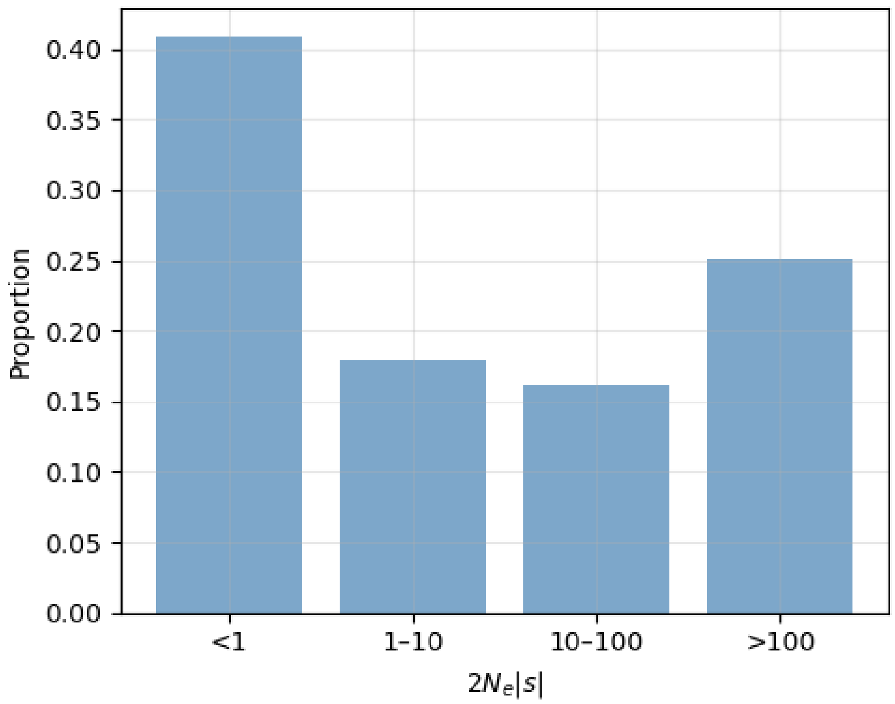


Fig. S59: Proportions of deleterious mutations in PPA, inferred from non-synonymous SNPs using the DFE model based on the maximum likelihood estimates in Fig. 3.

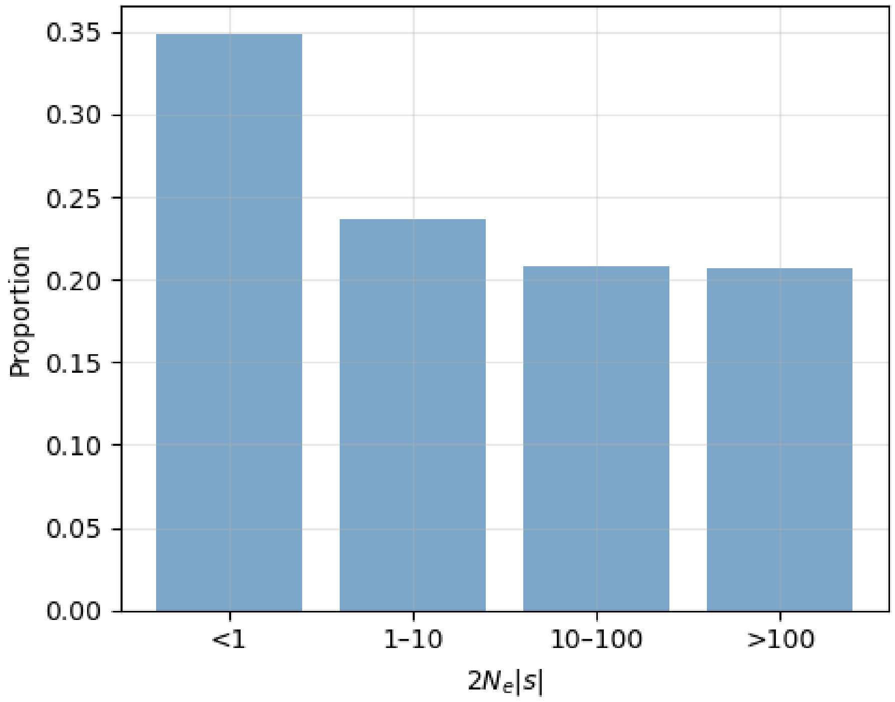


Fig. S60: Proportions of deleterious mutations in PTE, inferred from non-synonymous SNPs using the DFE model based on the maximum likelihood estimates in Fig. 3.

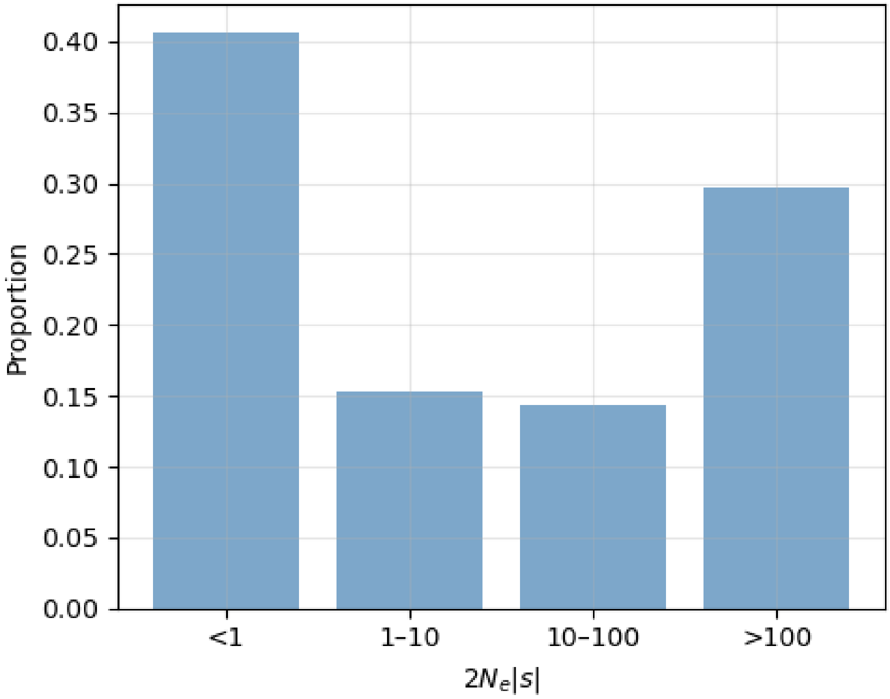


Fig. S61: Proportions of deleterious mutations in PTS, inferred from non-synonymous SNPs using the DFE model based on the maximum likelihood estimates in Fig. 3.

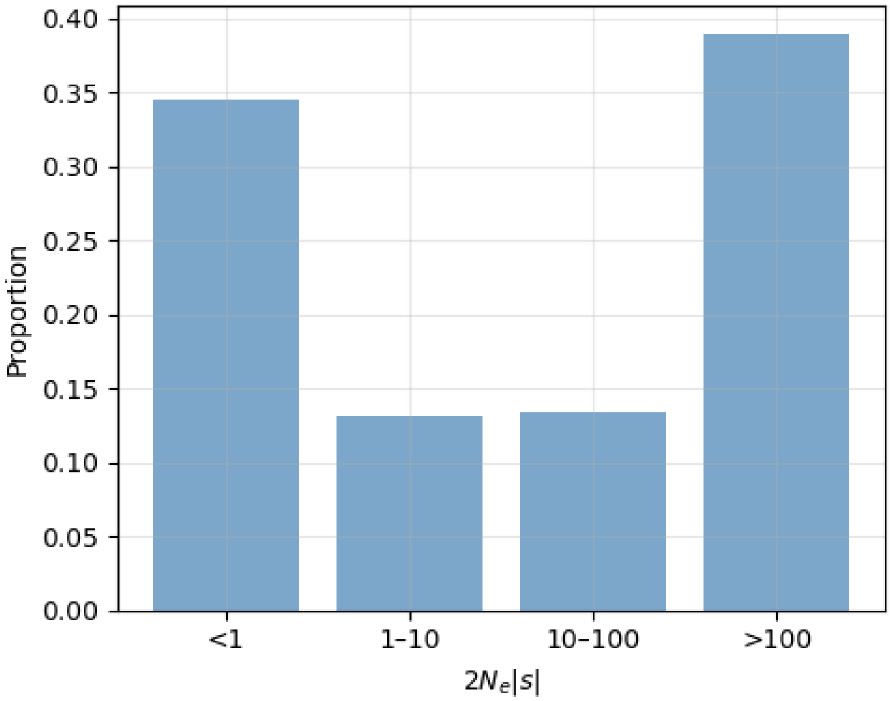


Fig. S62: Proportions of deleterious mutations in PTT, inferred from non-synonymous SNPs using the DFE model based on the maximum likelihood estimates in Fig. 3.

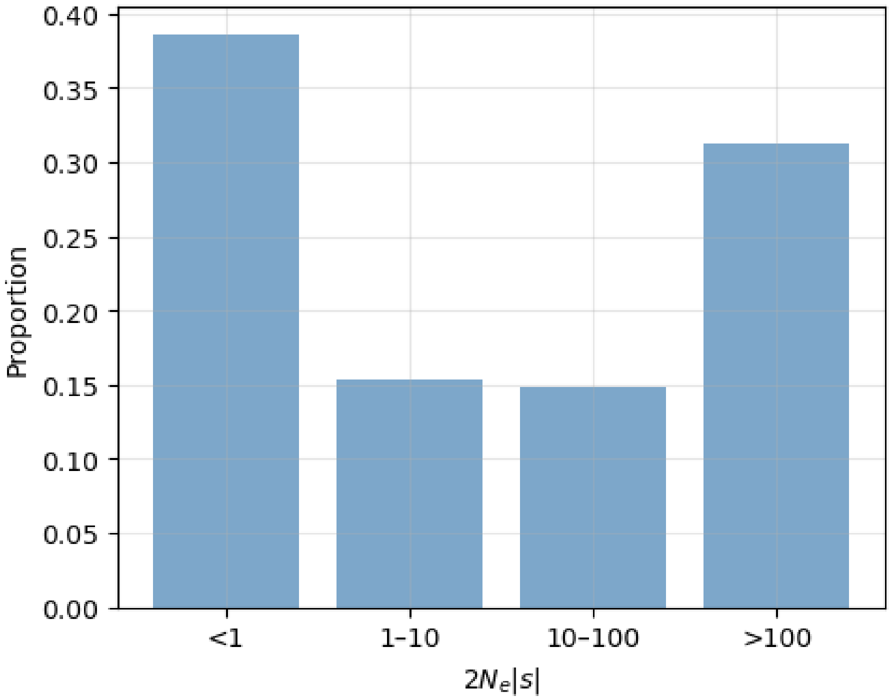


Fig. S63: Proportions of deleterious mutations in PTV, inferred from non-synonymous SNPs using the DFE model based on the maximum likelihood estimates in Fig. 3.

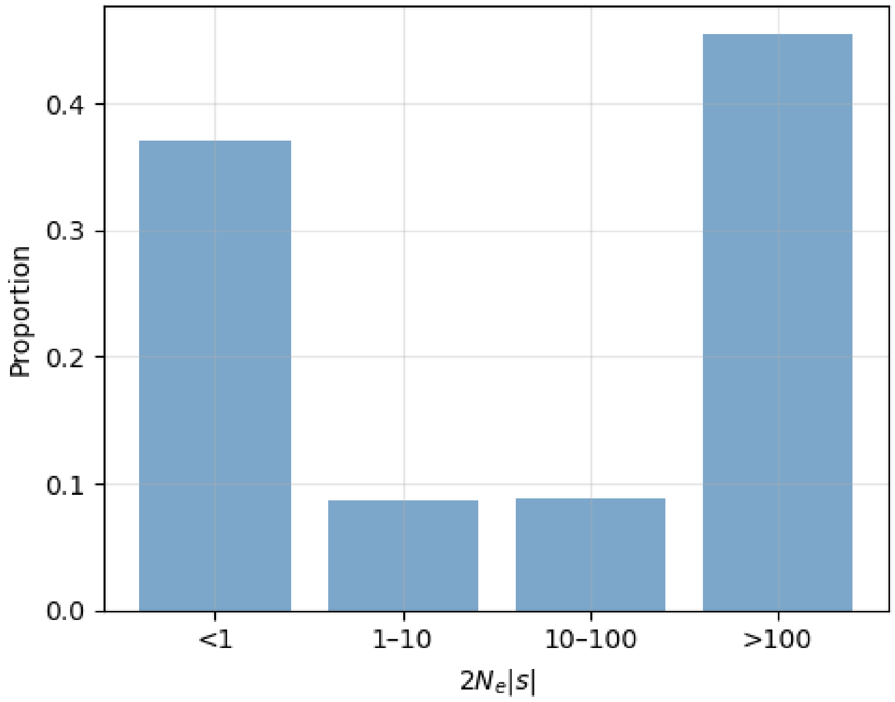


Fig. S64: Proportions of deleterious mutations in PA, inferred from non-synonymous SNPs using the DFE model based on the maximum likelihood estimates in Fig. 3.

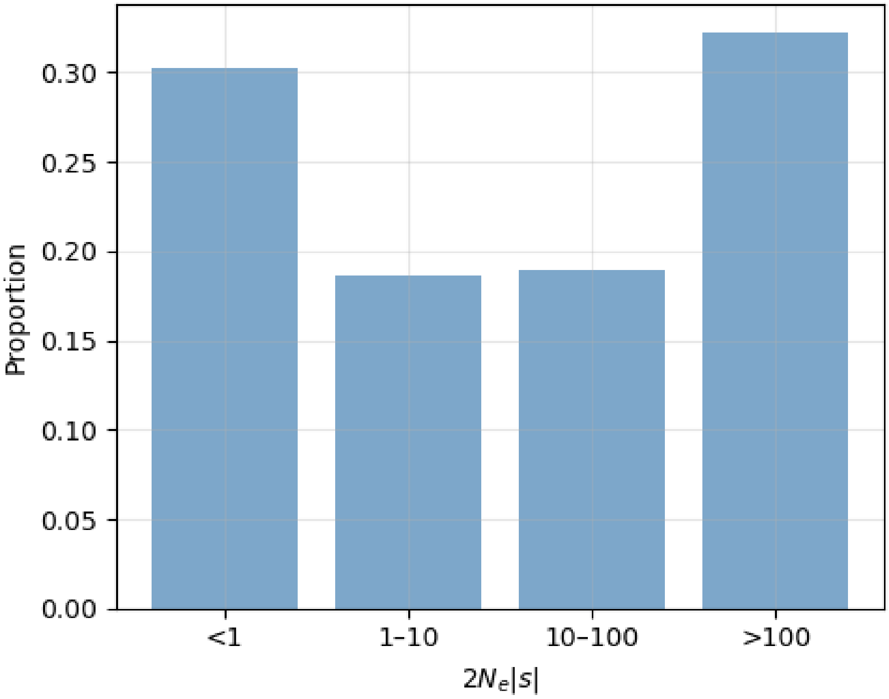


Fig. S65: Proportions of deleterious mutations in PP, inferred from non-synonymous SNPs using the DFE model based on the maximum likelihood estimates in Fig. 3.

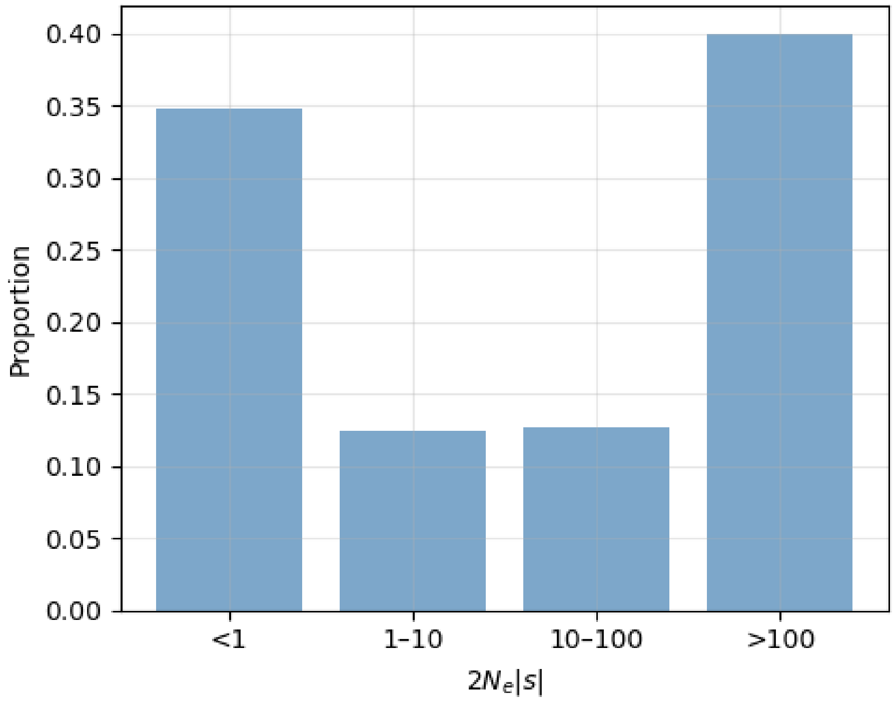


Fig. S66: Proportions of deleterious mutations in GBB, inferred from non-synonymous SNPs using the DFE model based on the maximum likelihood estimates in Fig. 3.

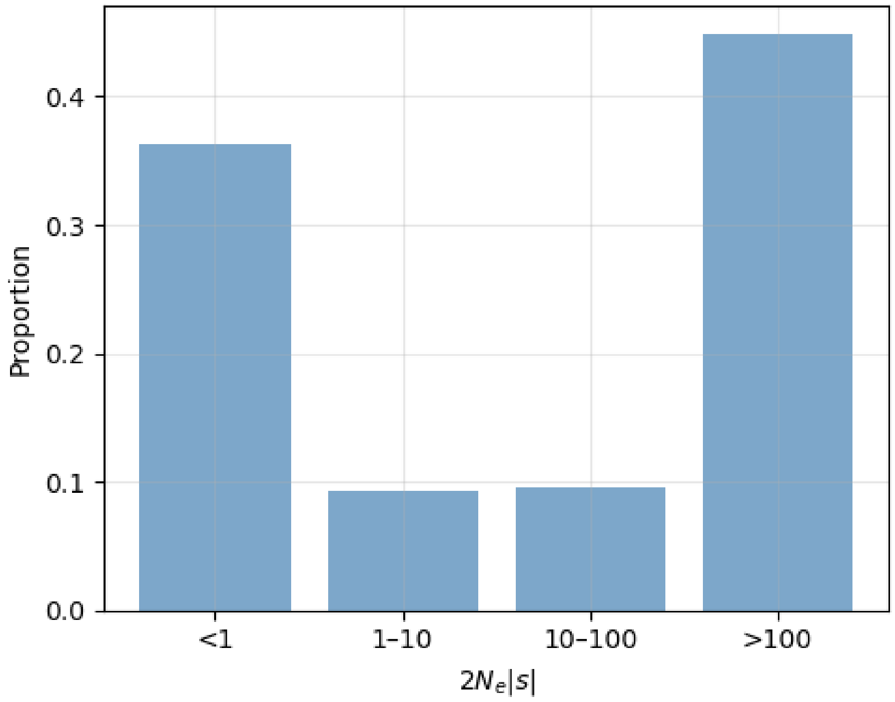


Fig. S67: Proportions of deleterious mutations in GBG, inferred from non-synonymous SNPs using the DFE model based on the maximum likelihood estimates in Fig. 3.

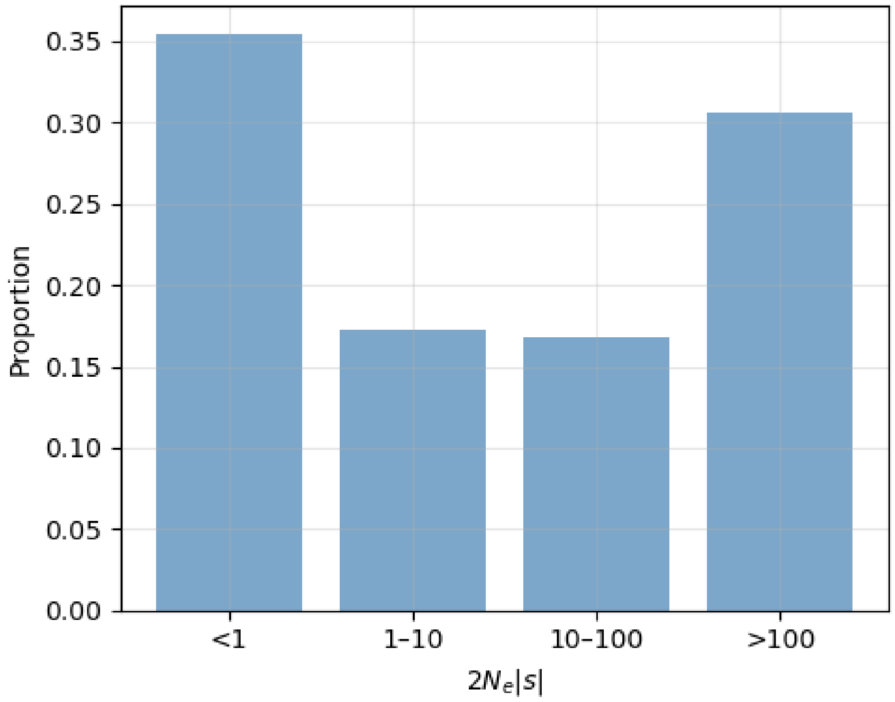


Fig. S68: Proportions of deleterious mutations in GGG, inferred from non-synonymous SNPs using the DFE model based on the maximum likelihood estimates in Fig. 3.

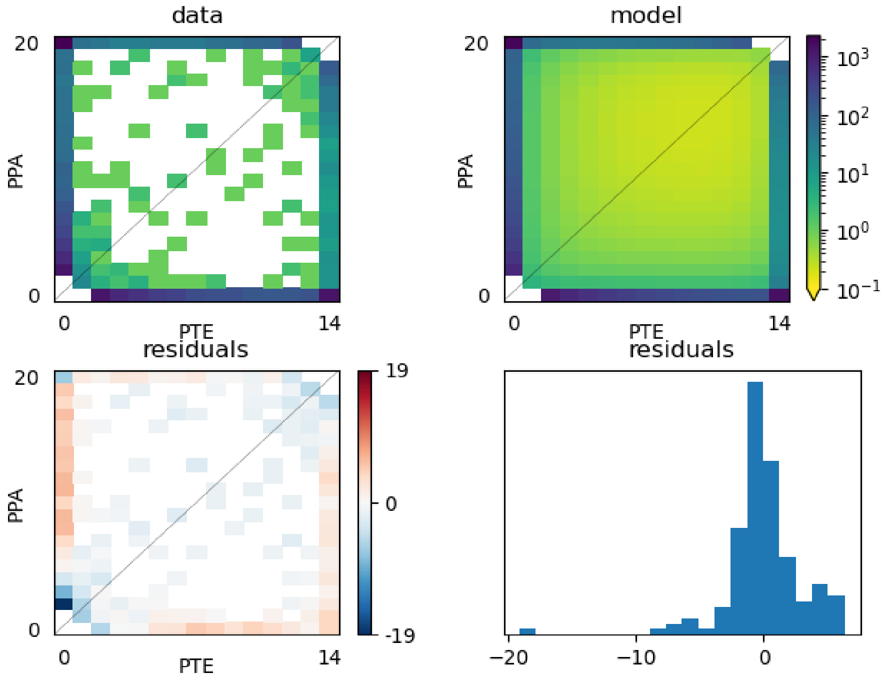


Fig. S69: Model fit to the joint AFS using synonymous SNPs from PPA and PTE. The upper left panel shows the observed joint AFS, the upper right panel shows the inferred AFS based on the maximum likelihood estimates in Table S89, and the remaining panels show the residuals between the observed and model AFS.

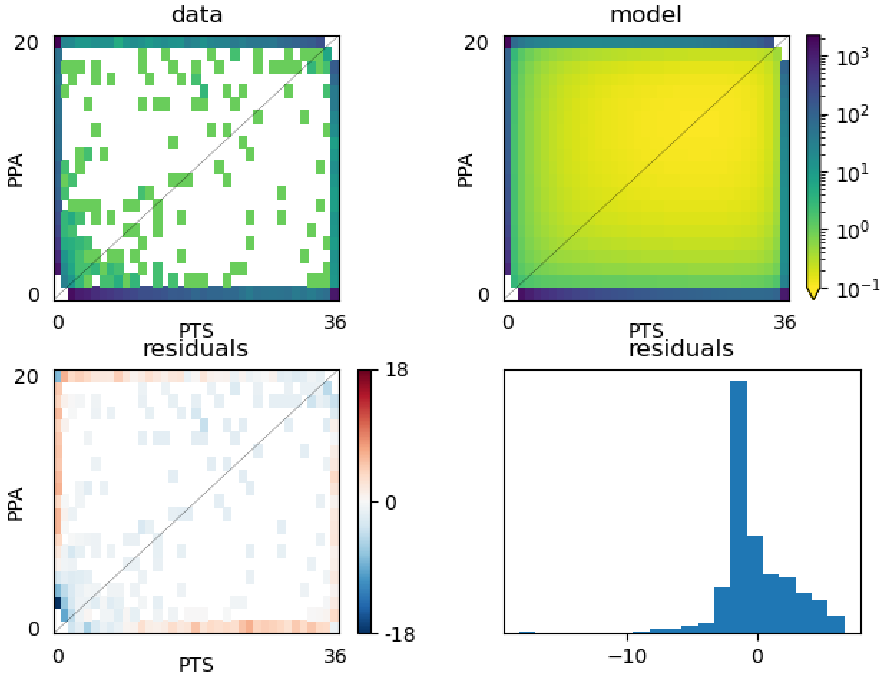


Fig. S70: Model fit to the joint AFS using synonymous SNPs from PPA and PTS. The upper left panel shows the observed joint AFS, the upper right panel shows the inferred AFS based on the maximum likelihood estimates in Table S89, and the remaining panels show the residuals between the observed and model AFS.

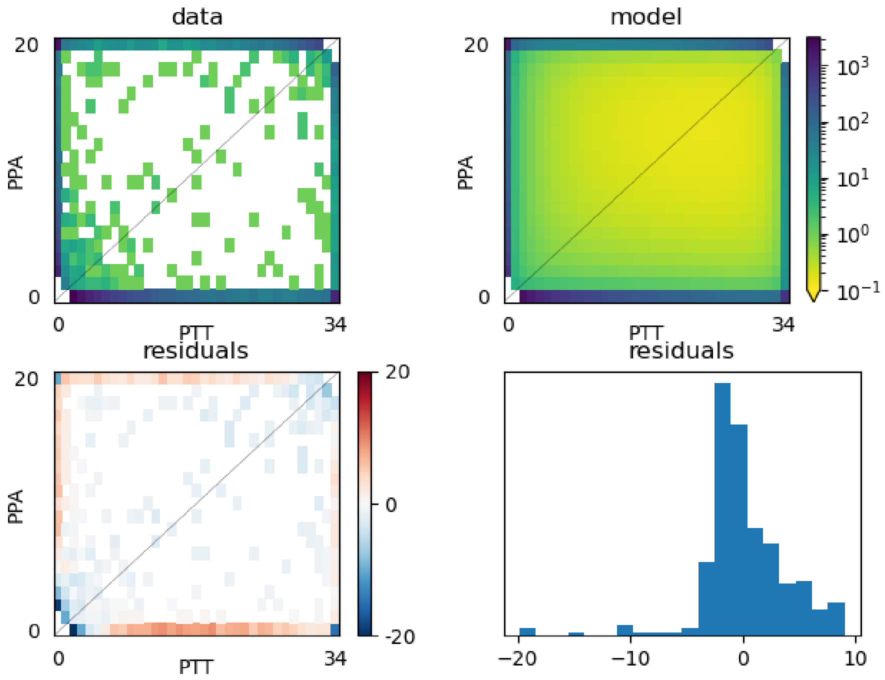


Fig. S71: Model fit to the joint AFS using synonymous SNPs from PPA and PTT. The upper left panel shows the observed joint AFS, the upper right panel shows the inferred AFS based on the maximum likelihood estimates in Table S89, and the remaining panels show the residuals between the observed and model AFS.

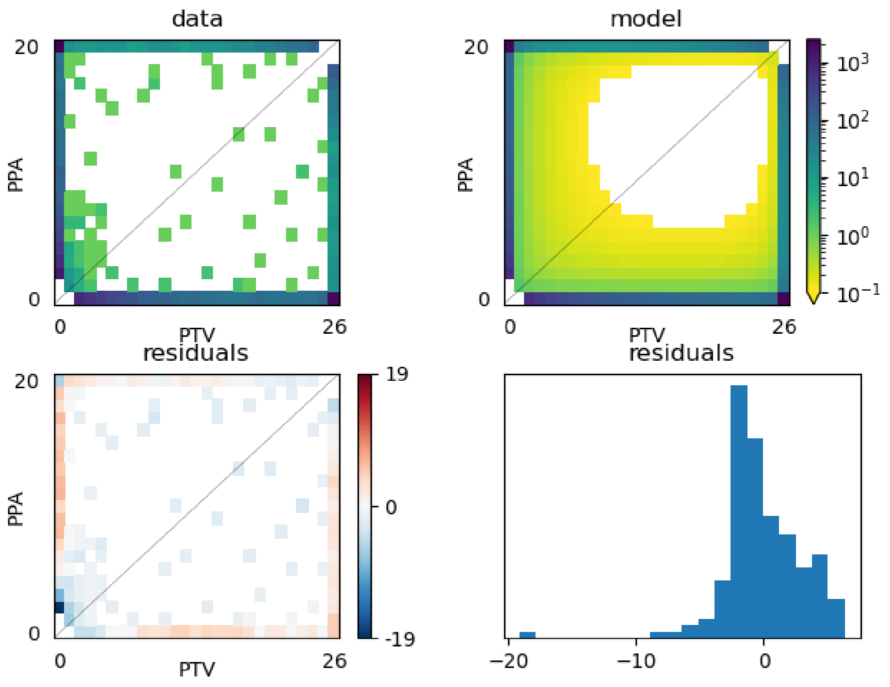


Fig. S72: Model fit to the joint AFS using synonymous SNPs from PPA and PTV. The upper left panel shows the observed joint AFS, the upper right panel shows the inferred AFS based on the maximum likelihood estimates in Table S89, and the remaining panels show the residuals between the observed and model AFS.

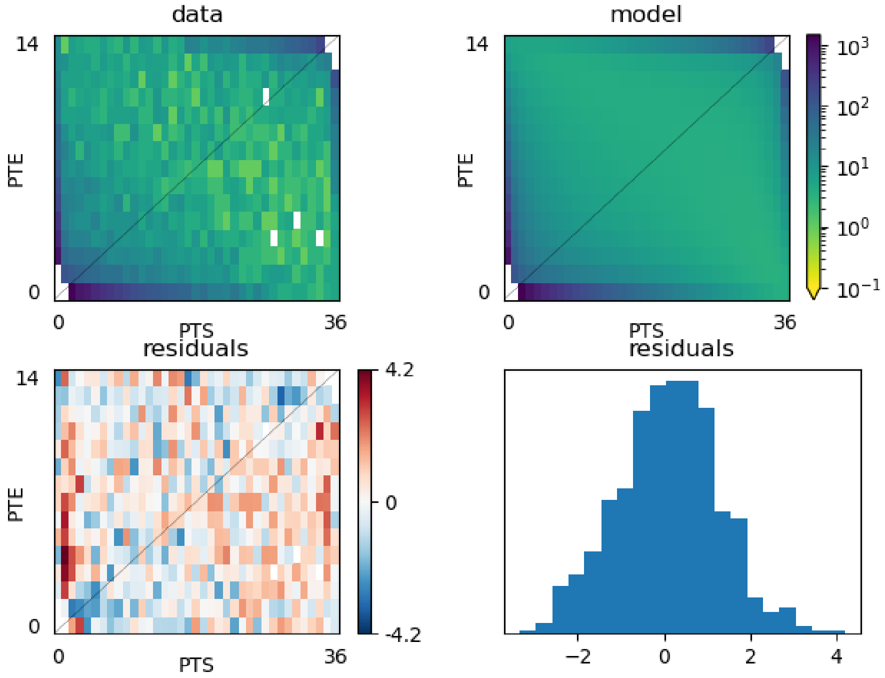


Fig. S73: Model fit to the joint AFS using synonymous SNPs from PTE and PTS. The upper left panel shows the observed joint AFS, the upper right panel shows the inferred AFS based on the maximum likelihood estimates in Table S89, and the remaining panels show the residuals between the observed and model AFS.

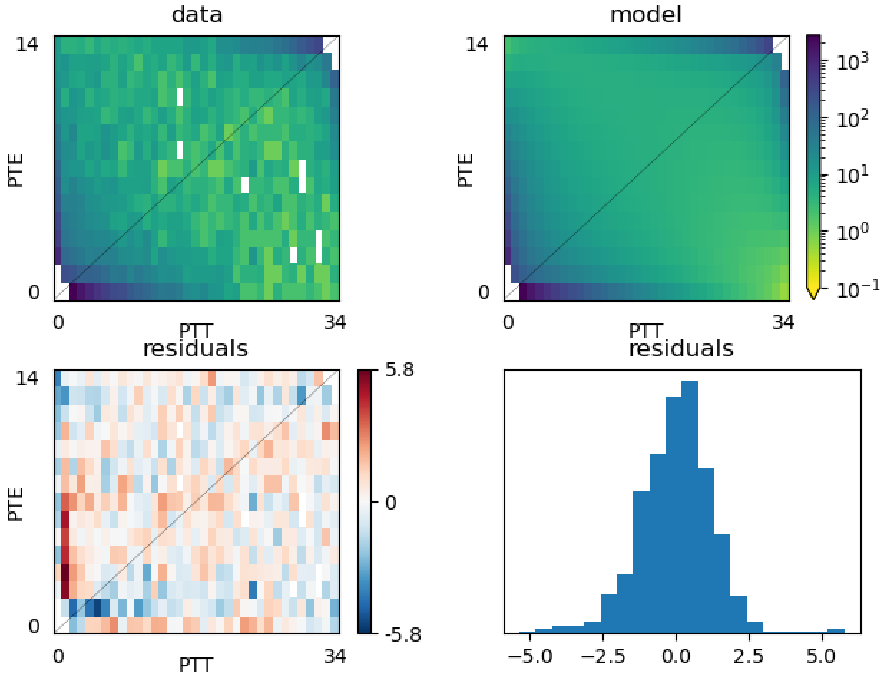


Fig. S74: Model fit to the joint AFS using synonymous SNPs from PTE and PTT. The upper left panel shows the observed joint AFS, the upper right panel shows the inferred AFS based on the maximum likelihood estimates in Table S89, and the remaining panels show the residuals between the observed and model AFS.

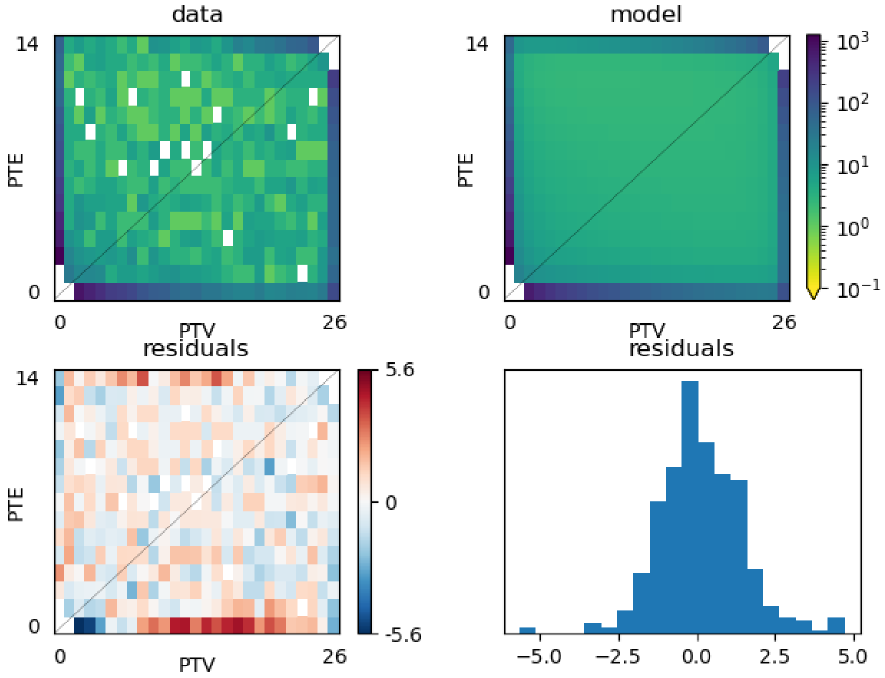


Fig. S75: Model fit to the joint AFS using synonymous SNPs from PTE and PTV. The upper left panel shows the observed joint AFS, the upper right panel shows the inferred AFS based on the maximum likelihood estimates in Table S89, and the remaining panels show the residuals between the observed and model AFS.

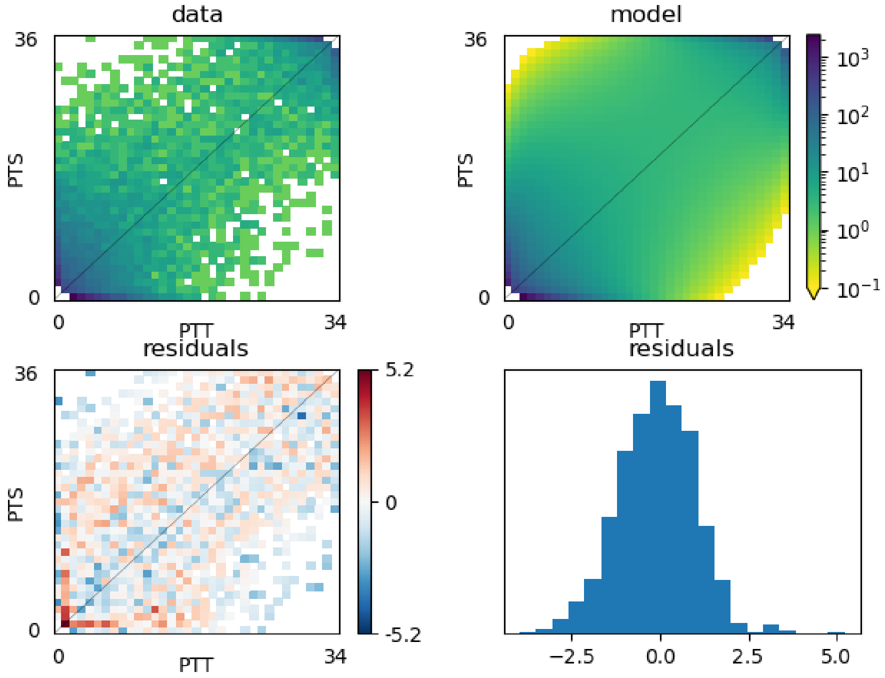


Fig. S76: Model fit to the joint AFS using synonymous SNPs from PTS and PTT. The upper left panel shows the observed joint AFS, the upper right panel shows the inferred AFS based on the maximum likelihood estimates in Table S89, and the remaining panels show the residuals between the observed and model AFS.

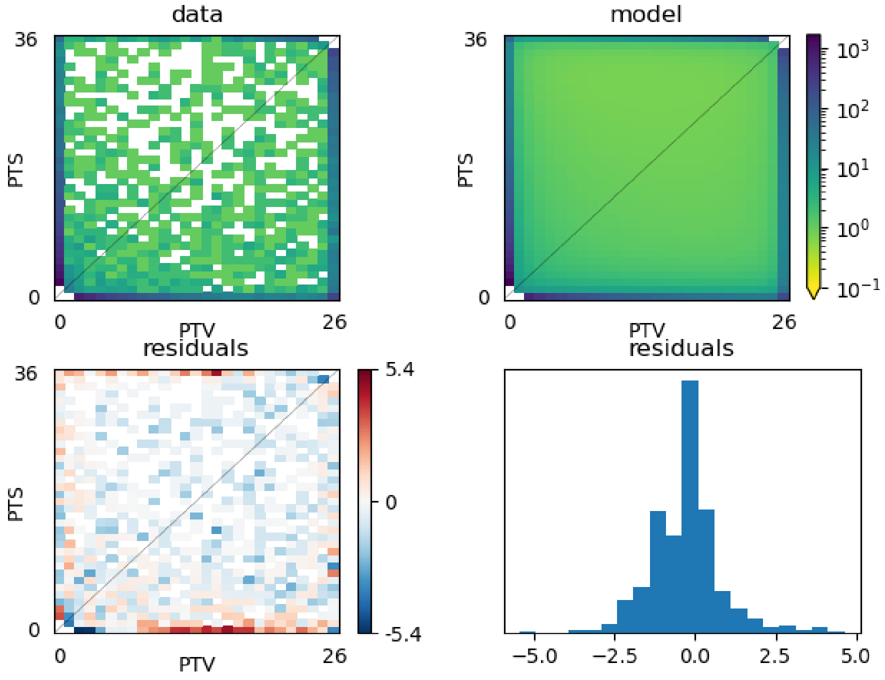


Fig. S77: Model fit to the joint AFS using synonymous SNPs from PTS and PTV. The upper left panel shows the observed joint AFS, the upper right panel shows the inferred AFS based on the maximum likelihood estimates in Table S89, and the remaining panels show the residuals between the observed and model AFS.

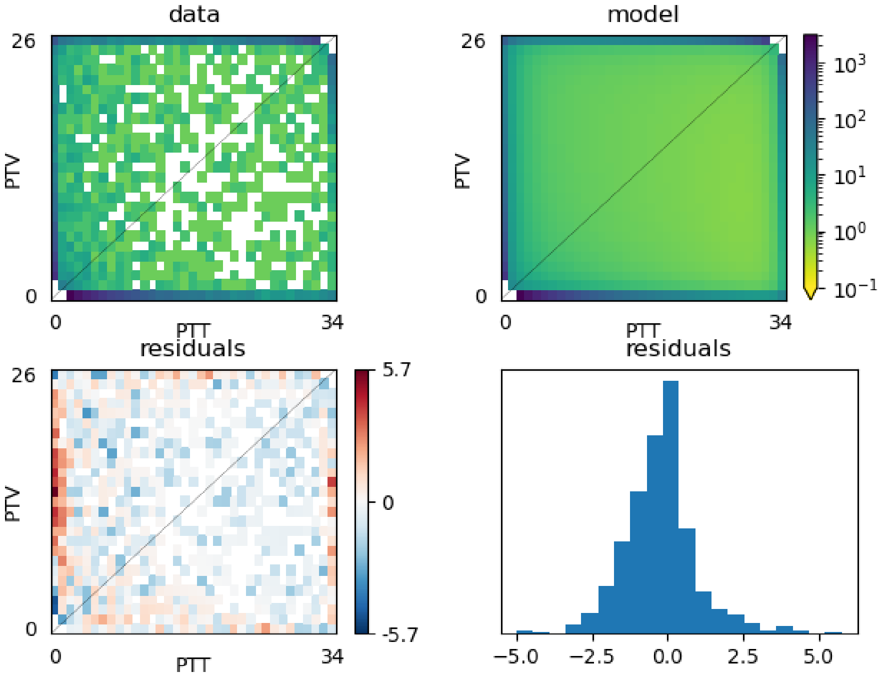


Fig. S78: Model fit to the joint AFS using synonymous SNPs from PTT and PTV. The upper left panel shows the observed joint AFS, the upper right panel shows the inferred AFS based on the maximum likelihood estimates in Table S89, and the remaining panels show the residuals between the observed and model AFS.

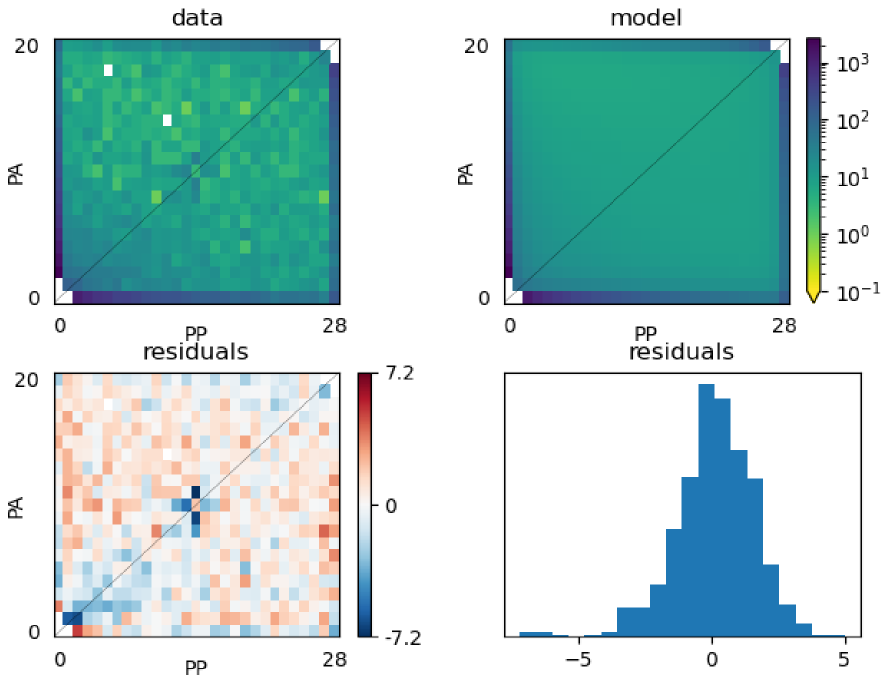


Fig. S79: Model fit to the joint AFS using synonymous SNPs from PA and PP. The upper left panel shows the observed joint AFS, the upper right panel shows the inferred AFS based on the maximum likelihood estimates in Table S89, and the remaining panels show the residuals between the observed and model AFS.

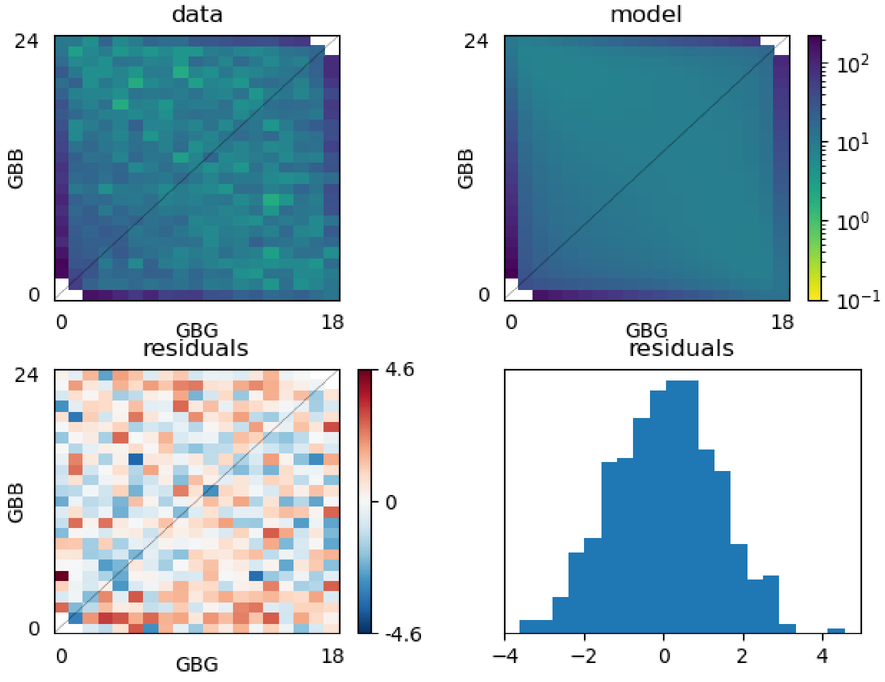


Fig. S80: Model fit to the joint AFS using synonymous SNPs from GBB and GBG. The upper left panel shows the observed joint AFS, the upper right panel shows the inferred AFS based on the maximum likelihood estimates in Table S89, and the remaining panels show the residuals between the observed and model AFS.

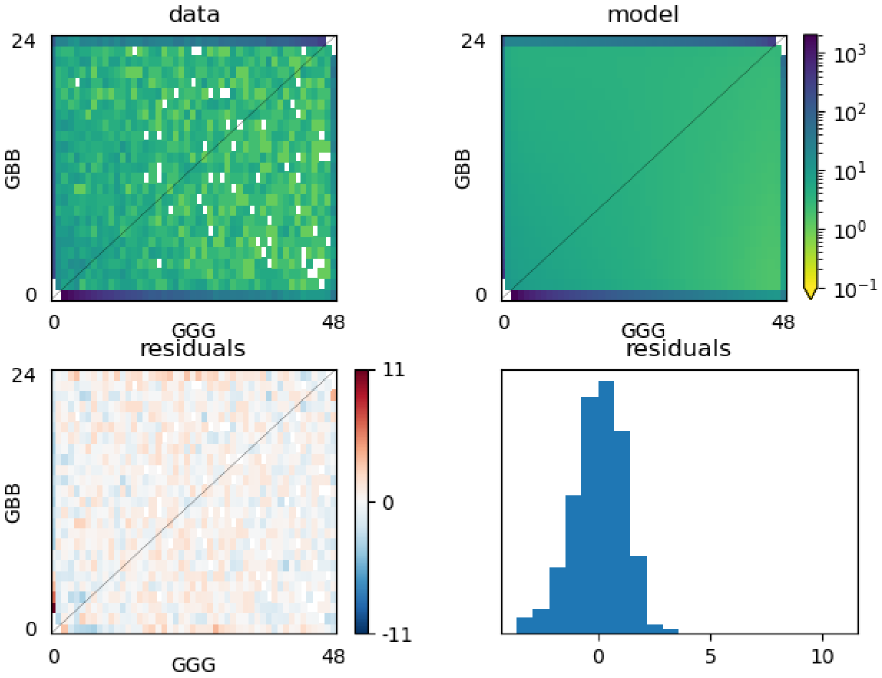


Fig. S81: Model fit to the joint AFS using synonymous SNPs from GBB and GBG. The upper left panel shows the observed joint AFS, the upper right panel shows the inferred AFS based on the maximum likelihood estimates in Table S89, and the remaining panels show the residuals between the observed and model AFS.

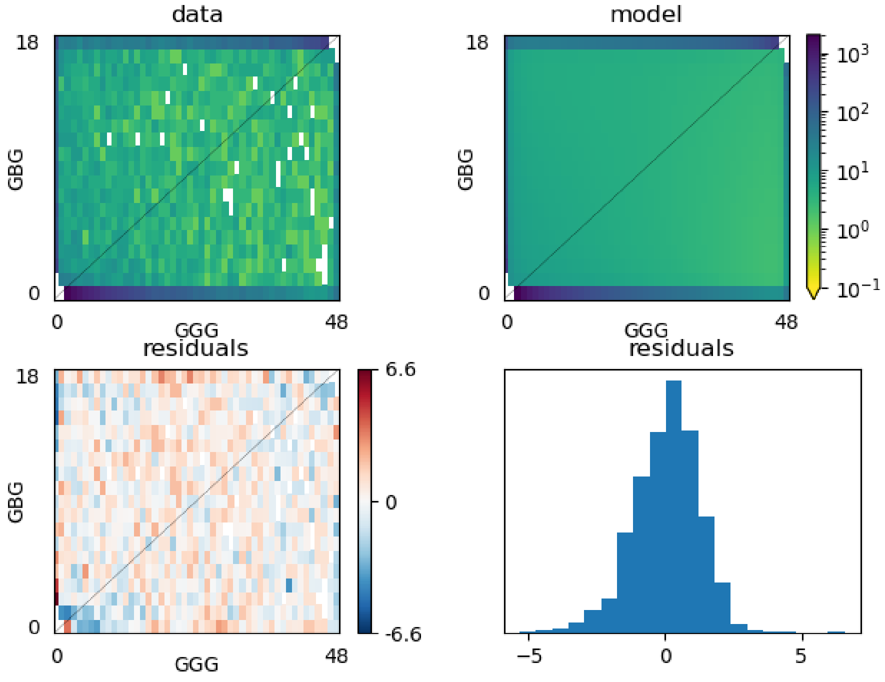


Fig. S82: Model fit to the joint AFS using synonymous SNPs from GBB and GBG. The upper left panel shows the observed joint AFS, the upper right panel shows the inferred AFS based on the maximum likelihood estimates in Table S89, and the remaining panels show the residuals between the observed and model AFS.

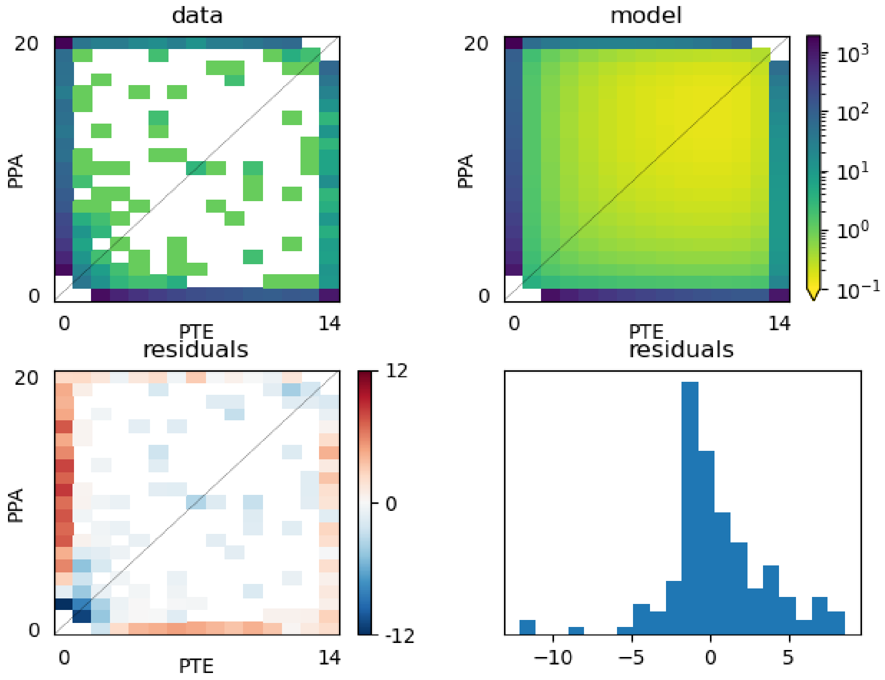


Fig. S83: Model fit to the joint AFS using non-synonymous SNPs from PPA and PTE. The upper left panel shows the observed joint AFS, the upper right panel shows the inferred AFS based on the maximum likelihood estimates in Fig. 3 and Table S89, and the remaining panels show the residuals between the observed and model AFS.

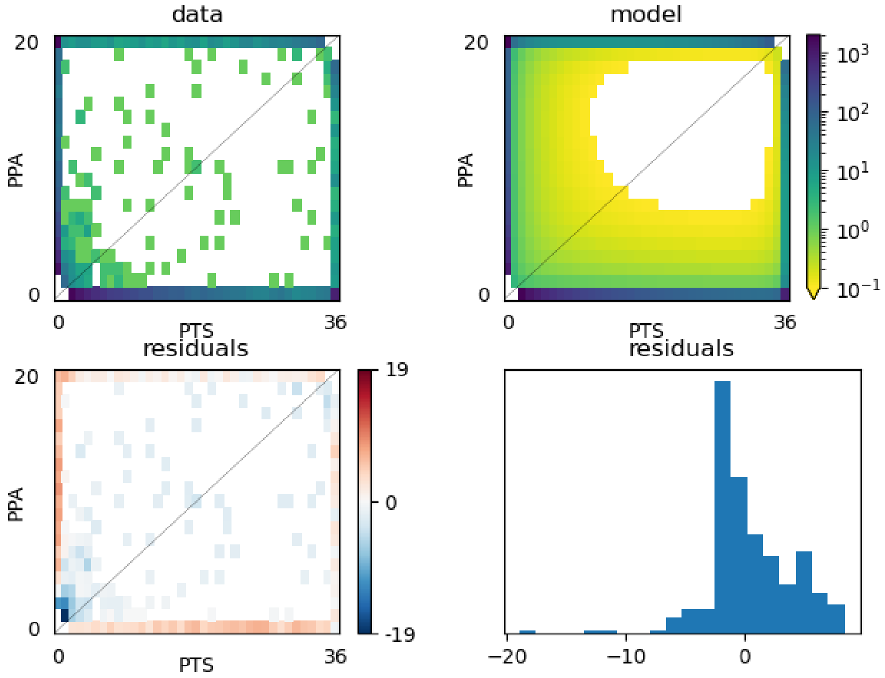


Fig. S84: Model fit to the joint AFS using non-synonymous SNPs from PPA and PTS. The upper left panel shows the observed joint AFS, the upper right panel shows the inferred AFS based on the maximum likelihood estimates in Fig. 3 and Table S89, and the remaining panels show the residuals between the observed and model AFS.

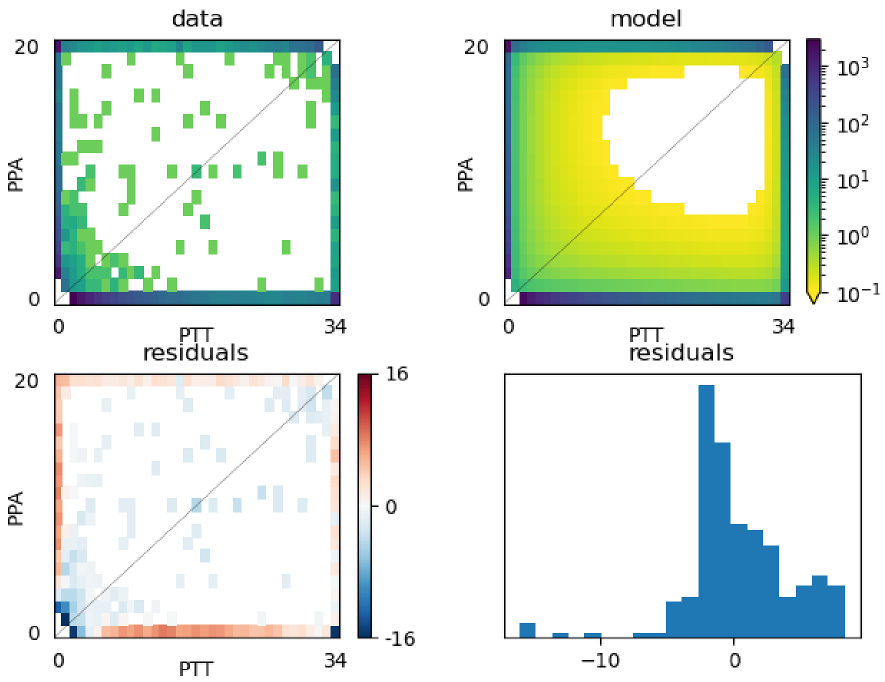


Fig. S85: Model fit to the joint AFS using non-synonymous SNPs from PPA and PTT. The upper left panel shows the observed joint AFS, the upper right panel shows the inferred AFS based on the maximum likelihood estimates in Fig. 3 and Table S89, and the remaining panels show the residuals between the observed and model AFS.

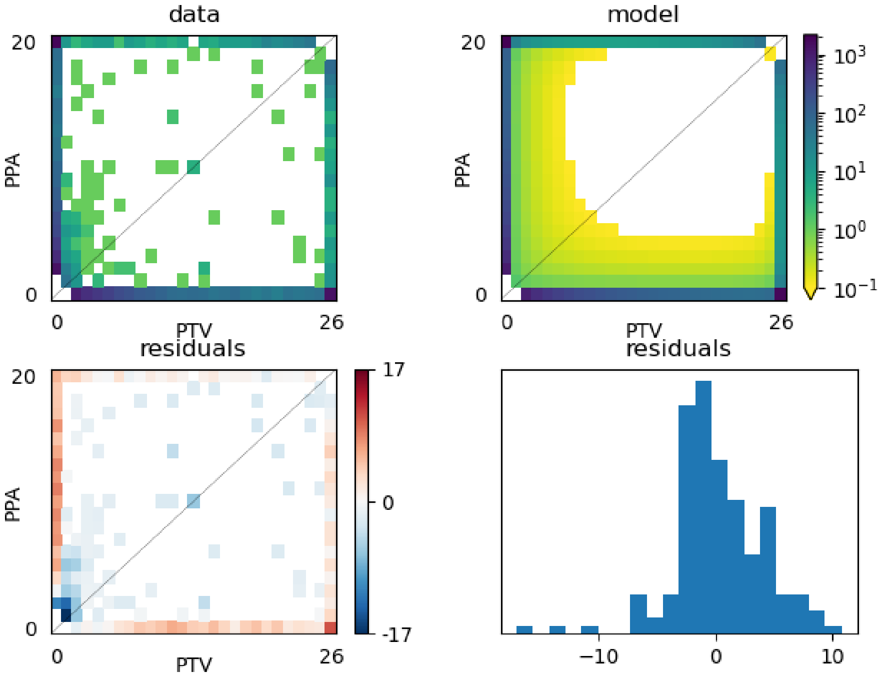


Fig. S86: Model fit to the joint AFS using non-synonymous SNPs from PPA and PTV. The upper left panel shows the observed joint AFS, the upper right panel shows the inferred AFS based on the maximum likelihood estimates in Fig. 3 and Table S89, and the remaining panels show the residuals between the observed and model AFS.

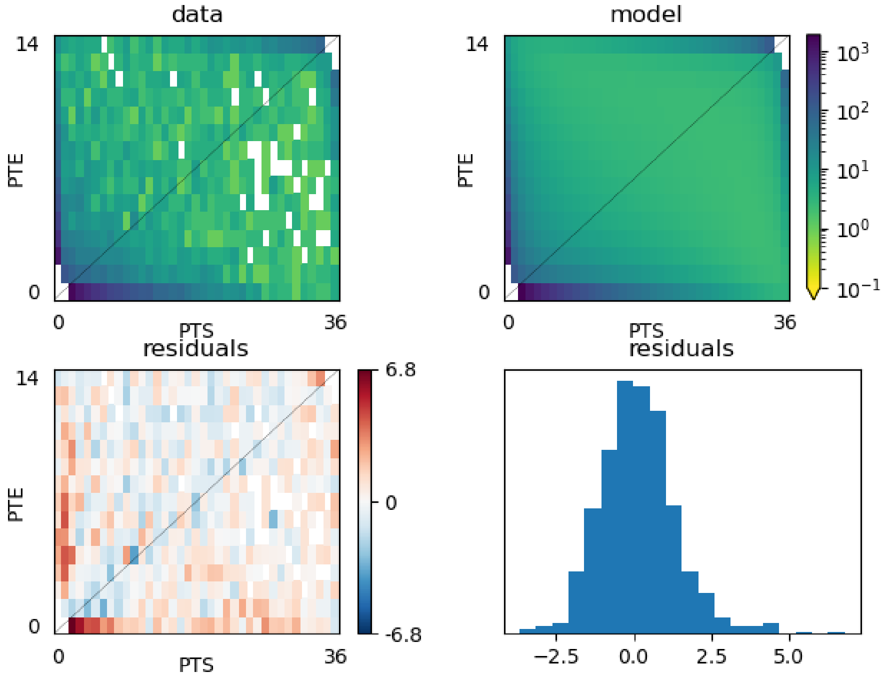


Fig. S87: Model fit to the joint AFS using non-synonymous SNPs from PTE and PTS. The upper left panel shows the observed joint AFS, the upper right panel shows the inferred AFS based on the maximum likelihood estimates in Fig. 3 and Table S89, and the remaining panels show the residuals between the observed and model AFS.

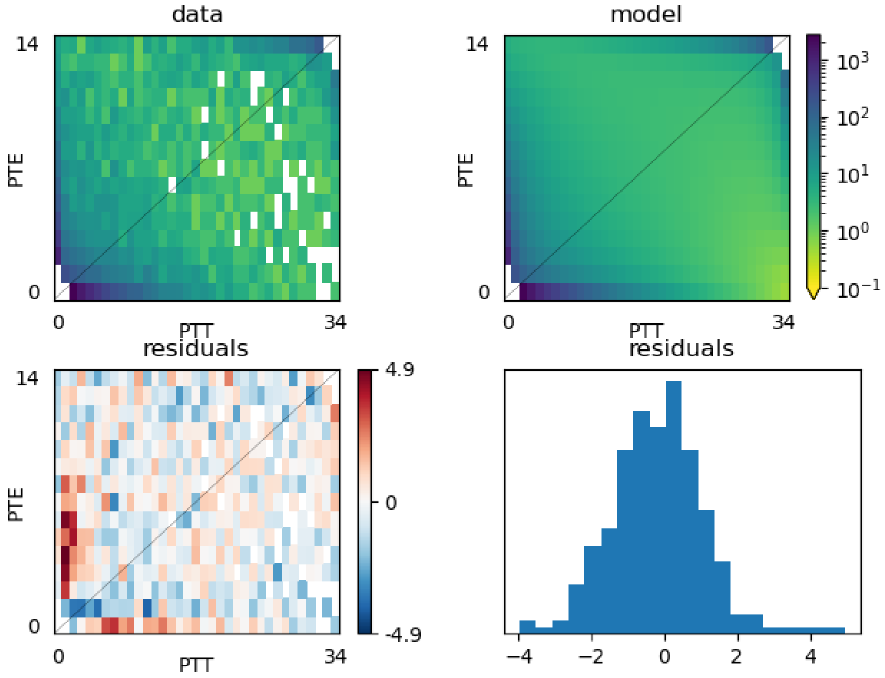


Fig. S88: Model fit to the joint AFS using non-synonymous SNPs from PTE and PTT. The upper left panel shows the observed joint AFS, the upper right panel shows the inferred AFS based on the maximum likelihood estimates in Fig. 3 and Table S89, and the remaining panels show the residuals between the observed and model AFS.

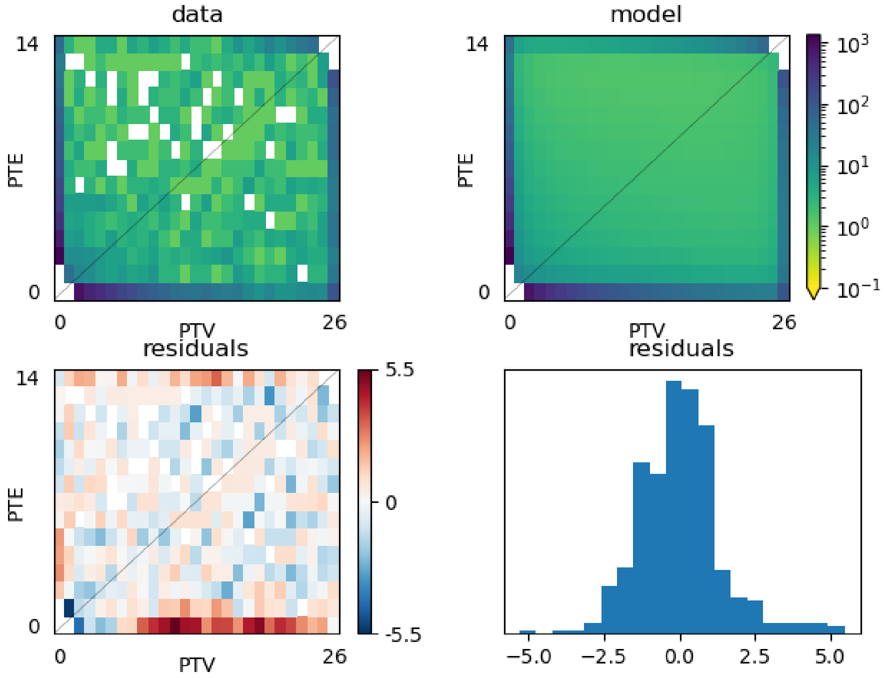


Fig. S89: Model fit to the joint AFS using non-synonymous SNPs from PTE and PTV. The upper left panel shows the observed joint AFS, the upper right panel shows the inferred AFS based on the maximum likelihood estimates in Fig. 3 and Table S89, and the remaining panels show the residuals between the observed and model AFS.

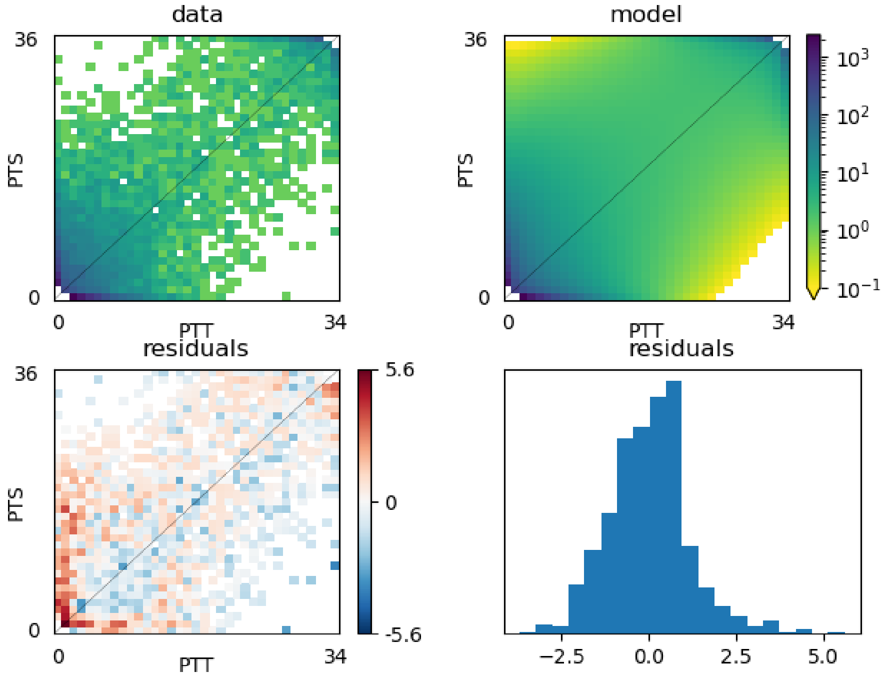


Fig. S90: Model fit to the joint AFS using non-synonymous SNPs from PTS and PTT. The upper left panel shows the observed joint AFS, the upper right panel shows the inferred AFS based on the maximum likelihood estimates in Fig. 3 and Table S89, and the remaining panels show the residuals between the observed and model AFS.

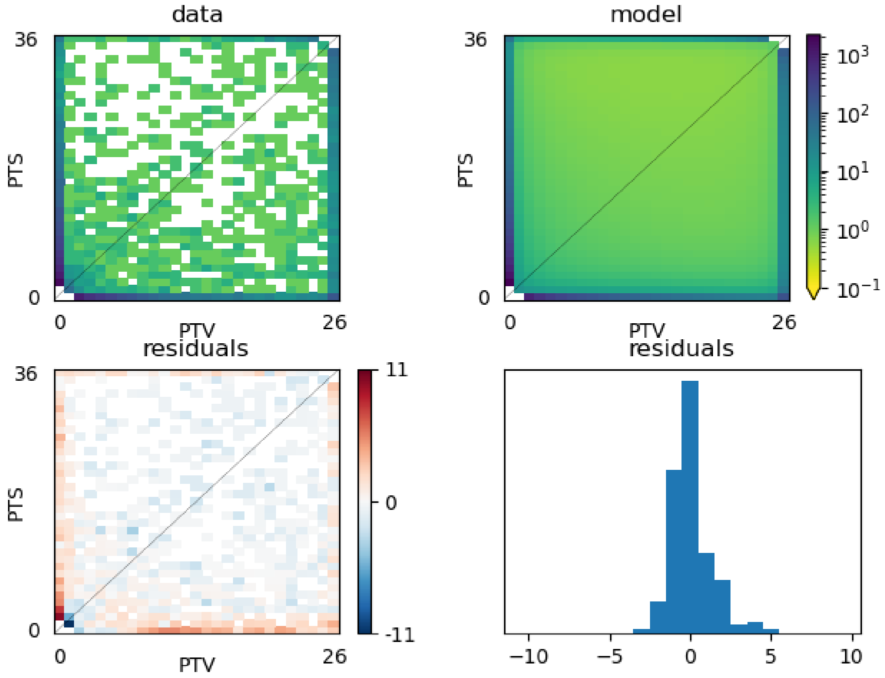


Fig. S91: Model fit to the joint AFS using non-synonymous SNPs from PTS and PTV. The upper left panel shows the observed joint AFS, the upper right panel shows the inferred AFS based on the maximum likelihood estimates in Fig. 3 and Table S89, and the remaining panels show the residuals between the observed and model AFS.

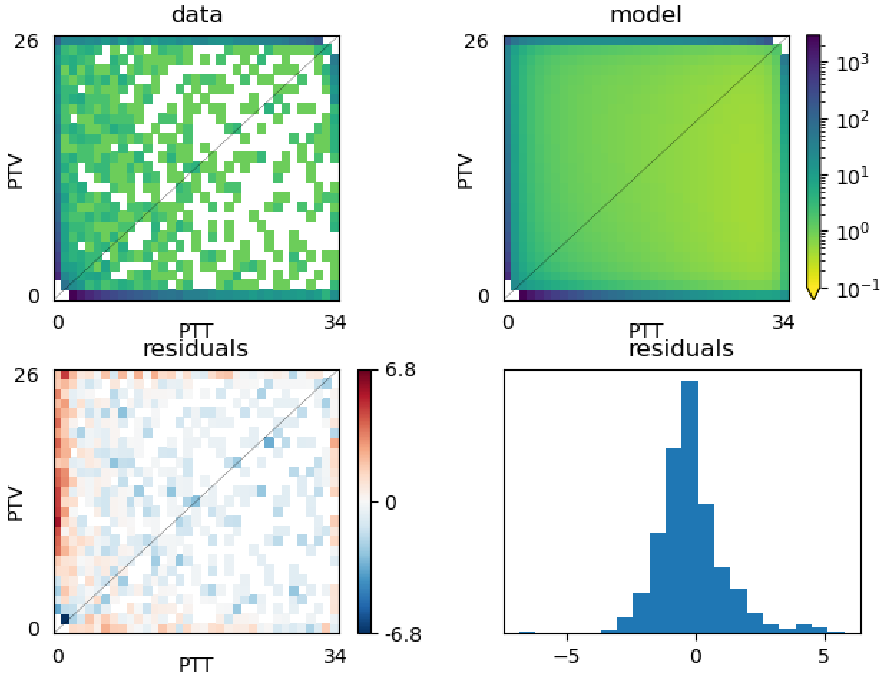


Fig. S92: Model fit to the joint AFS using non-synonymous SNPs from PTT and PTV. The upper left panel shows the observed joint AFS, the upper right panel shows the inferred AFS based on the maximum likelihood estimates in Fig. 3 and Table S89, and the remaining panels show the residuals between the observed and model AFS.

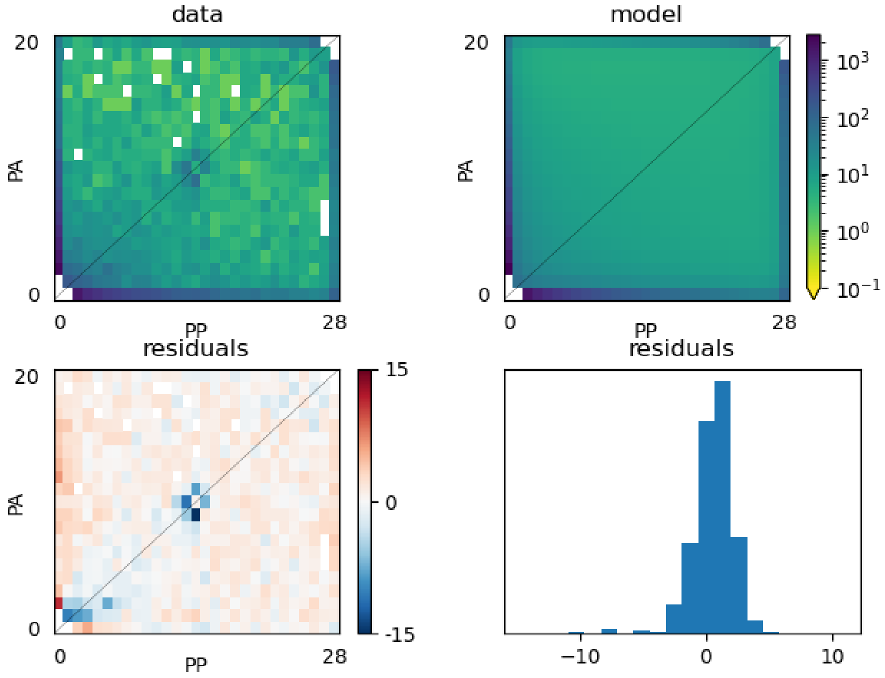


Fig. S93: Model fit to the joint AFS using non-synonymous SNPs from PA and PP. The upper left panel shows the observed joint AFS, the upper right panel shows the inferred AFS based on the maximum likelihood estimates in Fig. 3 and Table S89, and the remaining panels show the residuals between the observed and model AFS.

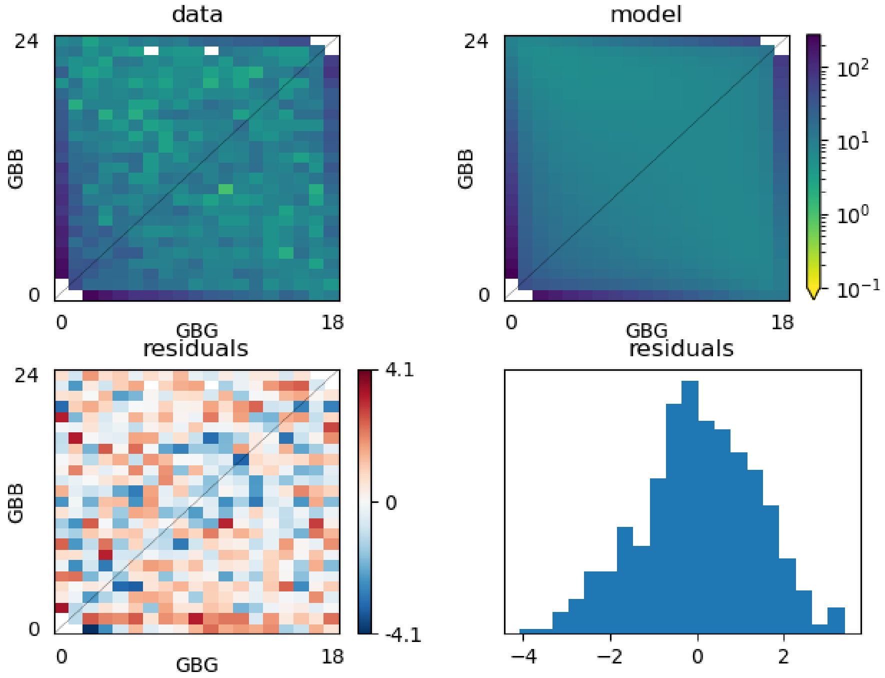


Fig. S94: Model fit to the joint AFS using non-synonymous SNPs from GBB and GBG. The upper left panel shows the observed joint AFS, the upper right panel shows the inferred AFS based on the maximum likelihood estimates in Fig. 3 and Table S89, and the remaining panels show the residuals between the observed and model AFS.

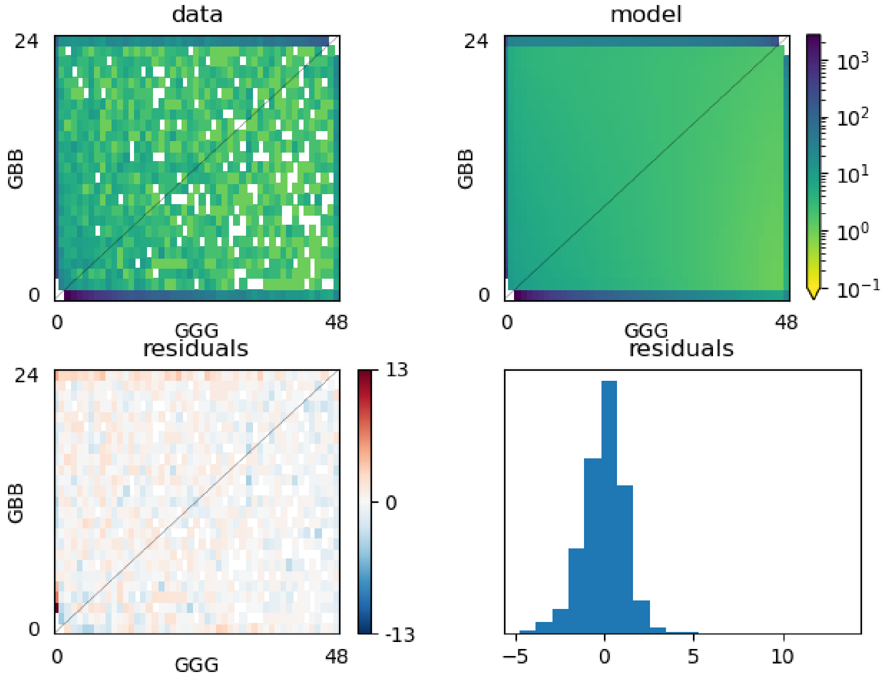


Fig. S95: Model fit to the joint AFS using non-synonymous SNPs from GBB and GBG. The upper left panel shows the observed joint AFS, the upper right panel shows the inferred AFS based on the maximum likelihood estimates in Fig. 3 and Table S89, and the remaining panels show the residuals between the observed and model AFS.

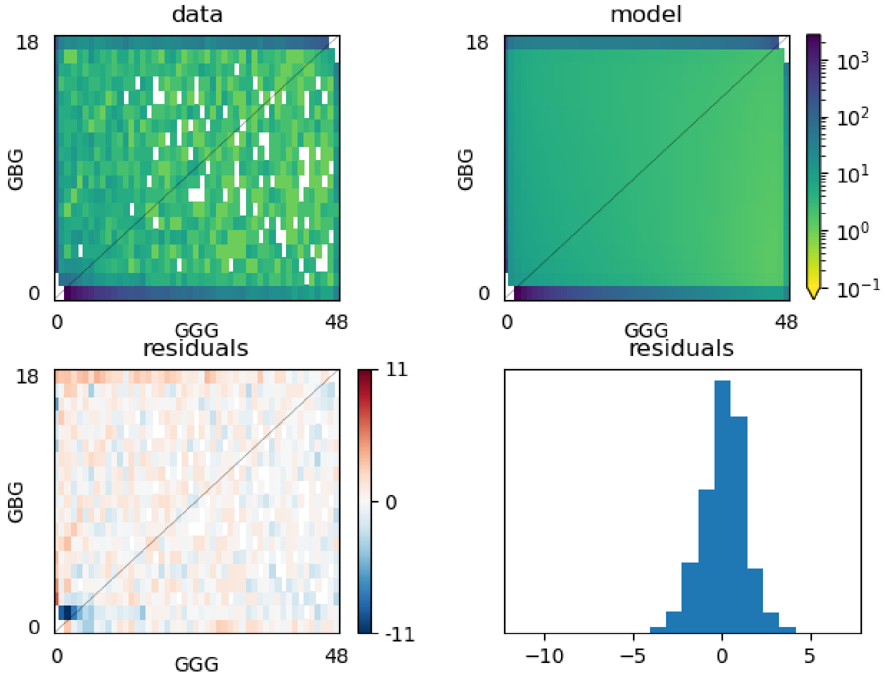


Fig. S96: Model fit to the joint AFS using non-synonymous SNPs from GBB and GBG. The upper left panel shows the observed joint AFS, the upper right panel shows the inferred AFS based on the maximum likelihood estimates in Fig. 3 and Table S89, and the remaining panels show the residuals between the observed and model AFS.
